# Multiple Routes to Metacognitive Judgments of Working Memory in the Macaque Prefrontal Cortex

**DOI:** 10.1101/2025.07.15.664842

**Authors:** Chuan Ning, Guobin Fu, Yan-Yu Zhang, Florent Meyniel, Liping Wang

**Author notes:** **Corresponding author:** Liping Wang, Address: Institute of Neuroscience, Chinese Academy of Sciences, 320 YueYang Road, Shanghai 200031, China. C.N., G.F., and Y.Y.Z. contributed equally to this work.

## Abstract

The ability to evaluate one’s own memory is known as metamemory. Whether metamemory is inherent to memory strength or requires additional computation in the brain remains largely unknown. We investigated the metacognitive mechanism of working memory (WM) using two-photon calcium imaging in the prefrontal cortex of macaque monkeys, who were trained to memorize spatial sequences of varying difficulties. In some trials, after viewing the sequence, monkeys could opt out of retrieval for a smaller reward, reflecting their confidence in WM (meta-WM). We discovered that PFC neurons encoded WM strength by jointly representing the remembered locations through population coding and their associated uncertainties. This WM strength faithfully predicted the monkeys’ recall performance and opt-out decisions. In addition to memory strength, other factors— trial history and arousal—encoded in baseline activity predicted opt-out decisions, serving as cues for meta-WM. We identified a code of meta-WM itself that integrated WM strength and these cues. Importantly, WM strength, cues, and meta-WM were represented in different subspaces within the same PFC population. The dynamics and geometry of PFC activity implement metacognitive computations, integrating WM strength with cues into a meta-WM signal that guides behavior.

## Introduction

Working memory (WM) refers to the short-term maintenance and control of information necessary for future actions^1^. However, WM is significantly limited, typically holding only three or four items at a time, and its precision decreases as the number of items increases^2,3^. Imagine you must remember to buy wine on your way home from work, email a friend later in the evening, and take a pill at 9 p.m. Given our self-awareness of WM capacity, we often set external reminders to aid our memory, such as scheduling alarm clocks on our smartphones. This process relies on metacognitive judgments about our WM limitations^4^. This meta-level WM process is referred to as “meta-WM”^5,6,7^. We routinely use meta-WM to improve our decisions, for example, by offloading our memory to the external environment^8^. Convergent findings have emphasized the importance of the prefrontal cortex (PFC) in two aspects of WM, including the maintenance of WM content^9–11^ and accurate metacognitive judgments about memory or perception in humans^12–14^, monkeys^15,16^, and rodents^17,18^. The connectivity between the prefrontal cortex and other brain areas is vital for metamemory and perceptual metacognition^16,19^, with different parts of the PFC used for these two domains^19,20^.

While metacognition research pertains to personal beliefs and knowledge about self-performance, the neuroscience of confidence has been mostly restricted to studying the neural representation of uncertainty within sensory or motor systems^21^. Thus, a key challenge lies in understanding how the brain integrates the neural representation of uncertainty and performance judgment into a single, unified framework. Furthermore, a long-standing debate about metacognition concerns whether meta-WM is inherent to WM strength or requires additional computation^21^. On one hand, judgments of higher memory strength are generally associated with higher accuracy in memory tasks^7,22^. In perception decision-making, neurons in the parietal cortex jointly represent the decision of direction and the degree of certainty underlying the decision to opt out^23,24^. On the other hand, metamemory does not always reflect accuracy but rather favors the use of (sometimes misleading) cues, such as fluency^25^, processing experience, prior beliefs^26,27^, and interoceptive signals^28^.

Therefore, despite previous studies on confidence and metacognition, the neural bases of WM and meta-WM at the single- or population-level in the PFC remain largely unknown. Furthermore, there is still a lack of a complete conceptual framework to explain how the brain combines WM strength and cues into a meta-WM signal for further action. To investigate the neural representations and computations of metacognition about WM, in this study, we aimed to test 1) whether WM, cues, and metacognitive judgment (meta-WM) are represented by the same neurons in the PFC, and 2) whether and how PFC neurons integrate the current strength of WM and cues to inform meta-WM to control behavior.

## Results

### Paradigm and behavior

We trained two macaque monkeys (D and Z) to learn a delayed-sequence reproduction meta-WM task (Fig. 1A). Task difficulty was controlled by varying both the sequence length and spatial location combination. The monkeys had to maintain fixation on the cross at the center of the screen as the baseline to initiate a trial. On each trial, spatial sequences with a length of 1, 2, 3, or 4 items were visually presented during the sample period. Each item was randomly drawn (without replacement) from one of the six spatial locations of a hexagon and presented sequentially. The monkeys were required to memorize the sequence for a short delay with fixation and report it during the response period (see Methods). In a random 65% of the trials (“choice” condition), after the delay period, the monkeys were allowed to freely choose to perform (“memory” condition) or opt out of retrieval (“offload” condition, smaller reward). In the other trials (“forced-to-test” condition), the monkeys were forced to perform the retrieval task. Monkey did not know the condition before the decision period. This task design of prospective meta-WM encourages monkeys to estimate their confidence about their WM on each trial during the delay period, and to opt out when they are less confident about their WM in the “choice” condition.

**Fig. 1.**
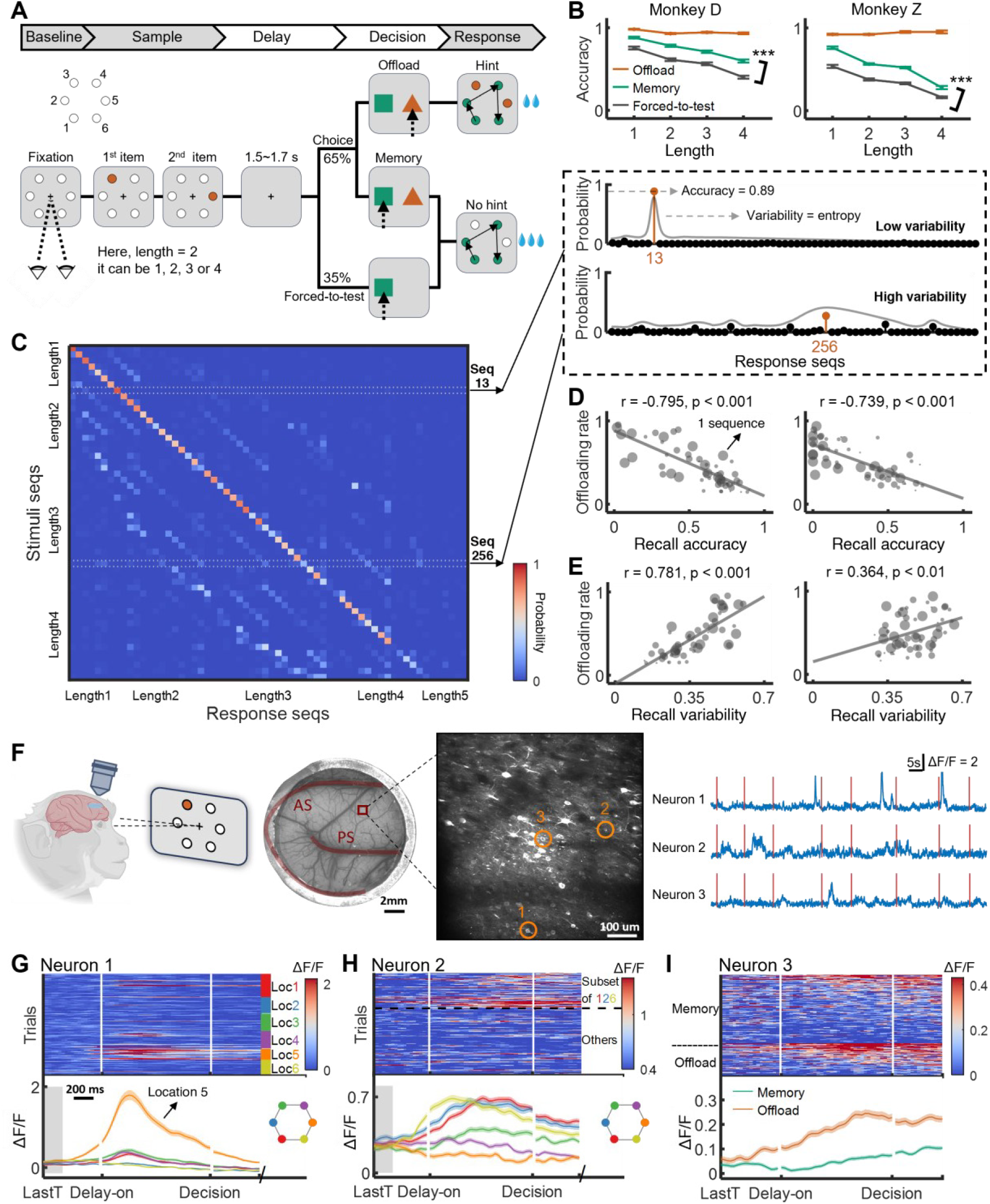
Task design, behavior, and two-photon calcium imaging. (**A**) Schematic of a delayed-sequence reproduction meta-WM task. Monkeys initiate a trial by fixating at the central cross for 1.1s (denoted as “baseline” period). On each trial, a sequence containing 1, 2, 3, or 4 items is presented sequentially in a clockwise order (from location 1 to 6) on the display (“sample” period). After a 1.5-1.7s delay period, in 65% of (choice) trials, the monkeys can freely choose either to perform the retrieval task (“memory” condition, with a large reward if correct, green square) or opt out of retrieval (“offload” condition, with a small reward if correct, red triangle). In the remaining 35% of trials, the monkeys are forced to perform the task (“forced-to-test” condition). The chosen options appear randomly on either the left or right side of the screen across trials. Monkeys were required to saccade to the items that did not appear during the sample period in a clockwise order, thereby minimizing the possibility of motor preparation for the stimuli during the delay. (**B**) Behavioral accuracy of the three conditions for different sequence lengths in both monkeys. Accuracy denotes the fraction of correct responses in each condition. Error bars represent SEMs across sessions. ***: p < 0.001 (two-sample t-test for each length). (**C**) Calculation of recall variability for monkey D. The stimuli-to-response matrix in the forced-to-test condition (left) includes both correct trials (diagonal) and error trials (non-diagonal). Each element of the matrix denotes the probability for monkey D to execute one particular sequence given a stimulus sequence. Recall variability is defined as the normalized entropy of response sequence probabilities given a stimulus sequence. Of the two example sequences [1-3] and [2-5-6] shown here (inset to the upper right), for monkey D, the former has low variability (concentrated probabilities), and the latter has high variability (widely distributed probabilities). (**D**) Pearson correlations between the behavioral accuracy (from the forced-to-test condition) and the offloading rate (from the choice condition) across all sequences for both monkeys. Offloading rate denotes the probability of choosing offload in choice trials. Each dot represents a sequence, and the dot size represents sequence lengths (larger = longer). 1 sequence: one sequence. (**E**) Pearson correlations between the recall variability (computed as in (C), from forced-to-test condition) and the offloading rate (from choice condition) across all sequences for both monkeys. (**F**) Two-photon calcium imaging setup. Each monkey has an image recording window over the LPFC with both the arcuate sulcus (AS) and the principal sulcus (PS) in view. Normalized calcium traces for three example neurons (orange circles) in one representative FOV (red square) are shown in the rightmost panel. Vertical red lines indicate the start of each trial. ΔF/F: normalized fluorescent intensity. (**G-I**), Three example neurons in (F). Each shows tuning to a single location (G); multiple locations ((H); [1], [2], [6]; and sequences only containing those locations, [1-2], [1-6], [2-6], [1-2-6]); or meta-WM decisions (I), respectively. Each panel contains both stacked single trial ΔF/F traces (top) and averaged traces (bottom). “LastT” labels on horizontal axes mark the onset times of the last target stimulus. Shaded areas around average traces represent SEMs across trials.

We examined the relationship between the probability of choosing the offload option (offloading rate), task performance (accuracy of sequence recall), and WM difficulty (sequence length and location combination) to test whether the offloading rate could reflect monkeys’ judgment of their own WM. WM performance declined as the sequence length increased. Crucially, the monkeys performed significantly better when they decided to perform the memory task than when they were forced to do it. This is true regardless of the sequence length (Fig. 1B, ps < 0.001 for each monkey) and the sequence type (Fig. S1A), suggesting that the meta-WM was not determined solely by the sequence locations and length but also by trial-to-trial variability that affects WM.

Figure 1C shows the proportion of response sequences for each stimulus sequence (also see Fig. S1B). We defined for each sequence, recall accuracy as the probability of correct retrieval, and recall variability as the entropy of the distributions of retrievals across trials (see two examples in Fig. 1C upper right). We found that, across sequences, the offloading rate measured in the choice condition was negatively correlated with recall accuracy and positively correlated with recall variability measured in the forced-to-test condition (Fig. 1D and 1E, ps < 0.001 for both monkeys). Thus, in this paradigm, the offloading decision reflects the monkeys’ accurate assessment of their own WM.

### Recordings and single-neuron responses

During the delay, monkeys had to maintain WM and meta-WM representations, which respectively correspond to a (first-order) representation of the remembered locations and a (second-order) representation of the strength of the first-order representation. To search for the neural representations of the two aspects of WM and their dynamics, we injected TET-Off GCaMP6f virus into the LPFCs of the two monkeys to enable two-photon calcium imaging (Fig. 1F and Methods) and recorded more than 15,000 single neurons [monkey D, 4,758 neurons from 12 fields of view (FOVs); monkey Z, 10,960 neurons from 16 FOVs (Fig. S2A)]. We first focused on neural activity during the late delay period (1.1 seconds before decision). Many neurons showed selectivity for single-or multiple-location (22.3% for monkey D, 10.2% for monkey Z, see examples in Fig. 1G-H). Meanwhile, some neurons exhibited selectivity for meta-WM decision before the decision was made (see example in Fig. 1I). The distributions of cell selectivity for location and meta-WM decision are shown in Figure S2B and relatively consistent across the two monkeys (Table S1).

### Representation of WM and memory strength in the LPFC neural population

Next, we examined whether the PFC population neurons represent the WM of sequential locations and their uncertainties (i.e., WM strength). We decoded single-trial WM representations of locations from population neural activity during the delay period. For each FOV, we trained six linear support vector machines (SVMs) to decode the probability of each location (one SVM per location). The decoded probability of a sequence in WM was defined as a joint probability of multiple locations (since the locations were always presented clockwise, we did not decode their ordinal positions). In addition, the uncertainty of the sequence representation in WM was measured as the entropy of the decoder’s probability distributions across all sequences. We inverted and normalized the entropy as a trial-by-trial measure of WM strength (Fig. 2A, see Methods).

**Fig. 2.**
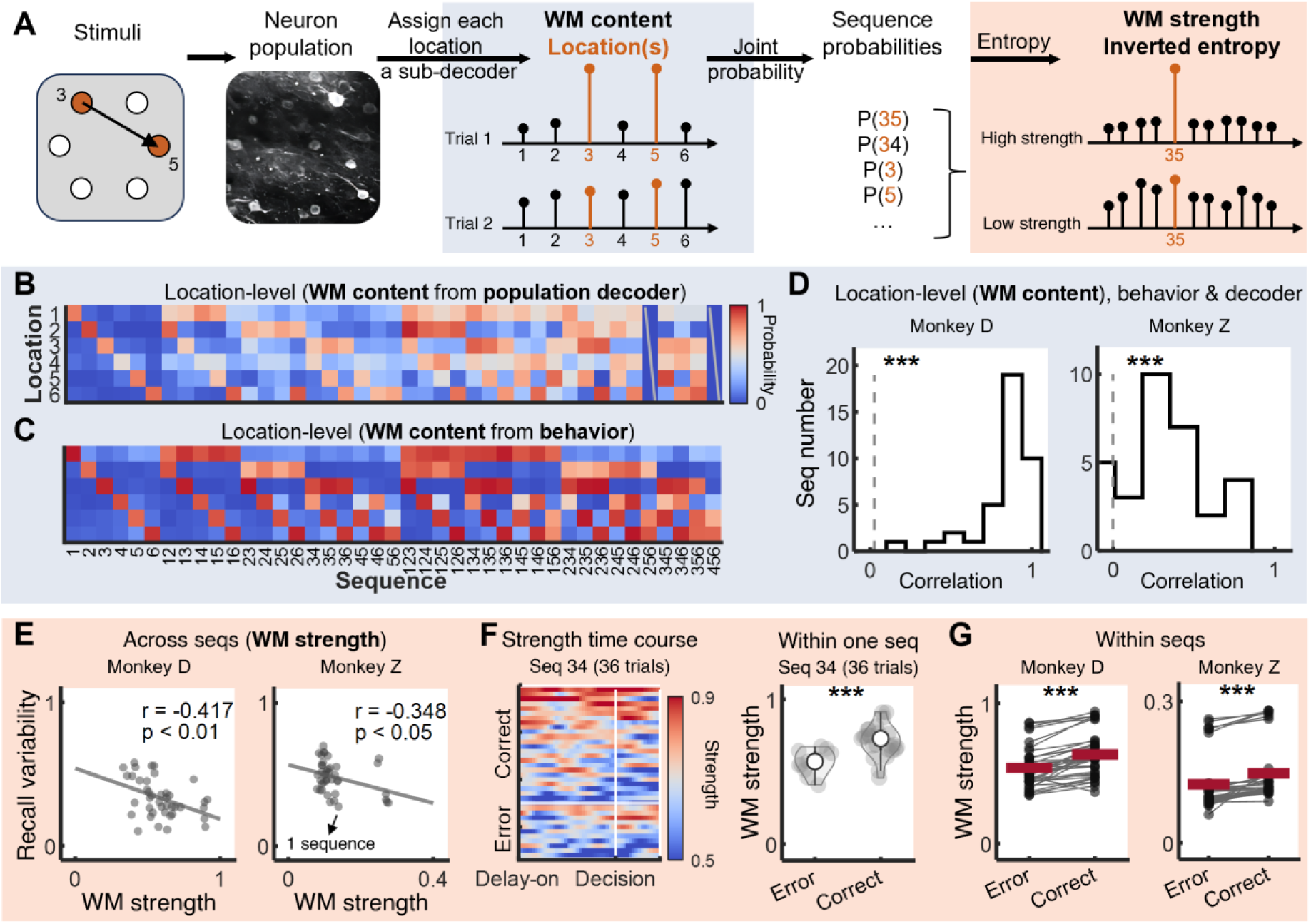
Joint representation of WM content and its strength in the LPFC neural population. (**A**) Schematic illustration of a population decoder that represents both WM content (locations) and WM strength. We used the decoded location probability distribution to compute the decoded probability of each sequence (e.g., [3-5]) with multiple items. This was achieved through a joint probability approach, which involved calculating the product of all possible locations’ probabilities. Neural decoder-based WM strength was defined and computed as the inverted normalized entropy of the distribution of decoded probabilities of all sequences ([3-5], [3-4], [3], [5], etc.; see Methods subsection “Neural decoder-based WM strength”). (**B**) Location probability distributions based on an example neural population decoder (monkey D, FOV 8), representing, for each sequence in the forced-to-test condition, the decoded probabilities of each location. The missing columns correspond to sequences that lacked correct trials during the session. For similar analysis using all data across FOVs, see Figs. S3 and S5A. (**C**) Behavioral location probability distributions (monkey D), representing, for each sequence in the forced-to-test condition, the response probabilities of each location based on task performance. (**D**) Histogram of Pearson correlation between location probability distributions, given a same stimulus sequence, of neural population decoder (B) and behavior (C) (***: p < 0.001, two-tailed t-test against correlation value by chance). Gray dashed line, chance level. (**E**) Pearson correlations between the behavioral recall variability and the neural WM strength (both from forced-to-test conditions, including correct and error trials) across all sequences for both monkeys (monkey D FOV 8 & monkey Z FOV 9 are shown here; see Fig. S5B for other FOVs). Each dot represents one sequence. (**F**) (Left) Time courses of neural WM strength across all trials within an example sequence [3-4] (sorted by WM strength descending, separately for correct/error trials), involving trials in the forced-to-test and the memory conditions. (Right) Comparison of neural WM strength under correct/error trials within the example sequence [3-4] (***: p < 0.001, two-tailed two-sample t-test). Each dot represents a single trial. (**G**) Within-sequence comparison of neural WM strength between error trials and correct trials of all sequences (***: p < 0.001, two-tailed paired-sample t-test, from monkey D FOV 8 & monkey Z FOV 9), using forced-to-test and memory (correct + error) trials. Each dot represents a sequence and the red lines represent the mean across sequences.

We investigated whether decoding of WM locations from the LPFC neurons reflects the locations reported by monkeys. In the forced-to-test condition, for each sequence, we compared, on average across trials, the decoded locations (Fig. 2B) with behavior (Fig. 2C). In most FOVs across two monkeys, we found strong positive correlations for most sequences (example FOVs in Fig. 2D, all ps < 0.001; other FOVs in Figs. S3 and S5A). Similar results could be found in the choice condition (Fig S4).

We then examined whether the decoded WM strength predicted retrieval performance. Sequences with stronger WM strength across trials resulted in reduced variability of retrievals across trials in both monkeys (Figs. 2E and S5B). Furthermore, WM strength was higher in correct trials than in error trials for most sequences (see the example sequence in Fig. 2F, all sequences for each monkey in Fig. 2G, and Fig. S5C for other FOVs). Thus, the results indicate that the population activity of LPFC neurons jointly represents the remembered locations and their associated uncertainty, and this joint representation predicts the monkeys’ recall performance.

### Meta-WM representations in the LPFC neural population

We next asked whether the meta-WM judgment is also represented in the same neural population. At the single-neuron level, we found a substantial number of neurons selective to the meta-WM decision (whether to offload or not) during the delay period. Figures 3A-B show an example neuron with higher neural responses in the offloading trials than in the memory trials (Fig. 3A) and a significant association with offloading rates across sequences (Fig. 3B). We refer to such neurons as meta-WM neurons (see Methods).

**Fig. 3.**
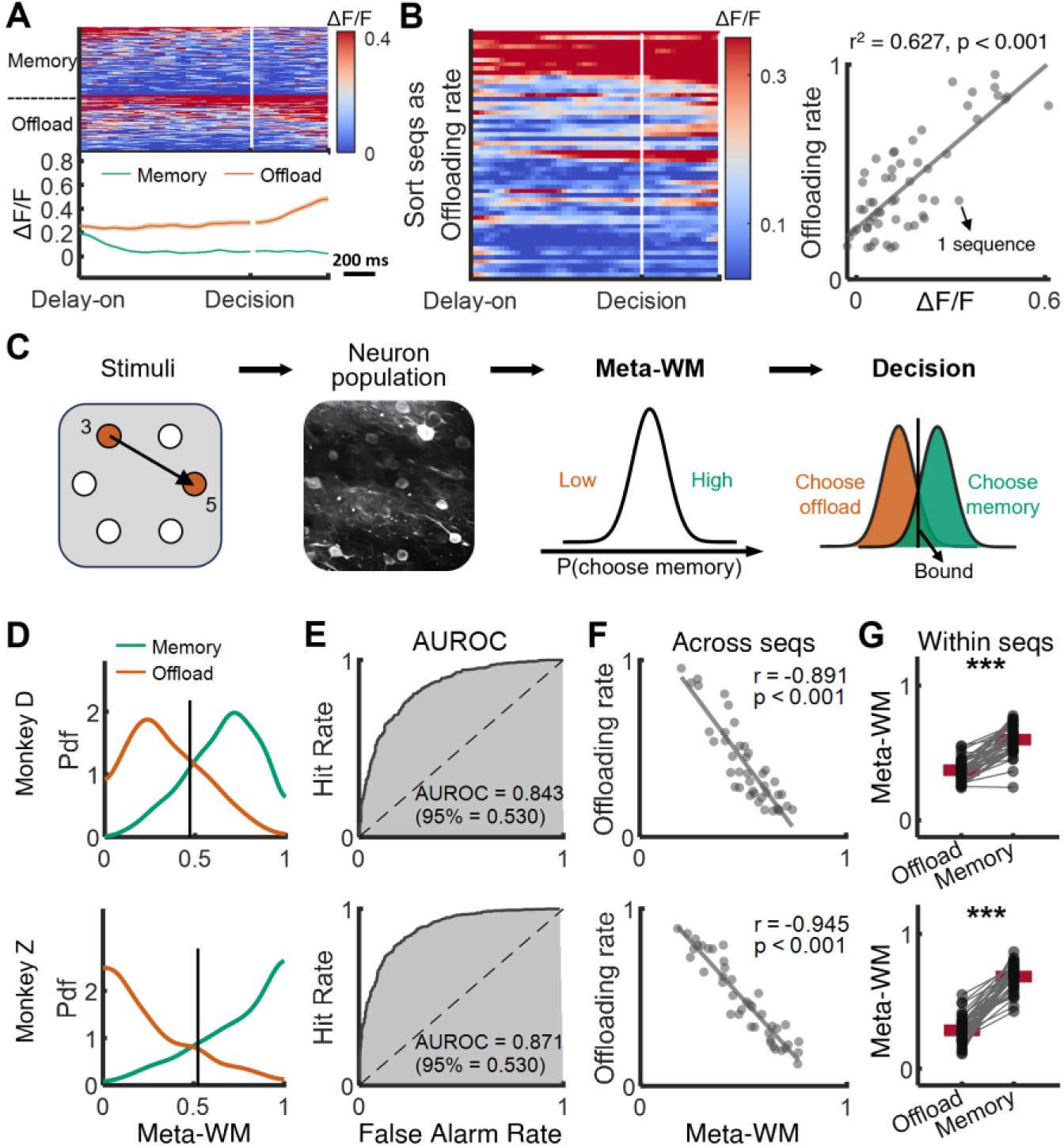
Representation of meta-WM judgements in the LPFC neural population. (**A-B**) An example meta-WM neuron, which is selective to meta-WM decisions as well as offloading rate. (A) Selectivity of meta-WM decisions. (B) (Left) Time courses of activity during the delay period, sorted by the offloading rate associated with each sequence. (Right) Linear regression between offloading rates and the delay-averaged neuron activity. Each dot represents one sequence. (**C**) Schematic illustration of a meta-WM neural decoder, which evaluates meta-WM scores for individual trials by decoding the probability of choosing the memory option in each trial, subsequently making decisions. (**D-G**), Results of the meta-WM decoder from FOV 8 of monkey D and FOV 9 of monkey Z, using choice (memory + offload) trials. (**D**) Probability density functions (Pdfs) of the estimated meta-WM scores for all memory trials and all offload trials for both monkeys. Black solid lines represent decision boundaries of the meta-WM decoder. (**E**) The meta-WM decoder’s performance was evaluated using the area under receiver operating characteristic (AUROC) curve. The “95%” represents 95th percentile of the AUROC computed from trial label-shuffled decoders as control. (**F**) Pearson correlations between the offloading rate and the estimated meta-WM scores across all sequences for both monkeys. Each dot represents a sequence. (**G**) Within-sequence comparison of meta-WM scores between the offload and the memory trials of all sequences (***: p < 0.001, two-tailed paired-sample t-tests). Each dot represents a sequence, and red lines represent the means across sequences.

At the population level, we used linear regression (SVM-based) to train a decoder of meta-WM decisions from single-trial neural responses during the late delay period from each FOV. The decoder returns a *meta-WM score* in favor of performing the memory task (Fig. 3C; see Methods). Note that all decoding results are cross-validated. We discovered that the meta-WM judgment at single trials could be accurately decoded from the delay responses in both monkeys (Figs. 3D-E and S6). The meta-WM score was significantly correlated with the offloading rate across sequences (Fig. 3F ps < 0.001). Importantly, within nearly all sequences, the meta-WM score was larger when monkeys chose to perform the memory task than when they chose to offload (Fig. 3G, ps < 0.001; see Fig. S7 for results from other FOVs). Therefore, the metacognitive judgment of WM (i.e., meta-WM) was also represented in LPFC neurons.

### The relationship between WM, memory strength, and meta-WM judgment

What is the relationship between the neural representation of WM (and its uncertainty) and the neural representation of the metacognitive judgment about WM (i.e., meta-WM)? There was a gradient of selectivity in PFC: some neurons were significantly tuned to only one aspect (Fig. S8), and others tuned to both (see example neurons in Fig 4A-B); at the population level, the coding of meta-WM and WM positively correlated across neurons (Fig. 4C).

**Fig. 4.**
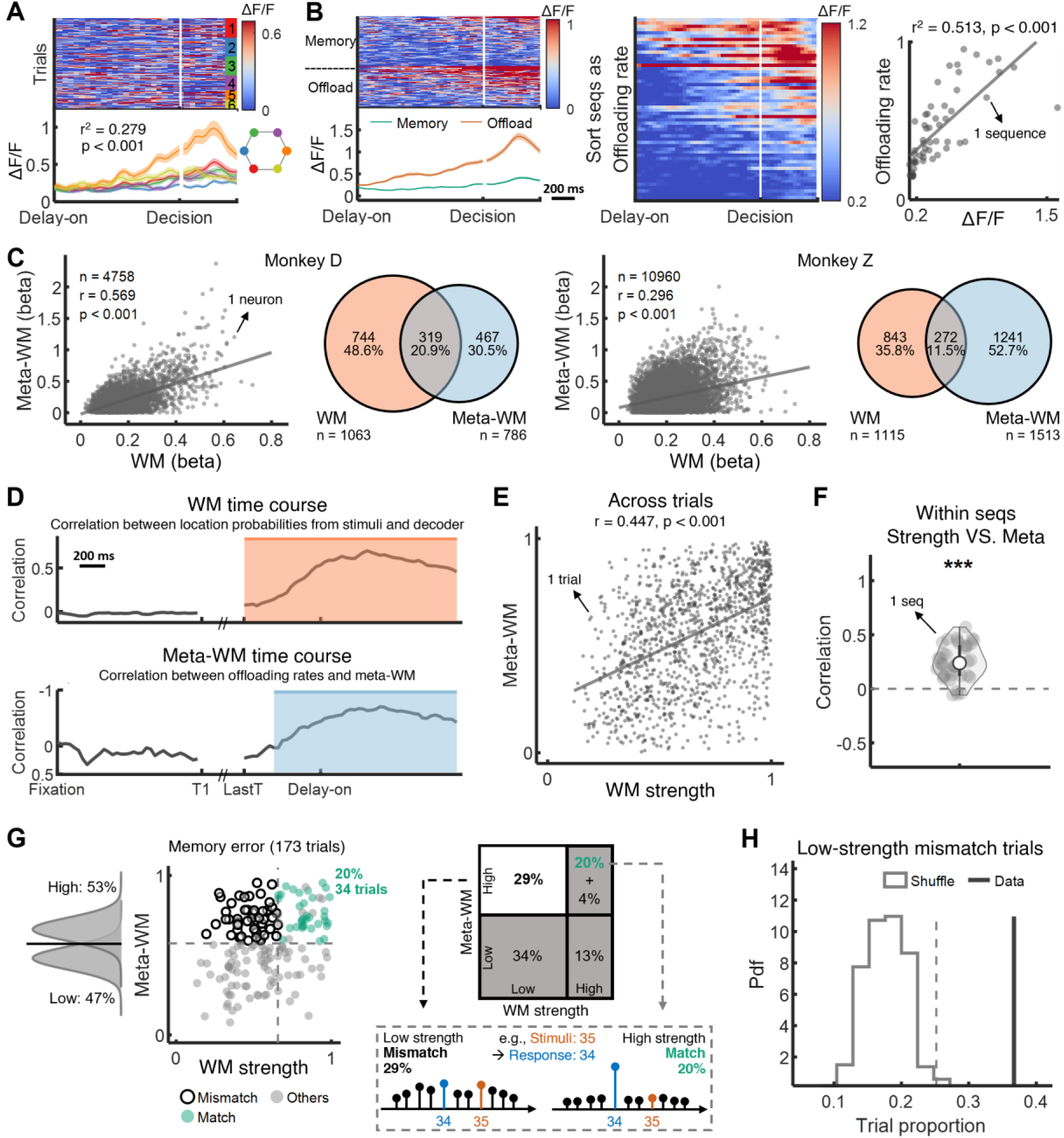
Relationship between WM vs. Meta-WM judgment. (**A-B**) An example neuron with mixed selectivity for both location WM and meta-WM. (A) Single trials (top) and averaged (bottom) neuronal activity grouped by trials with different locations. Linear regression between 6 locations and neuron activity resulted in r^2^ = 0.279, p < 0.001. (B) The same neuron in (A). (Left) Single trials (top) and averaged (bottom) neuron activity of memory and offload decisions. (Middle) Neuronal activity in different sequences sorted by offloading rates. (Right) Linear regression between offloading rates and neuronal activity. Shaded areas represent SEMs. Each dot represents one sequence. (**C**) Neuronal tuning strength (2-norm of beta coefficients of linear regression; see Methods subsection “WM & Meta-WM selectivity of each neuron”) for WM (location memory) and for meta-WM of all neurons and the corresponding Venn diagram of selective neurons from monkeys D (left) and Z (right). Each dot represents one neuron. (**D-H**) Results from an example FOV (FOV 8 of monkey D) using choice (memory + offload) trials. For data covering all FOVs & both monkeys, see Fig. S10. (**D**) Decoding timecourses of WM and meta-WM. Latency of WM (after ‘LastT’, the onset time of last target stimulus): mean = 0 ms, std < 0.001 ms. Latency of meta-WM: mean = 212.287 ms, std = 102.367 ms. Shaded areas are significant time periods, compared with the baseline period (single-tailed one-sample t-test, p < 0.01). (**E**) Pearson correlation between WM strength and meta-WM across individual trials. Each dot represents one trial. (**F**) Pearson correlation between WM strength and meta-WM of trials within each sequence, compared with chance level of 0 (two-tailed one-sample t-test). Each dot represents one sequence. (**G**) (Top) Memory-error trials are categorized into four groups according to high or low levels of WM strength and meta-WM, determined by median values of all choice condition trials. Among error trials, 24% exhibited high WM strength and high meta-WM, including 20% match trials (where the decoding probability of response locations was the highest among all sequences) and 4% others. Meanwhile, 29% showed a mismatch with low WM strength and high meta-WM. (Bottom) Two example memory-error trials with the sequence of stimuli [3-5] and response [3-4] illustrate match and mismatch. The high-strength match trial had concentrated probabilities responding as sequence [3-4], while the low-strength mismatch trial had widely distributed probabilities across many response sequences. (**H**) Test of trial proportion validity for low-strength mismatch trials in memory-error trials. Black line represents real data. Gray dashed line represents 99th percentile of a shuffle distribution. In each resampling iteration, trials were randomly sampled from all choice trials to evaluate the probability that the observed proportion of low-strength mismatch trials arose by chance or from random noise.

The fact that both WM and meta-WM are coded in the same population could promote that the neural representation of meta-WM is informed by the neural presentation of WM and, in particular, memory strength. If WM indeed informs meta-WM, one prediction is that the time course of decoding WM should parallel that of meta-WM and, in fact, precede it. This was the case indeed, with decoding of WM that occurred significantly earlier than meta-WM (212.3±102.4 ms, Figs. 4D and S8E). Another prediction is that when the strength of WM is higher, the meta-WM score should be higher; this correlation was clearly observed (Fig. 4E, p < 0.001). This correlation is partly driven by the fact that WM strength and the meta-WM score differ across sequences (Fig. S6C), but it remained significant across trials within each sequence (Fig. 4F, p < 0.001; Fig. S10 for results from other FOVs). Thus, these results indicate that the representation of WM and the associated uncertainty (rather than just the sequence type) informs the metacognitive judgment in the LPFC. Nevertheless, we should also notice that this process is not perfect: the correlation between meta-WM and WM strength could be stronger. Meta-WM may sometimes fail to be correctly informed by WM strength, which would result in error trials (e.g., meta-illusion^29,30^). We tested this possibility by analyzing single error trials, where monkeys chose to perform the task (memory condition) but made mistakes in their retrieval. When analyzing error trials, however, there could be another cause of error trials: monkeys could retain incorrect locations with strong WM strength and thus with a high meta-WM score. In this case, there would be no mismatch between WM and meta-WM.

A total of 173 error trials from a single example recording session in one FOV were included. Using the trial medians of WM strength and meta-WM from all choice condition trials (Fig. 4G) (also see Fig. S9 for other dividing criteria), 53% (93 out of 173) of error trials were classified as high-score meta-WM. In these 93 trials, we found that 34 trials showed high WM strength while encoding incorrect response sequences (Fig. 4G, green dots, the decoding probability of the *response* locations was the highest among all the sequences), suggesting the monkeys maintained the incorrect locations with strong WM strength and thus high-score meta-WM during the delay period. We thus excluded these error trials, as there is no mismatch between WM and meta-WM. Importantly, we still found a significant proportion (36%, 50 out of 139) of mismatch error trials (Fig. 4G, black circles), where weak WM strength erroneously resulted in a high meta-WM score, and this erroneous mapping was not due to internal noise (Fig. 4H, tested against a shuffled distribution, p < 0.01). The results of error-trial analyses from other FOVs for two monkeys are shown in the Figure. S10C.

Taken together, these results from the correlation and error-trial analyses indicate that there could be additional processes contributing to meta-WM beyond WM, and an incorrect mapping to meta-WM could result in errors in single trials.

### Baseline activity also contributes to the metacognitive judgment

Is the imperfect relationship between meta-WM and memory strength caused by a noisy readout of memory strength or additional intervening processes? We examined whether the arousal state and prior confidence of monkeys before sequence presentation could bias the meta-WM judgment^21,26,27,31,32^. We analyzed pupil dilation during the baseline period as it is often a behaviorally relevant proxy for arousal^31,32^. Pupil size during the baseline period was associated with different upcoming metacognitive judgments (offload or not) on the current trial (Figs. 5A-B; see other sequence lengths and the other monkey in Fig. S11). To test whether monkeys could derive prior confidence about the quality of their retrievals in past trials^33,34^, we assumed that the prior confidence corresponds to a leaky mean of past rewards, in which more recent trials have an increasing weight on the prior^35,36^ (see Methods). We found that the reward from the previous two (monkey D) or five (monkey Z) trials influenced the meta-WM judgment on the current trial (Fig. 5C).

**Fig. 5.**
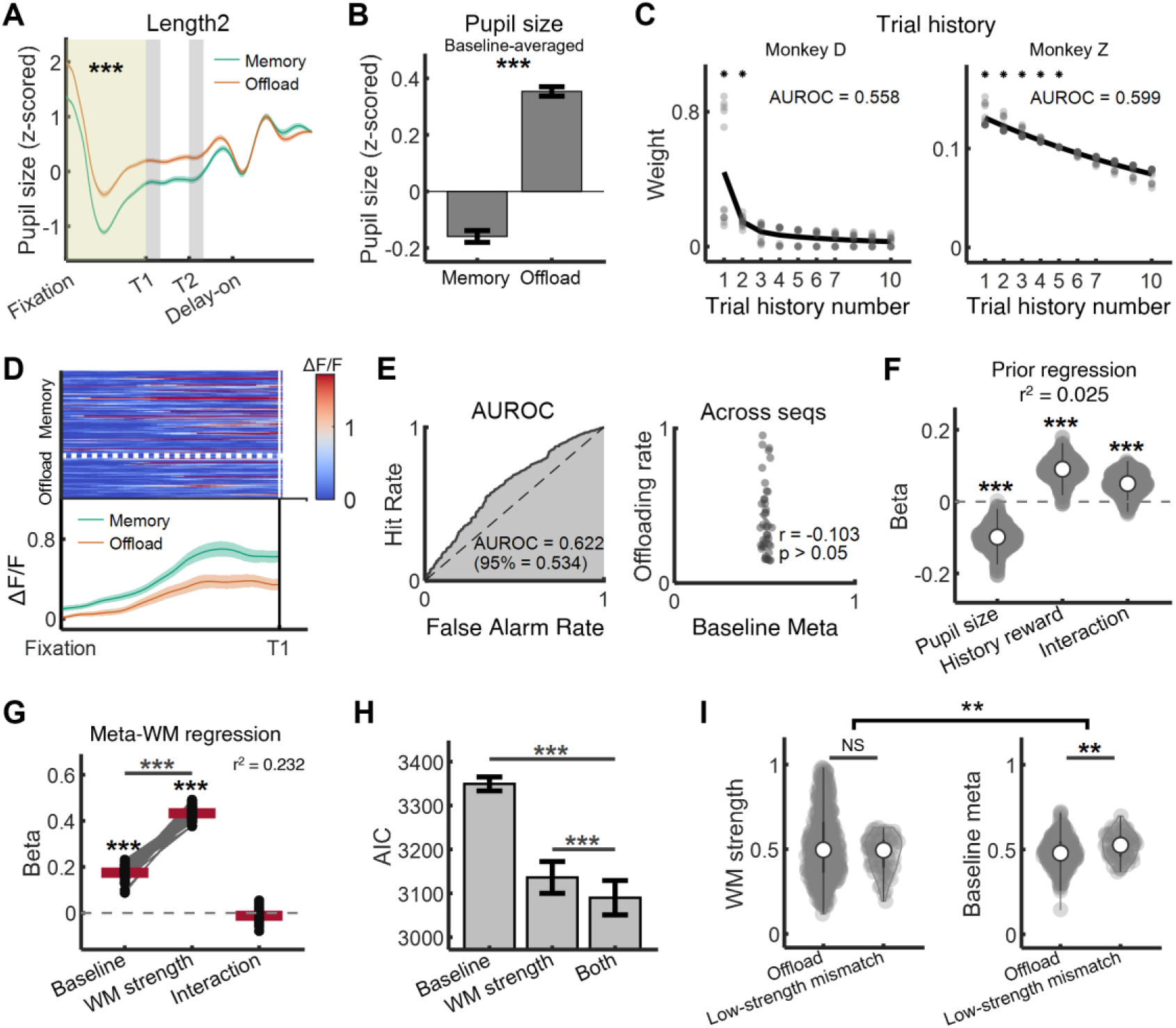
Neural activities during baseline predict meta-WM judgment. (**A**) Time course of pupil sizes under memory trials and offload trials, and comparison between them during the baseline period (yellow areas), taking length 2 trials as an example (***: p < 0.001, two-tailed t-test). Shaded areas around time course lines represent SEM across sessions. (**B**) Comparison between baseline-averaged pupil sizes of the memory trials and those of the offload trials (***: p < 0.001, two-tailed t-test), including trials from all lengths. (**C**) The distribution of trial history weight (see Methods subsection “Trial history”). Each dot represents fitted weight from one recording session. Stars represent the significant weights with p < 0.05, compared with chance level of 0.1 (two-tailed t-test). (**D**) An example neuron exhibiting decision selectivity during the baseline period. (Left) Single trials (top) and averaged (bottom) neuron activity of memory and offload decisions. Shaded areas represent SEMs across trials. (**E**) Performance of the baseline meta-WM decoder. (Left) AUROC was 0.622, and 95th percentile of the shuffle distribution of AUROC was 0.534. (Right) Pearson correlation between offloading rates and sequence-averaged baseline meta-WM. Each dot represents one sequence. For (E-I), data from choice trials of the FOV 8 (monkey D). (**F**) Prior (baseline meta-WM) linear regression, using coefficients of pupil size and history reward. Both coefficients and their interaction term significantly impacted baseline meta (two-tailed t-tests). Each dot represents a coefficient from a random resampling-based regression model, with red lines representing mean values. ***: p < 0.001. (**G**) Meta-WM linear regression, using coefficients of baseline and WM strength. While both baseline meta-WM and WM strength (measured during the delay) are positively impacted meta-WM (single-tailed one-sample t-tests), WM strength was a strong predictor of meta-WM overall (two-tailed paired-sample t-tests between coefficients). Each dot represents a coefficient from a random resampling-based regression model, with red lines representing mean values. ***: p < 0.001. (**H**) Comparison of Akaike information criterion (AIC) in three meta-WM linear regression models (***: p < 0.001, two-tailed t-test). Models with lower AIC are considered to have higher performance, as AIC incorporates a penalty for the number of parameters. The input factors were baseline meta-WM, WM strength, and both, respectively. Error bars represent STDs. (**I**) (Left) Comparison between the offload trials and the low-strength mismatch (subset of memory-error trials) trials in WM strength (left) and baseline meta-WM (right), and in the difference between WM strength and baseline meta-WM (bracket over left & right panels). Each dot represents one trial. All statistical tests are two-tailed t-tests. NS: non-significant. **: p < 0.01.

The neural activity during the baseline period of a large proportion of neurons (monkey D: 22.5% and monkey Z: 19.0%; see Methods) also covaried with the upcoming metacognitive judgment (see an example neuron in Fig. 5D). The metacognitive judgment could actually be successfully decoded from population activity during the baseline in the two monkeys, which we refer to as the “baseline meta score” (Figs. 5E, left and S12A). As a control, we verified that the average baseline meta score across presentations of each sequence type did not correlate with the offloading rate across sequences (Figs. 5E, right, and S12A, p > 0.05), which had not yet been presented to the monkeys. The baseline meta score recapitulated the effect of pupil-linked arousal and trial history (Fig. 5F). We determined the extent to which the baseline meta score and WM strength accounted for the meta-WM score with a linear regression analysis. Both components contributed to the meta-WM score, with a stronger effect of WM strength and no interaction between the two components (Figs. 5G, S12B, and S13). Model comparison indicated that the two components provide a better account of the meta-WM score than each component in isolation (Figs. 5H, S12C), suggesting that the baseline effect contributes to metacognitive judgment in addition to WM strength. The baseline activity accounted for some error trials that resulted from a high meta-WM score with a weak WM strength (Fig. S14). Compared to the offload trials, which generally have weak WM strength and low meta-WM scores, the error trials displayed similar WM strength but significantly higher baseline meta scores (Fig. 5I). These results suggest that baseline activity is an additional, sometimes misleading factor contributing to meta-WM.

### Three metacognition components in the PFC

Finally, we asked whether the prefrontal populations of neurons are highly overlapped or separated in encoding the three metacognition components (baseline, WM, and meta-WM). At the single-neuron level, we identified three groups of component-selective neurons in the lateral prefrontal cortex (LPFC) that included a significant number of mixed-selective neurons (Fig. 6A), which showed a preference for two or three components. At the population-neuron level, we examined whether each component could be linearly separated in the population activity. We searched the line in the n-dimensional activity space along which baseline activity was separated between memory and offload trials, and we termed this line the baseline subspace^37^; similarly, we searched the WM strength subspace and meta-WM score subspace in the population neurons (Fig. 6B; see Methods). These one-dimensional subspaces each explained a significant amount of variance (40.1%, 52.4%, and 71.5% for the baseline, WM, and meta-WM, respectively, from FOV 8 of monkey D; see other FOVs from two monkeys in Fig. S15). Importantly, those subspaces were oriented in a near-orthogonal manner, as evident by the large principal angles between them (Fig. 6C). To further quantify the degree of alignment across different subspaces, for any two components—e.g., baseline and WM—we calculated the variance accounted for (VAF) ratio^9,11^ by projecting the data from the baseline subspace to the WM subspace and computing the remaining data variance ratio after the projection. If the two component subspaces are nearly orthogonal, the projection from one subspace will capture little of the variance in the other subspace, resulting in a low VAF ratio. The result showed low VAF ratios for all cross-subspace pairs (Fig. 6D). The orthogonality between WM and meta-WM was slightly lower than other pairs (ps < 0.001), providing again the neural basis for the strong correlation between them. As a control, we randomly split the trials into two halves to obtain separate estimations of each subspace and computed the principal angles and VAF ratio; the orthogonality of the subspaces was lost (Figs. 6C and 6D). The neural dynamics in each subspace confirmed their contributions throughout the duration of a trial (Fig. 6E). Therefore, the results suggested that, at the collective level, the three metacognition components were encoded in separate PFC neural subspaces, although with mixed selectivity in single neurons.

**Fig. 6.**
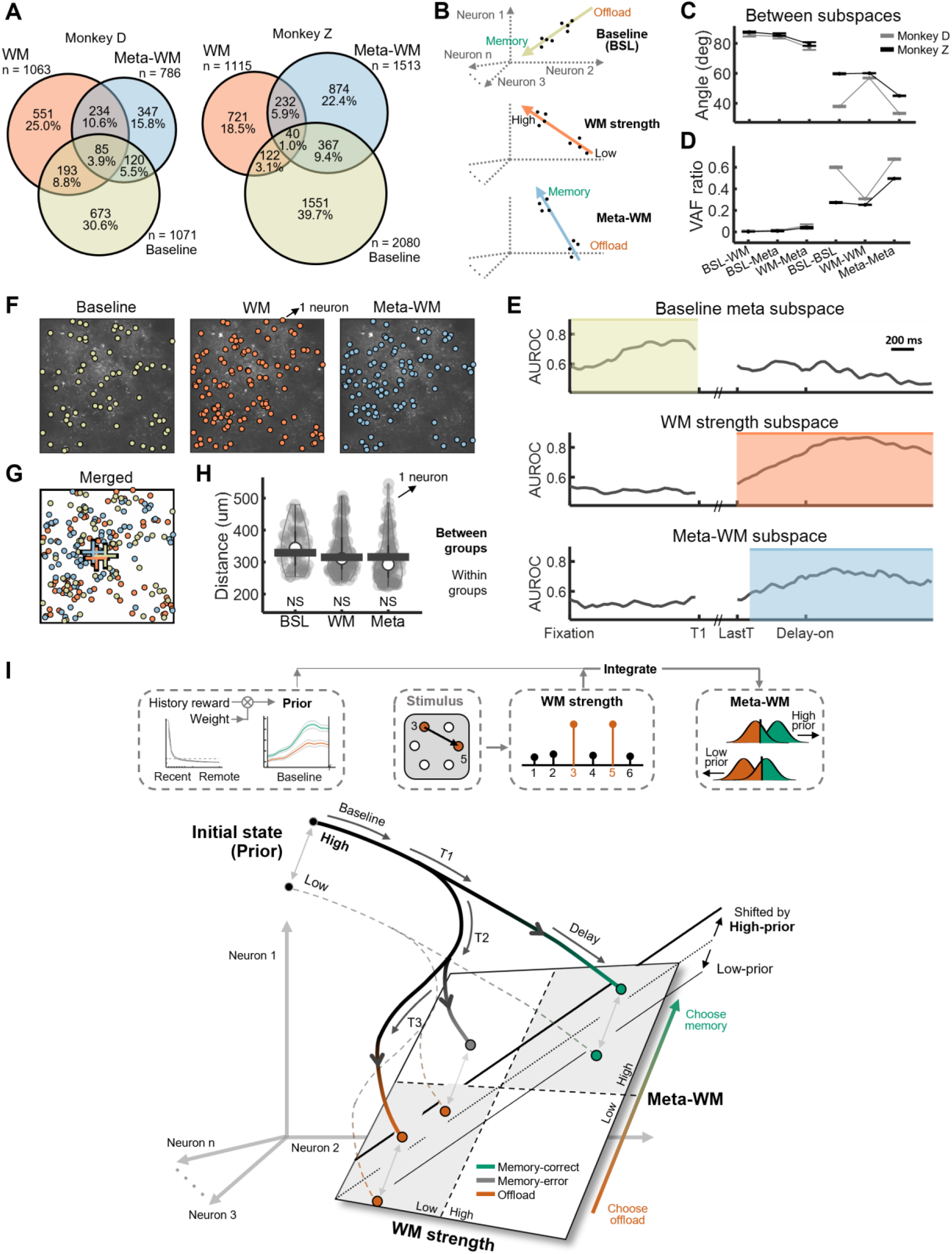
Three task variables in the PFC. (**A**) Venn diagram showing selective neurons from both monkeys related to WM (location memory), meta-WM, and baseline meta (prior), respectively. (**B**) Schematic of subspaces of baseline meta-WM, WM strength, and meta-WM in neuronal state space. For example, the meta-WM subspace represents an axis derived from the meta-WM decoder that captures meta-WM scores across trials, forming a vector within the neuronal state space (see Methods subsection “Subspace estimation and principal angles between subspaces”). BSL: Baseline. (**C**) Principal angles between different subspaces in (B). As a control, we randomly split trials in half to obtain two separate estimations of each subspace and computed their angle. deg: degree. Each FOV was individually analyzed; error bars represent SEM across FOVs. (**D**) Variance accounted for (VAF) ratio between different subspaces (see Methods subsection “VAF ratio between subspaces”). Each FOV was individually analyzed; error bars represent SEM across FOVs. (**E**) Time courses of AUROC to differentiate between memory and offload trials within the baseline meta subspace and the meta-WM subspace, as well as between low-strength and high-strength trials within the WM strength subspace. Shaded areas in the WM strength and meta-WM subspaces indicate significant time periods, compared to the baseline period (single-tailed one-sample t-test, p < 0.01). (**F-G**) (F) Spatial organization of selective neurons for baseline meta, WM, and meta-WM (from FOV 4 of monkey D; 66, 120, 108 neurons, respectively). Each dot represents a neuron. In the merged panel (G), crosses represent spatial centroids. (**H**) Comparison of spatial distances within groups and between groups (two-tailed t-test; FOV 4 of monkey D). Each gray dot represents the average spatial distance between one neuron and others within the same group in (F). The black bar represents a calculation similar to that of the gray dots, but specifically for results between different groups, and it shows the median value among neurons. NS: non-significant. (**I**) Illustration of sequential mapping from WM strength to meta-WM and prior integration effect. In the PFC neural state space, the baseline signal influences neural states along the meta-WM judgment axis. With incoming external WM signals, these states shift sequentially along the judgment axis, owing to sequence-specific features (e.g., length) and additional, sequence-independent processes. The linear integration of baseline and WM signals’ projections onto this axis results in the final meta-WM decision controlling behavior.

Two-photon imaging enabled us to explore whether the neurons representing metacognition components are anatomically segregated in the PFC. An example FOV demonstrated that the three groups of component-selective neurons were nearly uniformly distributed (Fig. 6F and 6G). For each neuron, we first calculated the spatial distances to other neurons and then compared the intra-group distances with the distances between groups (see Methods^38^). We found that the spatial distances within the groups showed no significant difference compared to those between the groups for all three groups of neurons (Fig. 6H, ps > 0.05). This suggests that the three components were anatomically intermingled in the PFC. Similar results are observed in other FOVs and the second monkey (Fig. S16; see results in Fig. S17 for defining selective neurons based on their contributions to corresponding subspaces). Therefore, while the metacognition components were functionally separate in subspaces of the PFC population activity, they remained anatomically intermingled.

Finally, we summarize our findings by proposing a conceptual model for the metacognitive WM system implemented in PFC (Fig. 6I). The representations of metacognitive processes were composed of neural activities and their dynamics in the separate neural ensembles of baseline, WM, and meta-WM. The baseline activity already shifts the neural states along the meta-WM judgment axis (regarding whether to offload or not; note that the effect of baseline activity is additive and does not interact with the upcoming WM strength). Then, as the sequence is presented and stored in WM, sequence-specific features (e.g., spatial locations in the current task) and trial-by-trial WM-specific features, further push the neural state along the metacognitive judgment axis, owing to the correlation between WM-strength and meta-WM in the neural space. Presumably, on the WM trajectory, each point, jointly representing the WM locations and their uncertainty, can be further unfolded as the vectorial combinations of neural representations of multiple location signals, which were likely propagated from posterior sensory regions. Specifically, as the sequence difficulty (e.g., sequence length) escalates, the WM strength (e.g., tuning strength of spatial location) weakens due to limited resources. Thus, the projection of the neural state onto the metacognitive judgment axis leads to the final meta-WM decision to control behavior.

## Discussion

We recorded tens of thousands of neurons in the lateral PFC of macaque monkeys performing the spatial delay-sequence reproduction meta-WM task. We found that PFC neuronal responses could represent two aspects of WM: 1) the remembered locations and the associated uncertainty in WM, which corresponds to WM strength, through population coding, and 2) the metacognitive judgments of WM uncertainty about the remembered locations, which predicted the probability of offload decisions. Furthermore, neural activity during the baseline period, resulting from the internal neural states and history of rewards, also predicted metacognitive judgments on top of memory strength. Crucially, these three components of metacognition in WM—baseline prior, WM, and meta-WM—were functionally represented by three distinct subspaces of the population activity of anatomically intermingled neurons. We thus proposed that neural activity in the macaque PFC underlies the process of metacognition in WM by integrating several components of the metacognitive judgment, providing a novel and unifying model of metacognitive computations.

### Meta-WM, memory strength, and meta-memory cues in the PFC neural population

The current study identifies a neural implementation of metacognition that integrates first-order and cue-based components that have previously been studied separately in neuroscience^7,21,26^. In principle, metacognition should benefit from the first-order information about uncertainty carried by decision or memory systems. However, this is only possible if neural circuits make first-order uncertainty available to the metacognitive system. If not, then metacognition has no choice but to infer the probability that a decision or memory is correct from indirect cues. Experimental manipulations or models of uncertainty^6,39,40^ have provided evidence that first-order uncertainty contributes to metacognitive judgments, but it is more challenging to demonstrate this contribution on a trial-by-trial basis. Recent advances in neuroscience have made it possible to decode neural representations of (first-order) sensory or memory data and their associated uncertainty, then compare the decoded uncertainty with decisions^41^ or confidence reports^7,42^, and ultimately conclude that first-order uncertainty is available for higher-order processes such as metacognition. However, such a conclusion requires ruling out the possibility that the contribution of first-order uncertainty could be explained by other factors that serve as indirect metacognitive cues. The existence and use of such cues are well established, particularly in the metamemory literature^25–28^. Our results show that arousal and past performance are indeed cues to metacognitive judgment, but their contribution was in addition to memory strength.

Our results show not only that two components (first-order and cue-based) contribute to metacognition, but also that they coexist and are integrated in the lateral prefrontal cortex. This finding implies that the LPFC is capable of representing different reference frames^21^. One is world-centered, corresponding to whether the content of WM is informative about the presented sequence. The other is self-centered, corresponding to whether subjects feel confident about their memory recall. Our results suggest that these reference frames correspond to distinct subspaces of LPFC population activity.

We used two approaches^43^ to identify first-order and cue-based components of metacognition in the LPFC. One approach is correlational, aiming to identify neural correlates of metacognitive judgments^13,15,40,44–46^. To this end, we trained a decoder of the monkey’s offload decision (a self-centered reference frame). The other approach is code-driven: we decoded the memory strength from a population code of the presented sample locations (this reference frame is world-centered). This approach has been used previously, assuming a probabilistic population code, to decode perceptual uncertainty^41,42^ or WM uncertainty^7^ from sensory cortices. In contrast, correlates of metacognition have typically been reported beyond sensory cortices using the correlational approach. In previous studies, these correlates were often related to either metacognitive judgment or its components^16,23,40,41,44,46–48^. Some studies have come close to combining the two components, e.g., Okazawa and colleagues found that neural geometry in the parietal cortex encoded not only the (first-order) decision variable but also task difficulty (a potential cue for metacognition), but the latter did not affect monkey confidence^24^. Here, we extended these previous findings by combining both approaches (code-driven, correlational) and identifying neural representations of metacognitive judgment and its components (first-order, cue-based) in activity subspaces of the same region, the LPFC, thereby providing a unified framework that integrates the neural representation of uncertainty and the neural representation of performance judgment in WM.

### Potential computational mechanisms of metacognition in the PFC

Apparently, how the metacognitive process is implemented and shaped within the neural landscapes of the prefrontal circuit remains only partly understood. Nevertheless, based on our neural data, we can posit that: 1) the potential multiple neural subpopulations (baseline, WM and meta-WM), with certain mixed-selectivity, in the prefrontal circuits may serve as the mechanistic underpinning for the information transition in different neural subspaces^49,50^; the WM (locations and their uncertainties) and prior signals could be firstly encoded in other brain regions such as visual, parietal or subcortical cortices and then propagated to PFC for maintenance. 2) These multiple routes converge in the PFC and interact through a structure-enabled gain modulation mechanism with meta-WM to inform metacognitive judgment^51^, suggesting that the metacognitive computation could be implemented within the PFC rather than through reciprocal connections between brain regions. However, due to the lack of recordings from the sensory cortex, the present results do not exclude the simultaneous presence of the metacognitive computation within the PFC and between brain regions. Multiple circuitries for metacognition in WM may coexist within the brain. 3) The nature of this interaction or mapping may be predetermined by the anatomical connectivity between populations, likely established during task learning in monkeys; this connectivity may differ between subjects and partly account for differences in meta-WM performance. Examining the development of connections among different neural populations in the PFC and across brain regions during learning is crucial. While further causal interventions are needed for each metacognitive component (the current study only shows correlation), this proposal necessitates additional theoretical and experimental explorations in the broader domain of metacognitive computations.

## Acknowledgments

We thank Marion Rouault, Gouki Okazawa, Xi Jiang, and Zhenghe Tian for their critical comments on the manuscript. This work was supported by the STI2030-Major Project (2021ZD0204102), the National Science Fund for Distinguished Young Scholars (32225022), the CAS Project for Young Scientists in Basic Research (YSBR-071), and the Shanghai Municipal Science and Technology Major Project 2021SHZDZX to L.W. FM is supported by a European Research Council grant (ERC StG 947105-NEURAL-PROB).

## Materials and Methods

### Experimental models

Two healthy male monkeys D and Z (Macaca mulatta, weight: 8.5/10 kg, age: 8/9 years) participated in the study. Both monkeys were housed in individual cages. Food was available ad libitum, while water intake was restricted to daily requirement in the cage. On testing days, monkeys received juice rewards for correct responses during the experiments. All experimental procedures were approved by the Animal Care Committee of the Institute of Neuroscience, Chinese Academy of Sciences (CEBSIT-2020035R02).

### Behavioral task

#### Visual stimuli

All visual stimuli were presented on a 24-inch touch monitor (Dell P2418HT) with a refresh rate of 60 Hz and a resolution of 1920×1080. The stimuli were generated using Matlab (MathWorks, MA, USA) with extensions from Psychtoolbox^54,55^. In this study, stimuli were created from six spatial locations arranged in a hexagonal configuration. In each trial, locations were presented sequentially on the screen, with sequence lengths ranging from 1 to 4 trial-by-trial randomly (see Fig. 1A). Each spatial location (if sampled) would be sampled only once per sequence, and each sequence would be exhibited in a clockwise direction, such that sampled locations with smaller numerical labels would always be shown before those with larger labels (for example, if locations 2, 4, 5 were sampled, the sequence would always be 2-4-5). For both monkeys, there were a total of 56 sequences: 6 (or 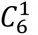) length-1 sequences, 15 (or 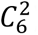) length-2 sequences, 20 (or 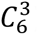) length-3 sequences, and 15 (or 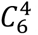) length-4 sequences. All possible combinations of stimulus locations were used in the experiment and each trial randomly involved only one sequence. Given the relatively low task performance for length-4 sequences in the training session, we primarily focused on sequences of lengths 1, 2, and 3 in the two-photon imaging experiment, However, for consistency, a limited number of length-4 sequences were still included during imaging experiment for behavioral analysis. The ratios of the four sequence lengths (length 1, 2, 3, and 4) in all trials were 20:20:20:9 for Monkey D and 20:20:20:3 for Monkey Z.

#### Task structure

Monkeys were trained to perform a delayed-sequence reproduction meta-WM task. The monkeys were required to make behavioral reports using eye saccades to appropriate locations on a monitor screen. Each monkey was seated in a primate chair positioned 35 cm away from the monitor during the experiments. All experiments were carried out in a dark room when the light was off. Here, we used an Eyelink 1000 Plus eye-tracking system with a sampling rate at 1000 Hz to track the monkeys’ gaze positions during each trial.

Each trial began with the monkey fixating on a central cross (0.98° in diameter), which had to be maintained within an invisible window with a radius of 2.57°. The monkeys maintained this fixation for a baseline period of 1.1 seconds during which a hexagonal layout of six empty circles (each 4.08° in diameter) were presented, positioned 11.55° from the fixation point. During the subsequent sample period, 1 to 4 red stimuli were sequentially displayed on the screen in a clockwise manner, each displayed for 0.2s with a 0.4s inter-stimulus interval. This was followed by a delay period (1.5-1.7 s), during which the layout disappeared, leaving only the central cross visible. Subsequently, in the decision period, each trial would fall under either the “choice” condition (65%) or the “forced-to-test” condition (35%). Under the “choice” condition, the monkeys were required to select via saccade between a green square representing “memory” option (no hints during the response) and a red triangle representing “offload” option (previously displayed locations reappear as hints). The choice symbols (green square / red triangle) appeared randomly on either the left or right side of the screen from trial to trial to prevent anticipatory motor preparation signals during the delay period. Under the “forced-to-test” condition, only the “memory” option was shown randomly on the left or right side. After the decision, there was a second delay period (another 1.5 - 1.7 s), which was not included in the analysis. Note that in this study, “delay period” only denotes the first delay, i.e. the interval between the sample and the decision periods. Following this second delay, the response period began with the hexagonal layout reappearing on the screen. All circles were empty under the “memory” and “forced-to-test” conditions, while in the “offload” condition, the previously shown sample stimuli were displayed again as hints. In summary, each trial included the following periods: baseline, sample, delay, decision, second delay, and response.

The monkeys’ behavior varied during the response period depending on the condition of the decision period. In memory trials, monkeys were required to make saccades to the targets that did not appear during the sample period in a clockwise order. After selecting all the correct targets, they returned their gaze to the central fixation cross to submit the trial. If correct, they received a reward of 3 drops of water. A correct trial necessitated responses in a clockwise order. In offload trials, all red stimuli from the original sequences reappeared simultaneously. The monkeys first made saccades to these cued locations (behavior akin to note-taking), then skipped over them and made saccade to the remaining targets. Upon successful completion, they received a reduced reward of 2 drops of water. Monkey D received 3 water drops for correct responses under memory and forced-to-test conditions, 2 for correct offload trials, and 0 for error trials, following a 3-2-0 reward scheme; monkey Z was rewarded according to a 2-1-0 scheme.

On a single experimental day (one session), several runs (typically 5 runs) were conducted. Each run comprised one-third forced-to-test trials and two-thirds choice trials. Specifically, for monkey D, each run included 69 forced-to-test trials (with sequence lengths of 1, 2, 3, and 4 involving 20, 20, 20, and 9 trials, respectively), totaling 207 trials per run (69 forced-to-test trials and 138 choice trials). Similarly, each run for monkey Z included 63 forced-to-test trials and 126 choice trials, totaling 189 trials per run. Consequently, each session encompassed approximately 1000 trials (at least 800).

### Two-photon calcium imaging

#### Surgery procedures

Two sequential surgical procedures were performed on each monkey under general anesthesia. During the first surgery, three head posts were implanted to stabilize the head position during subsequent training and recording sessions: two on the posterior of the head and one on the middle of the forehead. A Y-shaped steel frame was then attached to the head posts. Following a recovery period of 10 days, both monkeys were trained to perform the behavioral task over the course of a year until they met established performance criteria (significant correlation between accuracy and offloading rate across sequences).

Following the training period, a second surgical procedure was performed. A 22 mm craniotomy was created over the lateral prefrontal cortex, extending from the principal sulcus (PS) to the arcuate sulcus (AS), approximately corresponding to Walker’s areas 46 and 8^56^. The dura mater (16 mm in diameter) was removed to expose the underlying cortex. Viral injections and implantation of an imaging window were then carried out. Approximately 500 nl of an equal mixture of rAAV2/9-hsyn-tTA (3×10^12 vg/ml) and rAAV2/9-TRE3-GCaMP6f (1×10^13 vg/ml) was injected at a depth of 600 μm to deliver TET-Off GCaMP6f virus^57^ (from Gene editing facility of Institute of Neuroscience, Chinese Academy of Sciences). The injection sites were positioned along the PS.

The imaging window unit was constructed by attaching an 18 mm diameter, 0.17 mm thick glass coverslip to a titanium ring (15 mm outer diameter, 13 mm inner diameter). The unit was carefully placed on the cortical surface, with the titanium ring secured to the skull using dental acrylic. To protect the coverslip, a steel shell was placed over the entire imaging window.

#### Set-up

In vivo two-photon calcium imaging was performed using a Thorlabs two-photon microscope and a femtosecond laser (Chameleon Discovery, Coherent). A 16× objective lens (0.8 N.A., Nikon) was used to image a 700 μm × 700 μm area at 30 frames per second. The recording depth ranged from 80 μm to 250 μm below the pia, and each imaging experimental day (session) lasted 3–5 hours. We aimed to cover most regions with GCaMP6f expression within the recording windows located along the principal sulcus (PS). A total of 28 fields of view (FOVs) were recorded: 12 FOVs from monkey D (4,758 neurons) and 16 FOVs from monkey Z (10,960 neurons); for the position and neuron count of each FOV, see Figure S2 and Table S1.

#### Imaging data processing

We implemented the image processing pipeline using MATLAB and Python. Images were first motion-corrected using a rigid motion-correction algorithm from the NoRMCorre package^58^. Source extraction was performed using the Suite2p package^59^, which is based on singular value decomposition (SVD). Scores^59^ were calculated for each extracted spatial component, and regions of interest (ROIs) marking putative neuronal locations were selected by thresholding these scores. The resulting fluorescence traces were sampled at a frame rate of 30 Hz. The ΔF/F was calculated as (F - F₀) / F₀, where F represents the raw fluorescence signal and F₀ is defined as the mode value within the preceding 30-second window.

#### ROI alignment of two-session recording of the same FOV

In this study, we recorded the same FOV across two consecutive sessions to obtain additional (double) trials for further analyses. Thus, a total of 27 FOVs were examined over 54 recording sessions, with the exception that FOV 3 of monkey D was imaged in 3 consecutive recording sessions. After processing the imaging data, the ROIs from both sessions were aligned and merged to double the number of trials for each ROI.

To assess the similarity between ROIs across two consecutive recording sessions (A and B) for alignment purposes, we employed a weighted composite similarity score incorporating both structural and functional features:

(1) Median position (weight = 1): Spatial proximity of ROIs, with a threshold of less than 10 pixels in median position distance.
(2) Size (area) (weight = 0.5): Comparison similarity of the area encompassed by each ROI.
(3) Aspect ratio (weight = 0.5): Evaluation of the shape characteristics (x pixel size / y pixel size) of the ROIs.
(4) Stimulus-evoked neuronal activity (weight = 2): Assessment of the similarity in neuronal responses to each stimulus sequence.

For each term listed above, we first generated a similarity queue sorted in descending order. Subsequently, we assigned scores ranging from 20 to 1 to each item in the queue, multiplying each score by the corresponding term weight. The alignment procedure comprised the following steps:

(1) Session A to session B: For each ROI in session A, identify up to top 20 ROIs in session B within a 10-pixel median position distance. Rank these candidate ROIs based on the weighted composite similarity score.
(2) Session B to session A: For each ROI in session B, identify up to top 20 ROIs in session A within a 10-pixel median position distance. Rank these candidate ROIs based on the weighted composite similarity score.
(3) Identification of overlapping A-B pairs: Determine the set of ROI pairs that are mutually identified in both steps above, indicating a bidirectional match.

### Behavioral and single-neuron analysis (Related to Figure 1)

#### Accuracy and offloading rate

The length-averaged accuracy (Fig. 1B) was calculated by dividing the number of correct trials by the total number of trials in the relevant conditions (offload, memory, forced-to-test) for each length. The sequence-averaged accuracy was determined by dividing the number of correct trials by the total number of trials in the forced-to-test condition for each sequence (Fig. 1D). The sequence-averaged offloading rate was determined by dividing the number of offload trials by the number of choice trials for each sequence (Fig. 1D).

#### Behavior recall variability

In this study, we quantified working memory (WM) representations in terms of their content (locations) and strength. Behavior recall variability was measured by the normalized entropy of the probability distribution of response sequences following the presentation of a stimulus sequence during the sample period.

We computed *entropy* in a classical manner, using the distribution of all possible response probabilities of sequences, where:

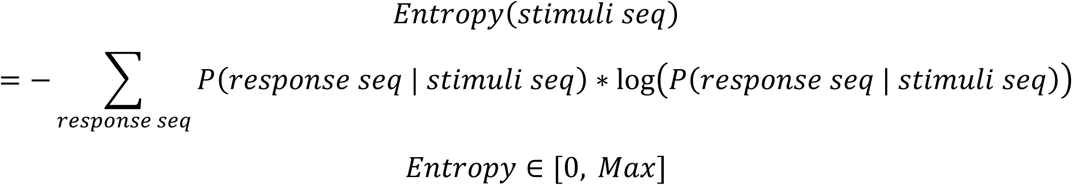

Here, we also compute the theoretical upper bound of *entropy* by calculating the entropy of a uniform distribution, denoted as *Max*. Finally, we determined the recall variability by normalizing the entropy from [0, Max] to [0, 1], where:

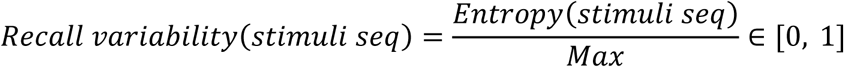

In this context, a recall variability of 1 represents a uniform probability distribution (which would be noisy WM), and a recall variability of 0 indicates that the probability of one specific sequence is 1 (representing very high precision of WM).

For example, in Figure 1C, responses to the sequence [1-3] of monkey D were mostly concentrated around its corresponding physical stimulus, indicating low variability. In contrast, responses to the sequence [2-5-6] were widely distributed, reflecting high variability.

#### WM selectivity of each neuron

We conducted linear regression for each neuron to quantify its tuning properties as follows:

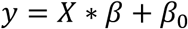

For the regression of location WM, *y* represented the neuronal activity for each trial (a column vector with 1 by number of trials), *β*_0_was the intercept term, *β* was the regression coefficient to be fitted (a 1-by-6 vector, indicating weights of 6 locations), and *X* represented the stimulus sequence for each trial (a design matrix with 6 by number of trials, with a one-hot-like 1-by-6 vector for each trial, e.g. stimulus sequence [3-5] would be [0 0 1 0 1 0].). Only correct trials (in both the memory and forced-to-test conditions) were used in the regression of location WM.

After conducting linear regression, we obtained *β*, *r*^2^(the coefficient of determination, representing regression explained variance), and *p* value of regression (representing the overall effectiveness of location *β*s to predict neural activity) for each neuron. Those neurons whose *p* values of linear regression were less than 0.01 would be considered location WM selective neurons. We further computed 2-norm of 6 location *β*s to evaluate tuning strength of each neuron, thereby summarizing the six coefficients by a single scalar metric.

#### Decision selectivity of each neuron

We conducted linear regression for each neuron to quantify the tuning properties as follows:

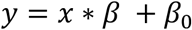

For the regression of decision, *y* represented the neuronal activity for each trial (a column vector with 1 by number of trial), *β*_0_ was the intercept term, *β* was the regression coefficient to be fitted, and *x* represented the behavioral label for each trial (0 for offload and 1 for memory).

After conducting linear regression, we obtained *β*, *r*^2^(the coefficient of determination, representing regression explained variance), and *p* value for each neuron. Those neurons whose *p* values of linear regression were less than 0.01 would be considered decision selective neurons.

### Neural activity-based WM analysis (Related to Figure 2)

#### Location probability distributions of behavior

As mentioned in the “Behavior recall variability” section above, we computed stimuli-to-response matrix of behavior data from the forced-to-test condition (Fig. 1C), including both correct trials (diagonal) and error trials (non-diagonal). Each element of the matrix denotes the probability for a monkey to execute one particular sequence given the presentation sequence during the sample period.

Based on this matrix, we could in turn compute each stimulus location’s behavioral probability distribution from the stimuli-to-response matrix of behavior data (Fig. 2C). Each element of the matrix denotes the probability for a monkey to execute one particular location given the presentation of a specific sequence during the sample period.

#### Neural decoder-based WM strength

WM strength was defined as the inverted normalized entropy of the distribution of all possible decoded probabilities of sequences, given the neuronal population activity.

We trained six linear binary Support Vector Machines (SVMs) as sub-decoders to decode the probability of each location being represented by neuronal data, in which the labels for training were binary (0 or 1; from stimulus label) and then in testing set the output of the decoder was the probability. Given the time-averaged neuronal activity during delay, we trained the decoder on correct trials and tested it on both correct trials and error trials, using location-balanced resampling and K-fold (K=5) cross-validation. Each SVM sub-decoder was optimized using the MATLAB function ‘fitcecoc’. This allowed us to predict the location probability distribution for each trial. Different sub-decoders were trained for trials of different sequence lengths.

The chance level of Pearson’s correlation between location probability distributions of behavior data and those predicted by neural decoders of locations was computed via substituting decoded location probability distributions with random values sampled from a uniform probability distribution.

We then used the decoded location probability distributions to compute decoded probability of each sequence with multiple items through a joint probability approach, which involved calculating the product of all possible locations’ probabilities, with the assumption that each location was encoded independently from the others. We denote the probability of location *i* as *p*(*location* = *i*|*activity*). Take sequence [3-5] as an example, the decoded probability of the sequence was then *P*(*seq* = 35 | *activity*), where:

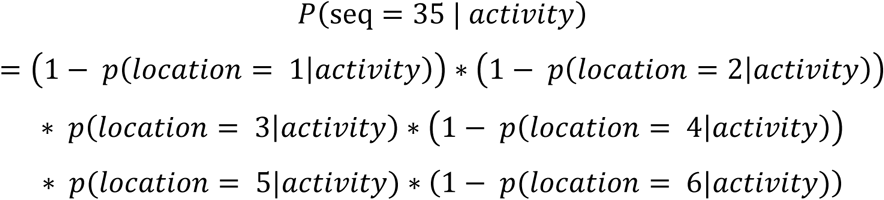

Then, we computed *entropy* in a classical way, through distribution of all possible decoded probabilities of sequences, where:

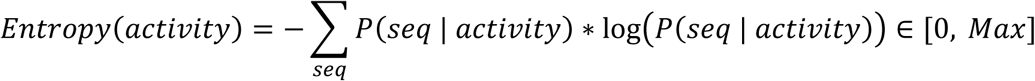

Here, we also compute the theoretical upper bound of the *entropy*, by computing entropy of a uniform distribution, denoted as *Max*. Finally, we computed WM strength by inverting and normalizing the entropy from [0, Max] to [0, 1], where:

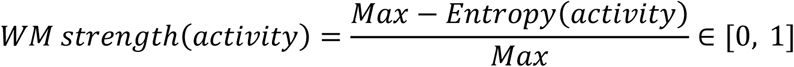

WM strength = 0 represented a uniform probability distribution over all possible sequences and WM strength = 1 represented distributions where probability of one specific sequence came to exactly 1.

For analyses utilizing pseudo-population where data from all sessions were combined, we resampled the same number of trials (including memory correct trials and forced-to-test correct trials) of each sequence type across sessions. The resampling trial numbers across sequences were determined in a sequence length-dependent and location-balanced manner. Within each sequence length, resampling was performed to balance the number of trials across different locations. This resampling was repeated 32 times. For each resampling, we trained and tested a new set of decoders from which to recompute WM strength. After all resampling procedures, we averaged the testing results to produce the final pseudo-population-based evaluations of WM (Fig. S3).

### Neural activity-based meta-WM analysis (Related to Figure 3)

#### Meta-WM selectivity of each neuron

We conducted linear regression for each neuron to quantify its tuning properties regarding putative meta-WM (or how well the monkeys might consider their WM of stimuli locations to be) as follows:

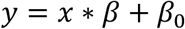

Where *y* represented the neuronal activity for each sequence (a column vector with 1 by number of sequences), *β*_0_ was the intercept term, *β* was the regression coefficient to be fitted (a scalar), and *x* represented the behavioral offloading rate for each sequence (a column vector with 1 by number of sequences).

After conducting linear regression, we obtained *β*, *r*^2^(the coefficient of determination, representing regression explained variance), and *p* value for each neuron. Those neurons whose *p* values of linear regression were less than 0.01 would be considered meta-WM selective neurons. We further computed the 2-norm of *β*, here was an absolute value, to evaluate the tuning strength of each neuron.

#### Neural decoder-based meta-WM

Here, meta-WM is defined as the probability with which a population of neurons encodes a decision of the memory option for a larger reward (likely judging current WM strength to be good).

We used a single linear SVM to decode the probability of choosing memory in the choice trials. The SVM functioned as a multivariable linear regressor by extracting the posterior probability as the output. Here, the input was the time-averaged population neuronal activity during the delay, and the output was the probability of choosing memory for the trials. We optimized the decoder to estimate the meta-WM values of the trials, using resampling to balance trial numbers for memory & offload trials and K-fold (K=20) cross-validation for training and testing. For example, if there are 600 memory trials and 500 offload trials, then in each resampling iteration, we randomly select 500 trials from each condition without replacement. The optimization was performed using the MATLAB function ‘fitcecoc’. Here, the posterior probabilities extracted from applying the optimized decoder over trial-wise data were denoted as meta-WM scores, which represented the probability of choosing memory in current trials.

To further discriminate between memory and offload decisions based on the estimated meta-WM scores from the SVM, we fitted an optimal decision boundary by maximizing the values of (hit rate – false alarm rate), as in Figure S6A. Hit/False alarm rates represent proportions of trials where meta-WM scores exceed the decision boundary in all memory/offload trials, respectively. Trials with meta-WM scores above the boundary were classified as memory, while those below the boundary were classified as offload.

The AUROC curves were plotted as results from a moving decision boundary from 0.01 to 0.99 with a step of 0.001, in which each decision boundary could result in a pair of hit rate and false alarm rate. We plotted all the paired results and connected all the dots to compute the area under the curve. The shuffled AUROC distribution was computed by first shuffling the trial decision labels of “memory” and “offload”, then training and testing meta-WM decoders.

### Analysis of relationship between WM strength and meta-WM (Related to Figure 4)

#### Dynamics analysis of WM and meta-WM

To explore the temporal dynamics of both WM (as in locations of constituent stimuli) and meta-WM of remembered sequences, we conducted two sets of correlations across time frames within each trial, ranging from the last target onset to the late delay. We resampled 16 times to compare latency times as shown in Figure 4D. This resampling procedure aligned with the methods described in the “Neural decoder-based WM strength” and “Neural decoder-based meta-WM” sections.

As illustrated in Figure 2, we used both location-level correlations and sequence-averaged WM strength to assess the neural dynamics of WM. Specifically, we employed a location-level correlation between location probabilities from the decoder (e.g., [0.1, 0.1, 0.9, 0.1, 0.9, 0.1]) and stimuli (e.g., [0, 0, 1, 0, 1, 0]) to track the dynamics of WM, testing when they began to deviate from the baseline. One-sample t-tests were used to determine whether the location-level correlation at each time frame (starting from the onset of the last target to 1.1 seconds into the delay period) had a higher mean value compared to that during the baseline period.

As shown in Figure 3, we computed Peason’s correlation between sequence-averaged behavioral metrics (offloading rates) and sequence-averaged meta-WM scores. This correlation was also used to investigate the dynamics of meta-WM over time and identify when they deviated from the baseline. Again, one-sample t-tests were performed to examine whether the correlation at each time point showed a higher mean value than that of the baseline period.

#### Validity of low-strength mismatch trial proportions in memory-error trials (Fig. 4H)

The shuffled distribution was computed based on 1,000 iterations of random resampling of trials. In the results of recording session shown in Figure 4H (monkey D FOV 8), there were 1,200 choice trials, of which 173 were memory-error trials (including 34 “match trials” where the probability distributions—from which WM strength was computed—peaked around the actual response sequences). For each resampling, 139 trials (excluding the 34 match trials from the 173 memory-error trials) were randomly selected without replacement from the 1,200 choice trials, to determine the proportion of trials that belonged to low-strength mismatches by chance. We then computed the 99th percentile of the shuffled distribution as the chance-level threshold (dashed line in Fig. 4H).

### Neural activity-based baseline meta-WM analysis (Related to Figure 5)

#### Pupil size

The pupil size was filtered between 0.01 Hz and 10 Hz using a second-order Butterworth filter and then z-scored^60^.

#### Trial history

We tested the role of trial history in generating prior beliefs. We first obtained a quantitative description of reward per trial from the approximation of real reward: Monkey D received 3 water drops for correct responses under memory and forced-to-test conditions, 2 for correct offload trials, and 0 for error trials, following a 3-2-0 reward scheme; monkey Z was rewarded according to a 2-1-0 scheme. We could then describe trial history as the number of rewards received over a certain number of past trials.

We multiplied the trial history values with exponential history weights over time^61,62^, such that more recent trials would have greater impact on a current meta-WM judgment. We then summed the adjusted trial history to obtain the weighted mean history reward value. This value was then used to predict whether the current trial would be classified as a memory or offload trial. We optimized this decision discrimination ability - measured by AUROC - to determine the optimal mean (μ) of an exponential distribution. The parameter can be further utilized to generate historical weights.

#### Baseline meta-WM selectivity of each neuron

We refer to the decision selectivity during the baseline period as baseline meta-WM (prior) selectivity, since the results in Figure 5D indicated that baseline meta-WM exhibited variability at the trial level but not at the sequence level. After conducting linear regression, we obtained *β* (the regression coefficient), *r*^2^ (the coefficient of determination, representing regression explained variance), and *p* value for each neuron. Those neurons whose *p* value of linear regression less than 0.01 were considered baseline meta-WM (prior) selective neurons.

#### Neural decoder-based baseline meta-WM

The decoder architecture and training approach, as well as the decision boundary fitting, were exactly the same as those used for the meta-WM neural decoder described above except that the input was the baseline-averaged neuron activity instead of delay-averaged.

#### Linear regression of trial-level meta-WM

In this analysis, we utilized baseline meta-WM and (delay) WM strength as predictors to regress the (delay) meta-WM of each trial, where:

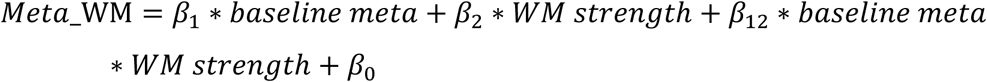

The regression coefficients were estimated using a resampling approach with replacement. For each resampling iteration, the number of samples matched the total number of trials. We used the resampled trials to conduct linear regression and to calculate the *β* coefficients and regression explained variance. This process was repeated 1,000 times to generate a distribution of *β* coefficients for statistical testing.

### Three task variables relationship analysis (Related to Figure 6)

#### Subspace estimation and principal angles between subspaces

Based on the previously decoded variables (baseline meta-WM, WM strength, and meta-WM), we performed linear regression to model these variables across trials using neuronal population activity. For example, we regressed meta-WM scores on neuronal population activity. To avoid overfitting, we employed Lasso regularization with 3-fold cross-validation in the linear regression. The beta coefficients obtained from these regressions defined an one-dimensional subspace (axis) for each variable, forming vectors within the neuronal state space. We then computed the angles between these one-dimensional subspaces^63^. As a control, we randomly split the trials into two halves to obtain separate estimations of each subspace and computed the angles between them; this process was repeated 112 times.

#### Variance accounted for (VAF) ratio between subspaces

For given two one-dimensional subspaces, which are vectors of beta coefficients,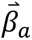 and 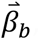 the VAF ratio^63,64^ for subspace pair (a, b) was defined as

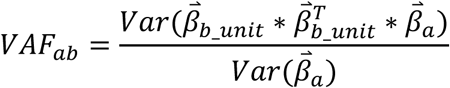

Where 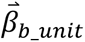 is the unit vector of 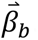. As a control, we randomly split the trials into two halves to obtain separate estimations of each subspace and computed the VAF ratio between them, and this process was repeated 112 times.

#### Time course of AUROC within subspaces

As previously described, for each variable, we derived a vector of regression coefficients 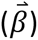 via linear regression to quantify the subspace. Taking the meta-WM subspace as an example, we projected the neuronal population activity matrix (F) at each time bin (33 ms) onto this vector to compute the estimated meta-WM scores (EMS):

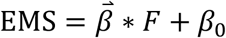

Here, 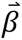 is a 1-by-n vector, where n denotes the number of neurons; *F* is an n-by-m matrix, with m representing the number of trials; and *β*_0_ is the intercept term from the linear regression model. This projection, from neuronal activity to the meta-WM subspace, yielded a scalar meta-WM score for each trial at each time bin. The same procedure was employed to estimate scores for the baseline meta and WM strength subspaces. Subsequently, we analyzed the time courses of AUROC to differentiate between memory and offload trials within both the baseline meta and meta-WM subspaces, as well as between low-strength and high-strength trials within the WM strength subspace.

#### Spatial organization of selective neurons

We quantified and compared the spatial distances within and between groups (baseline meta-WM, WM, meta-WM)^65^. We quantified the distance within groups by computing the average distance of each neuron to other neurons within the same group. We quantified the distance between groups by computing the average distance of each neuron to other neurons in different groups.

#### Subspace-selective neurons (Related to Fig. S17)

We randomly split the trials into two halves to obtain two separate estimations for each subspace and the corresponding neuronal coefficients, denoted as *β*_*A*_and *β*_*B*_ for a given example neuron. For each of the 112 resampling iterations, we computed the difference between the coefficients (*β*_*diff*_ = *β*_*A*_ − *β*_*B*_) and their mean (*β*_*mean*_ = (*β*_*A*_ + *β*_*B*_)/2). We then computed the mean difference between the coefficients across all resampling iterations (*β*_*diff*_*mean*_). A neuron was considered to contribute to the subspace in a given resampling if the absolute value of *β*_*mean*_exceeded five times the absolute value of *β*_*diff*_*mean*_. Neurons meeting this criterion in more than 99% of the resampling iterations were classified as subspace-selective, indicating a statistically significant and consistent contribution to the subspace.

---

**Fig. S1.**
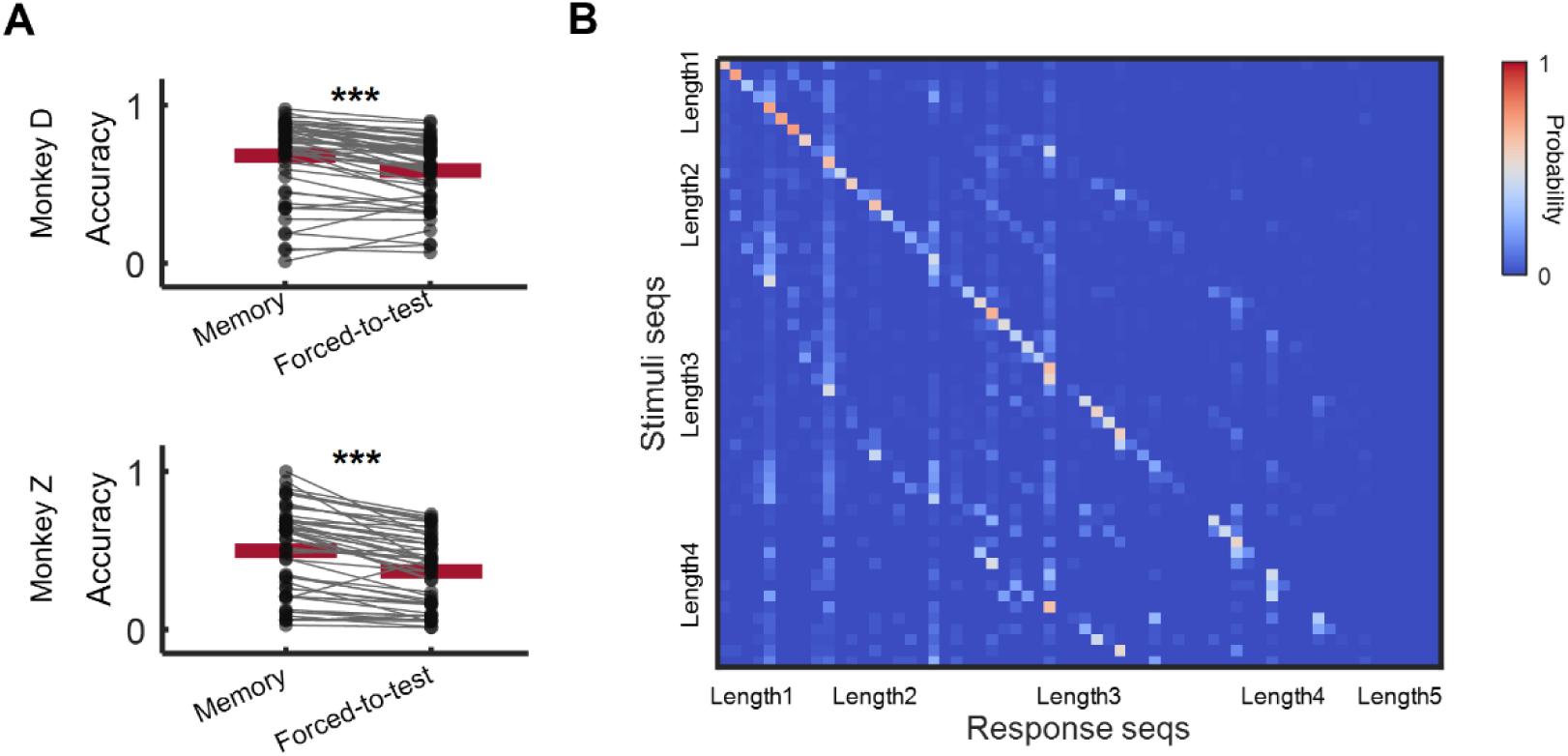
Additional behavior performance. (**A**) Within-sequence comparison of accuracy of memory trials and forced-to-test trials (two-tailed paired-sample t-test, p < 0.001 for both monkeys). Each dot represents one sequence, and red lines represent the mean accuracies across sequences. (**B**) Stimuli-to-response matrix of behavior data for monkey Z, including both correct trials (diagonal) and error trials (non-diagonal). Each element of the matrix denotes the probability for monkey Z to execute one particular sequence given the presentation of another specific sequence during the sample period.

**Fig. S2.**
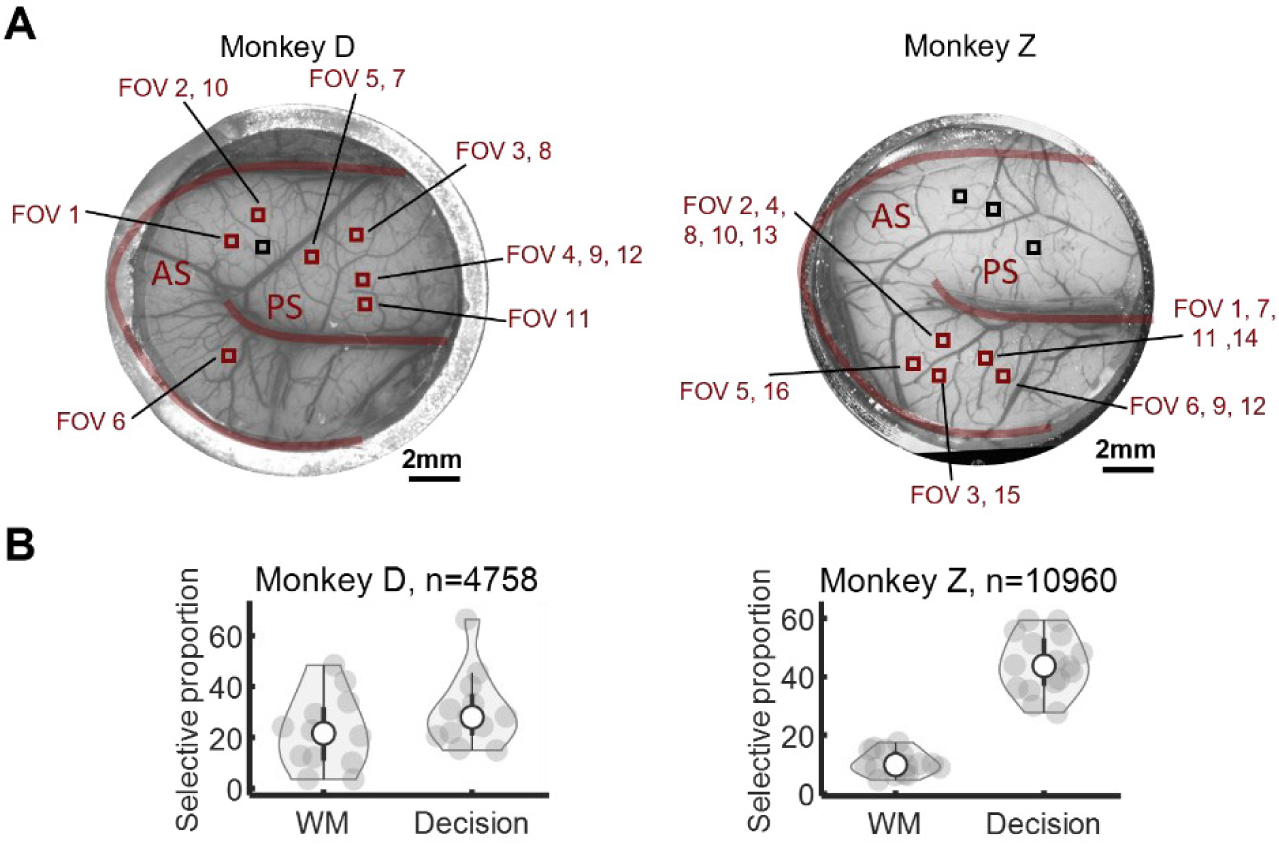
FOV distribution and task information-selective neuron proportions. (**A**) FOV distribution across imaging windows in both monkeys. Red squares denote FOVs utilized in data analysis, while black squares indicate those excluded from analysis due to poor GCaMP expression. (**B**) WM- and decision-selective neuron proportions in both monkey D and monkey Z. Each dot represents the proportion of neurons belonging to a selectivity category from one FOV. Overall, 22.3%/10.2% of neurons were WM-selective neurons and 28.6%/46.7% of neurons were decision-selective neurons in monkeys D and Z, respectively, after merging all FOVs.

**Fig. S3.**
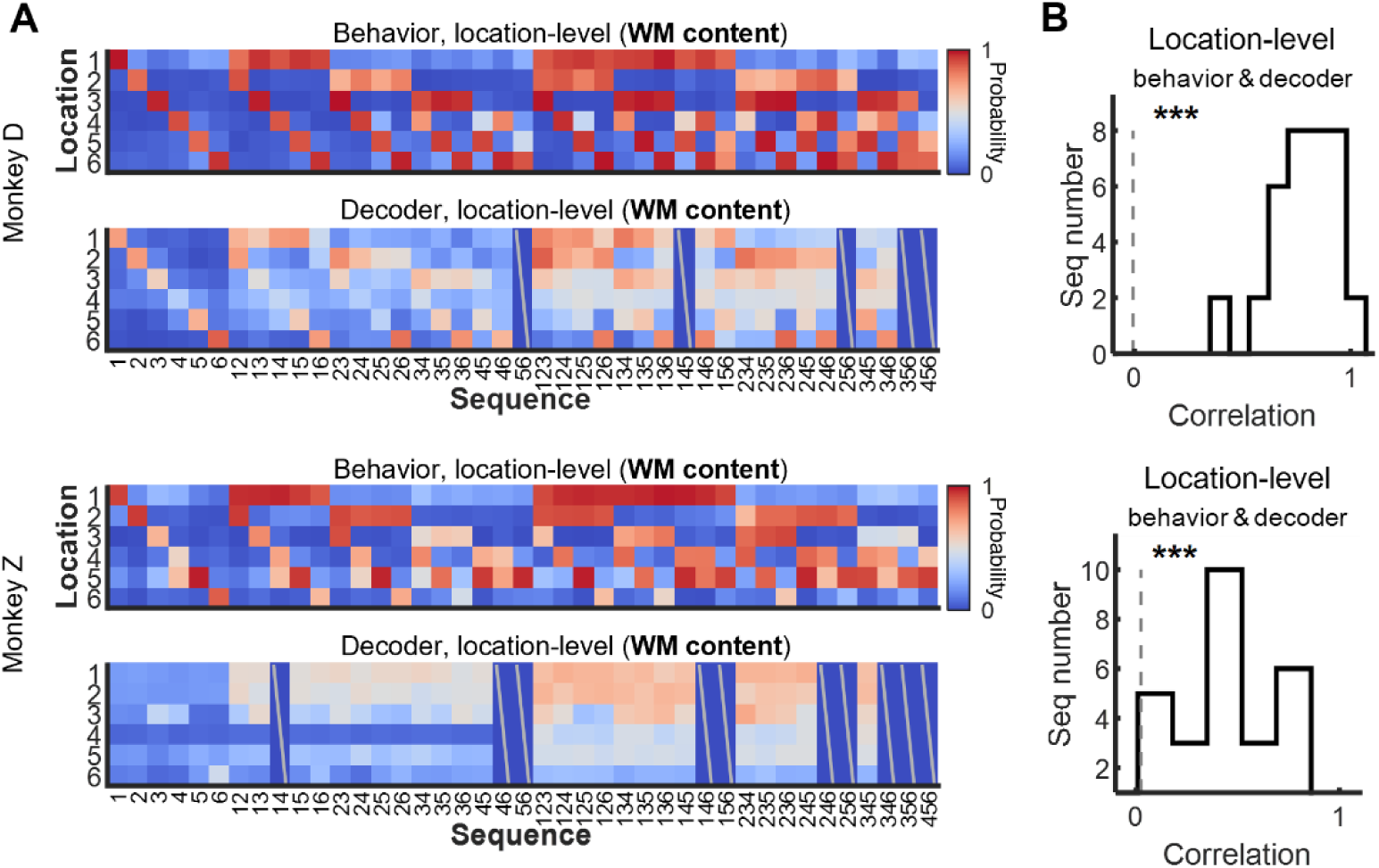
Pseudo-population results of neural WM location decoding for both monkeys. The pseudo-population results were obtained from resampled trials across sessions; see Methods subsection “Neural decoder-based WM strength”. For Monkey D, the pseudo-population results were derived from 10 FOVs encompassing 3,907 neurons. For Monkey Z, the results originated from 10 FOVs consisting of 7,215 neurons. The WM decoder was trained and tested on forced-to-test-correct and memory-correct trials, resampled from each FOV. (**A**) Illustration of location probability distribution of behavior data (based on accuracy) and the neuron decoder (based on pseudo-population data from 10 FOVs). (**B**) Histogram of Pearson correlation between location probability distributions of neural population decoder and behavior (A) (***: p < 0.001, two-tailed t-test against correlation value by chance).

**Fig. S4.**
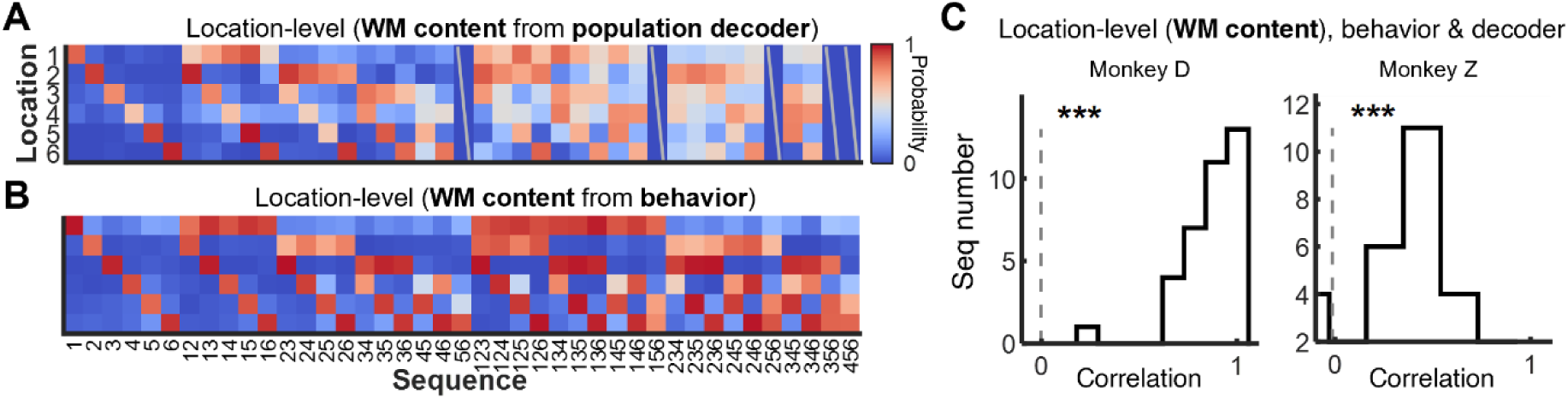
Additional analyses of stimulus location decoders-derived WM metrics. (**A**) Location probability distributions based on an example neural population decoder (monkey D, FOV 8), representing, for each sequence in the freely-choosing memory condition, the decoded probabilities of each location. (**B**) Behavioral location probability distributions (monkey D), representing, for each sequence in the forced-to-test condition, the response probabilities of each location based on task performance. (**C**) Histogram of Pearson correlation between location probability distributions of neural population decoder (A) and behavior (B) (***: p < 0.001, two-tailed t-test against correlation value by chance).

**Fig. S5.**
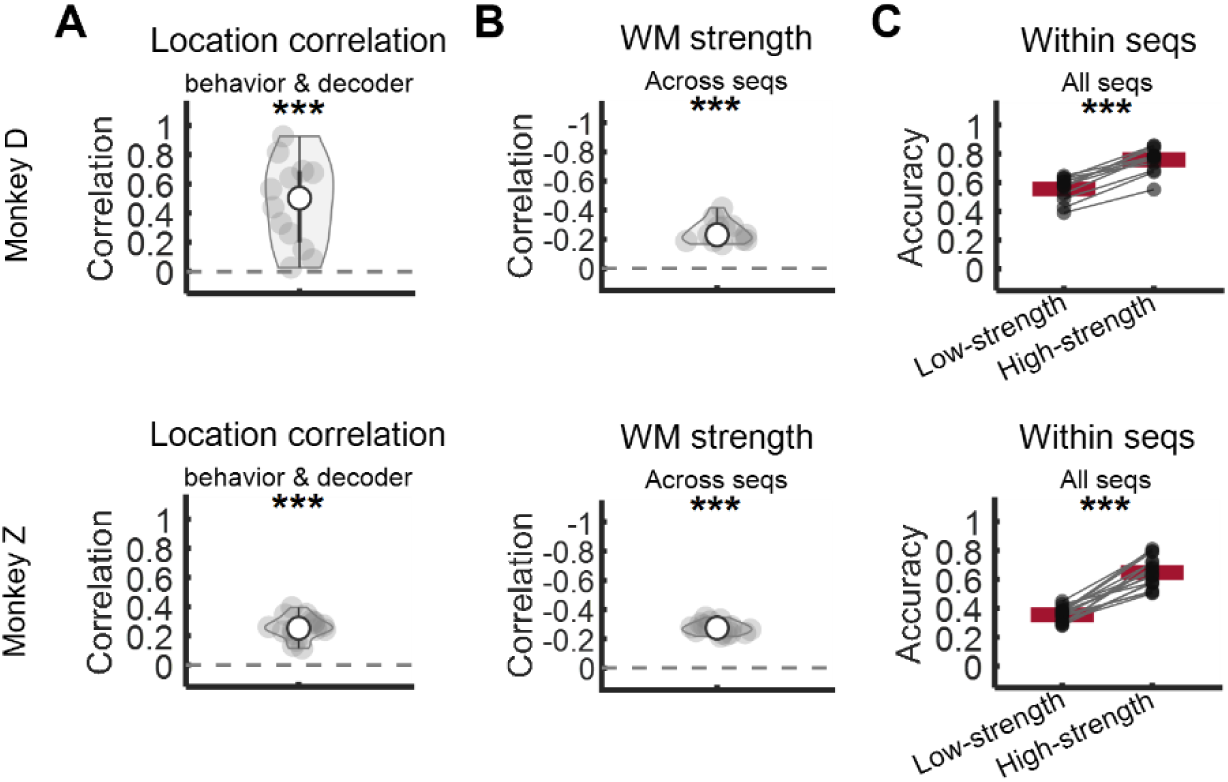
WM strength across FOVs and two monkeys. Each data point represents an averaged value from one FOV and red lines represent the mean across FOVs. (**A**) Pearson correlation of the location probability distribution between the population decoder (correct trials) and behavior. This correlation was compared to the chance level using a two-tailed one-sample t-test, with p < 0.001 for both monkeys. The chance level was computed as substituting behavior location probability distributions as random values sampled from a uniform probability distribution. In this analysis, we used forced-to-test-correct and memory-correct trials. (**B**) Pearson correlation between behavior recall variability and sequence-averaged WM strength, based on testing performance from forced-to-test (correct and error) trials. This correlation was compared to the chance level, with a two-tailed one-sample t-test showing p < 0.001 for both monkeys. (**C**) Within-sequence accuracy comparison between low WM strength and high WM strength trials using a two-tailed paired-sample t-test, showing p < 0.001 for both monkeys. The threshold dividing low and high WM strength trials was based on the median WM strength of all choice trials. Each dot represents averaged results across sequences from one session. WM strength was assessed from forced-to-test and memory (correct and error) trials.

**Fig. S6.**
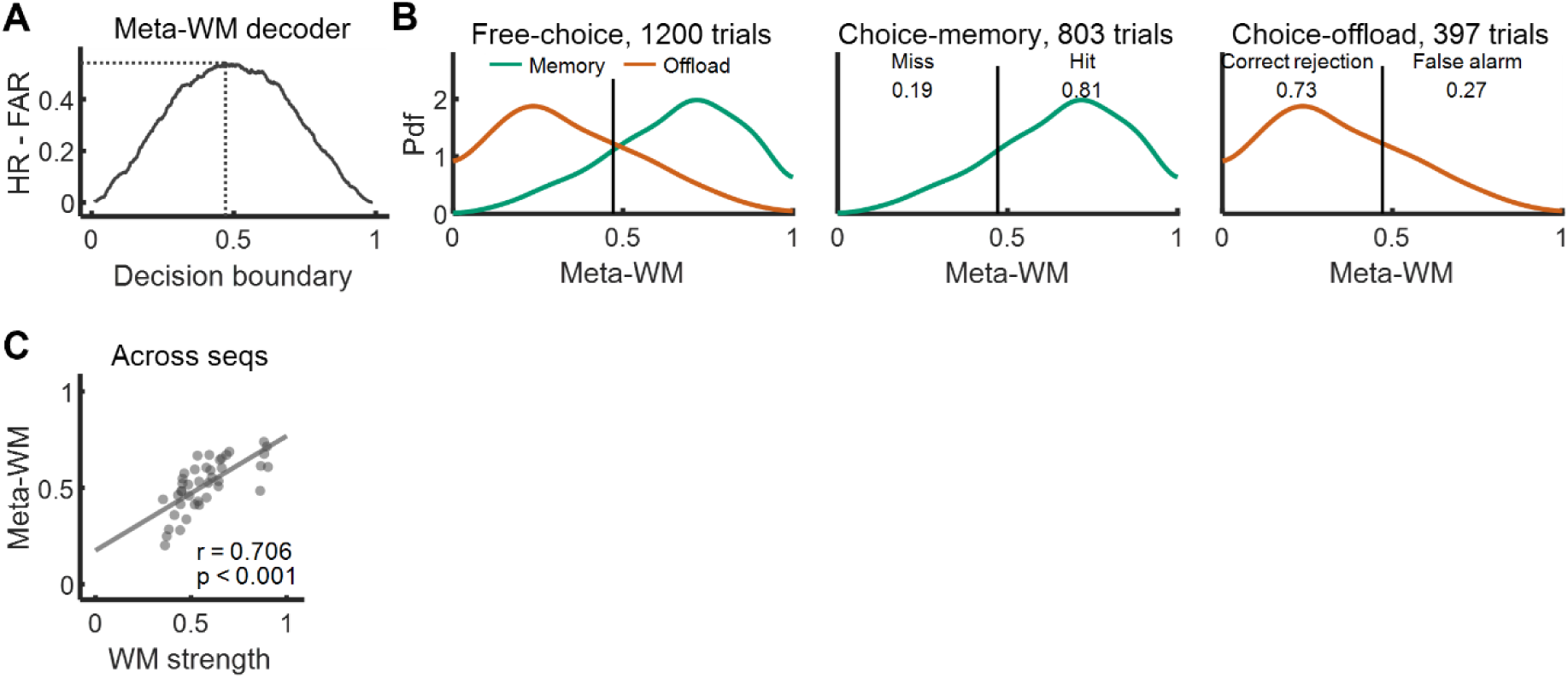
Meta-WM decoder performance details, including computation of the optimal decision boundary of the decoder. Results from FOV8 of monkey D. (**A**) The difference between the hit rate (HR) and the false alarm rate (FAR) for each decision boundary of the meta-WM decoder, ranging from 0.01 to 0.99 with a step of 0.001. The decoder reached its optimal value at 0.4710. (**B**) The trial distribution of meta-WM scores across all memory and offload trials, along with discrimination details of the meta-WM decoder (hit rate, miss rate, correct rejection rate, false alarm rate), given the optimal decision boundary identified in (A). (**C**) Pearson correlation between meta-WM and WM strength (both decoded from the neuron population) across sequences (r = 0.705, p < 0.001). Each dot represents a sequence, using all trials of the choice condition.

**Fig. S7.**
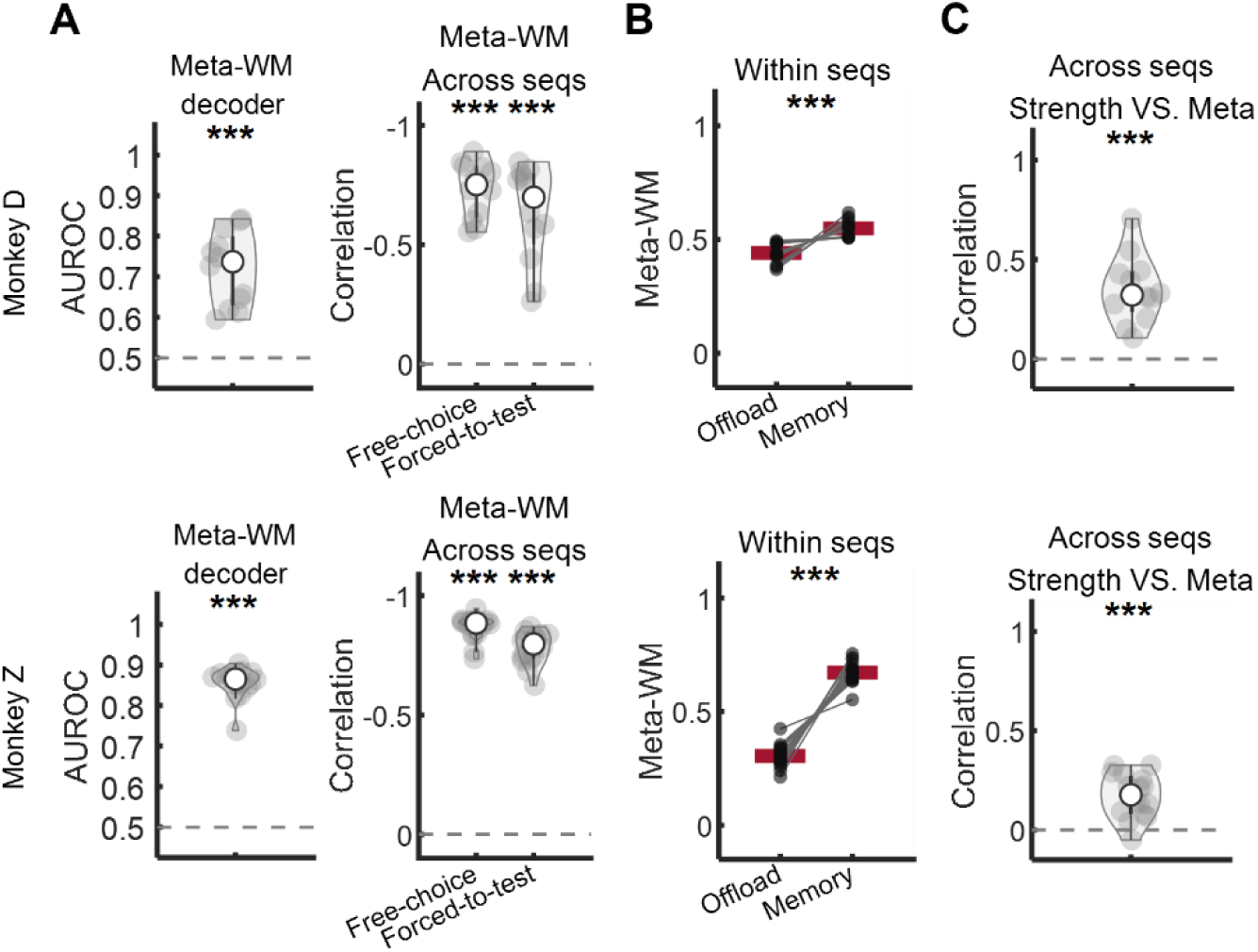
Meta-WM decoder performance across FOVs and two monkeys. Each data point represents an average value from a single session using choice (memory + offload) trials. ***: p < 0.001. (**A**) Performance of the meta-WM decoder. (Left) Comparison of AUROC with a chance level of 0.5 (using a two-tailed one-sample t-test, p < 0.001). (Right) Pearson correlation between the sequence-averaged behavioral offloading rate and the meta-WM scores of choice trials and forced-to-test trials, respectively. The correlation was compared to a chance level of 0 (two-tailed one-sample t-test, p < 0.001). (**B**) Within-sequence comparison of meta-WM scores between offload trials and memory trials (two-tailed paired-sample t-test, p < 0.001). Each dot represents the averaged results across sequences from a single session, with red lines indicating the mean. Paired dots originate from the same session. (**C**) Pearson correlation between meta-WM and WM strength across sequences, along with a comparison of this correlation to a chance level of 0 (two-tailed one-sample t-test, p < 0.001). Each dot represents the averaged result across sequences from a single session.

**Fig. S8.**
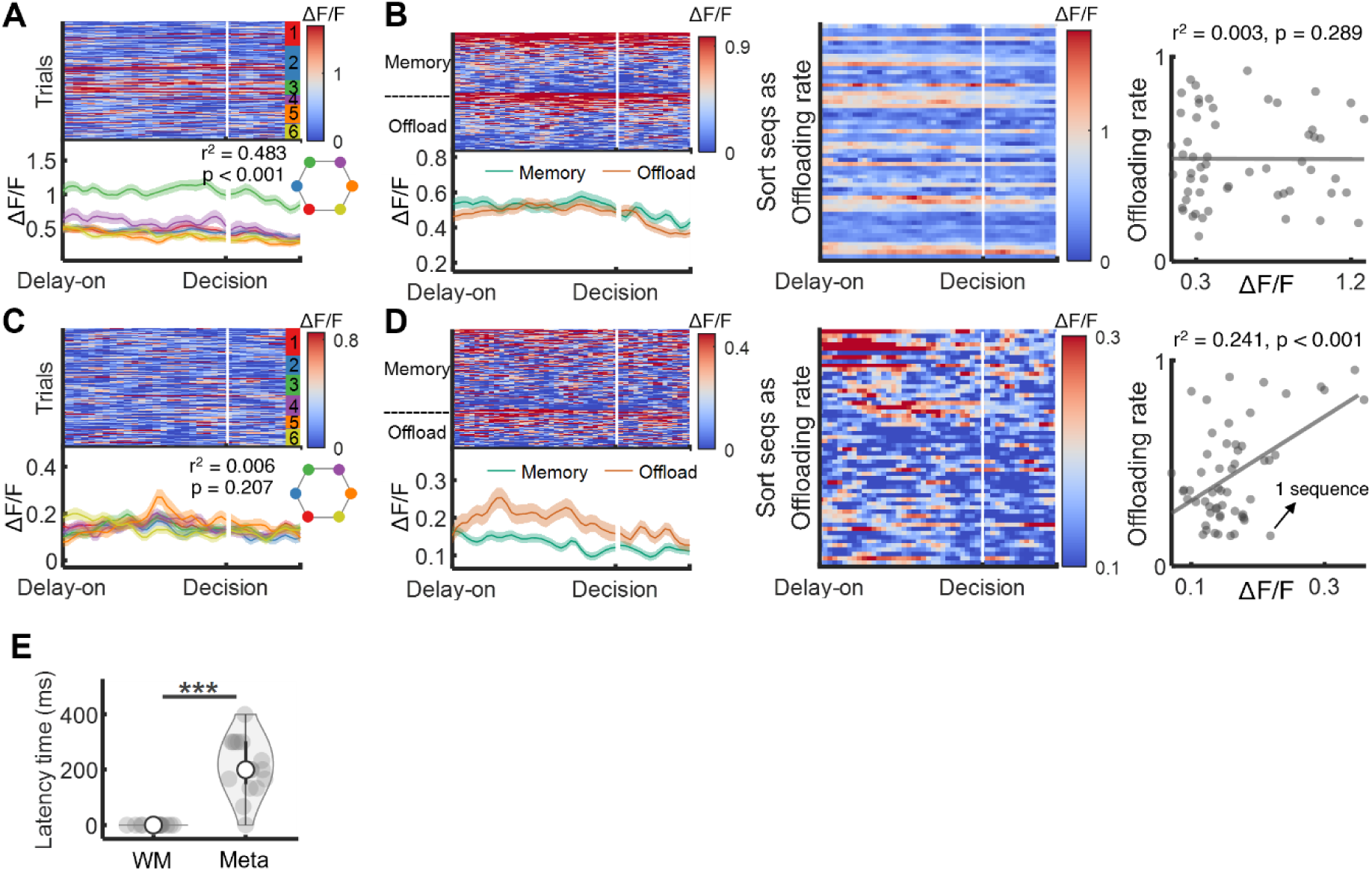
Pure selective neurons and latency time between WM strength and meta-WM representations. (**A-B**), Characteristics of a pure WM selective neuron. (A) Neuronal activity in response to different stimulus locations. The trials were grouped by whether any of the 6 specific locations was presented in a trial. (B) Neuron activity in memory and offload trials (left, choice trials were sorted as neuron activity descending), as well as in different sequences sorted by offloading rate (middle and right). Shaded areas represent SEMs. Each dot represents one sequence. (**C-D**), Same as (A-B), but of a pure meta-WM selective neuron. (**E**) Comparison of latency times between WM and meta-WM as in Fig. 4d. Resampled 16 times to compare latency times. Latency time of WM (after the last target of stimuli): mean = 0 ms, std < 0.001 ms. Latency time of meta-WM (after the last target of stimuli): mean = 212.287 ms, std = 102.367 ms. Each dot represents a result from one resampling, with a total of 16 resampling. Results from FOV8 of monkey D, using choice (memory + offload) trials.

**Fig. S9.**
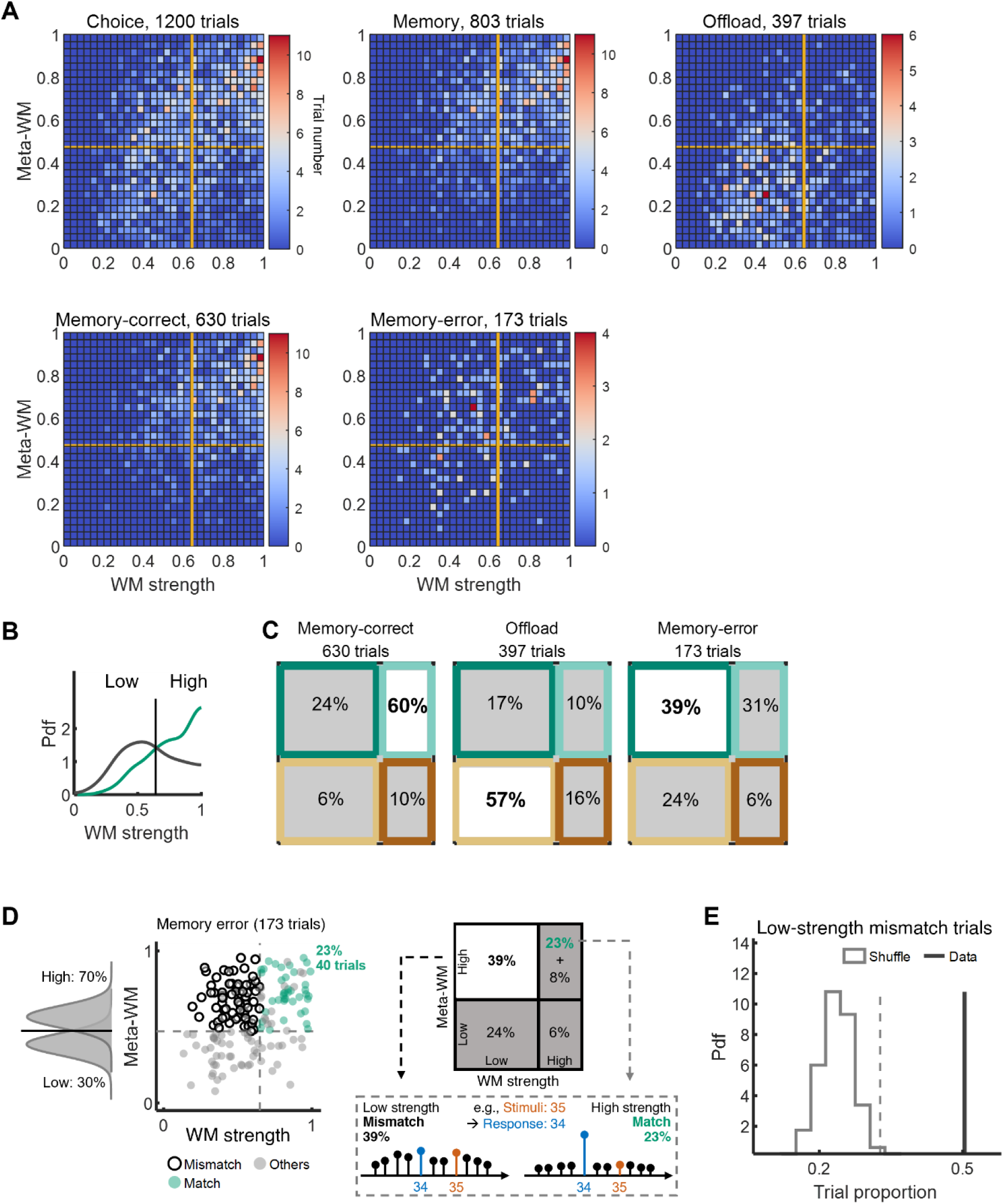
More additional results of relationship between WM strength and meta-WM using optimal boundaries. Results from an example FOV (FOV8 of monkey D) using choice (memory + offload) trials. The threshold dividing low meta-WM and high meta-WM was determined from the optimal decision boundary of the meta-WM decoder. Similarly, the threshold separating low WM strength from high WM strength was derived from the optimal boundary of WM strength, distinguishing between memory-correct and memory-error trials, akin to the criterion used in meta-WM. (**A**) Detailed two-dimensional trial distribution heatmaps of all conditions. (**B**) Probability density functions (Pdf) of the estimated WM strength for all memory-correct trials and all memory-error trials. The threshold (solid line) dividing low WM strength and high WM strength was derived from the optimal boundary. (**C**) Trial proportions of memory-correct trials, offload trials, and memory-error trials, categorized into four parts using the optimal boundaries. (**D**) (Top) Memory-error trials are categorized into four groups according to high or low levels of WM strength and meta-WM, determined by median values. Among error trials, 31% exhibited high WM strength and high meta-WM, including 23% match trials (where the decoding probability of response locations was the highest among all sequences) and 8% others. Meanwhile, 39% showed a mismatch with low WM strength and high meta-WM. (Bottom) Two example memory-error trials with sequence of stimuli [3-5] and response [3-4] illustrate match and mismatch. The high-strength match trial had concentrated probabilities responding as sequence [3-4], while the low-strength mismatch trial had widely distributed probabilities across many response sequences. (**E**) Test of trial proportion validity for low-strength mismatch trials in memory-error trials. Red line represents real data. Gray dashed line represents 99th percentile of a shuffle distribution.

**Fig. S10.**
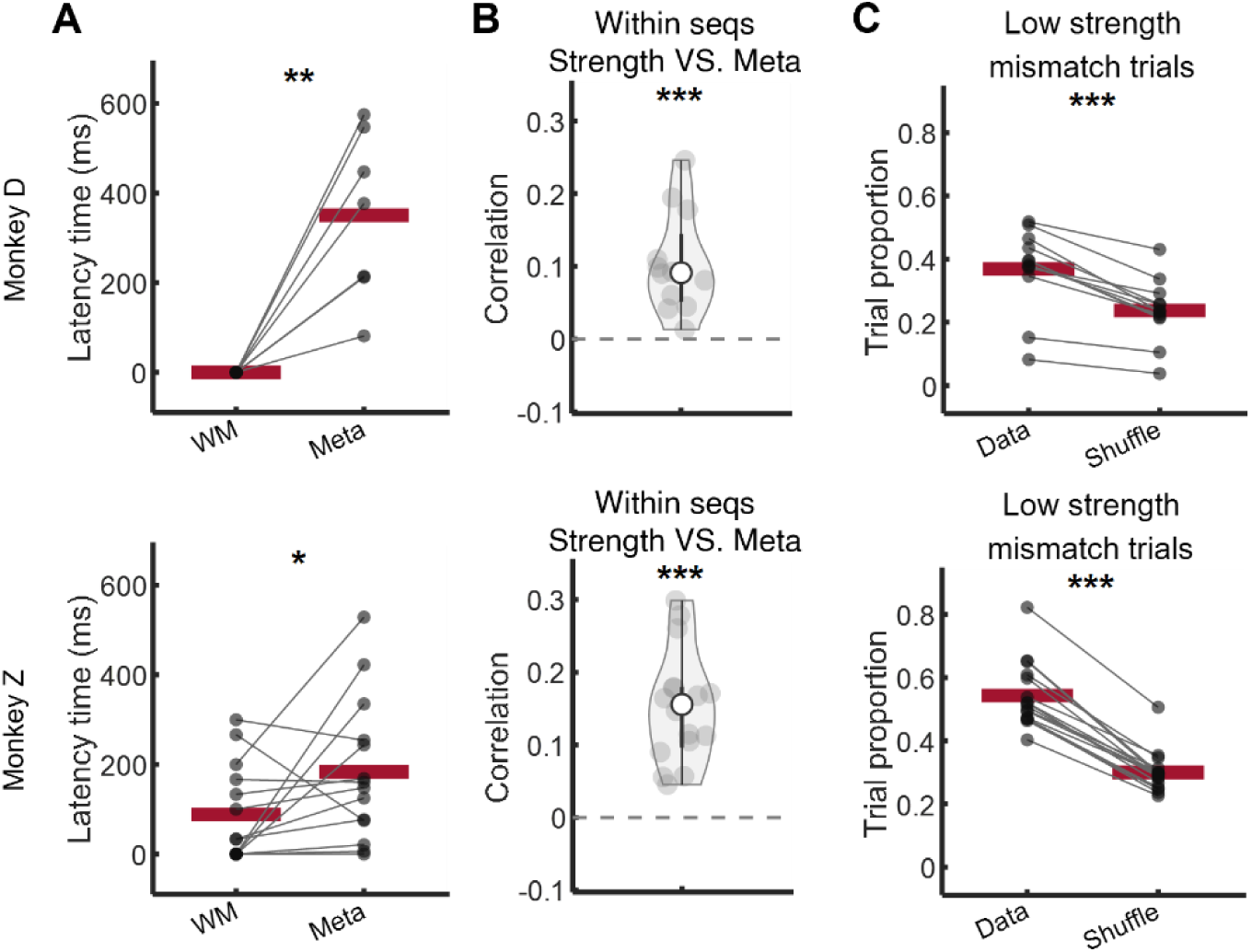
Relationship between WM strength and meta-WM across FOVs and two monkeys. In multi-FOV results here, each data point represents an averaged value from one FOV and red lines represent the means over FOVs. (**A**) Comparison of latency time between WM and meta-WM (single-tailed paired-sample t-test, for monkey D, p < 0.01; for monkey Z, p < 0.05). Time for each session was obtained in the same manner as in Fig. 4d. Latency time of WM (after last target of stimuli): for monkey D, mean = 0 ms, std < 0.001 ms; for monkey Z, mean = 92.764 ms, std = 114.000 ms. Latency time of meta-WM (after last target of stimuli): for monkey D, mean = 350.621 ms, std = 186.821 ms; for monkey Z, mean = 183.150 ms, std = 158.827 ms. Here, only FOVs whose latency time were not outliers were used in this analysis. The outliers were computed as the data points that more than three scaled MAD from the median, which is Matlab “isoutlier” default method. The scaled MAD is defined as c*median(abs(A-median(A))), where c=- 1/(sqrt(2)*erfcinv(3/2)) and A is input. (**B**) Pearson correlation between meta-WM and WM strength within each sequence. Overall mean correlation values are greater than chance for both monkeys (two-tailed one-sample t-test). Each dot represents averaged results across sequences from one session. (**C**) Comparison of trial proportion of low-strength mismatch trials in memory-error trials with shuffle level (two-tailed paired-sample t-test, for both monkeys). The ‘Data’ term represents proportions of low-strength mismatch trials in memory-error trials from real data. The ‘Shuffle’ term represents the results computed from a shuffled resampling (1,000 times of resampling; see methods subsection “Test of trial proportion validity of low-strength mismatch trials in memory-error trials”). *: p < 0.05; **: p < 0.01; ***: p < 0.001.

**Fig. S11.**
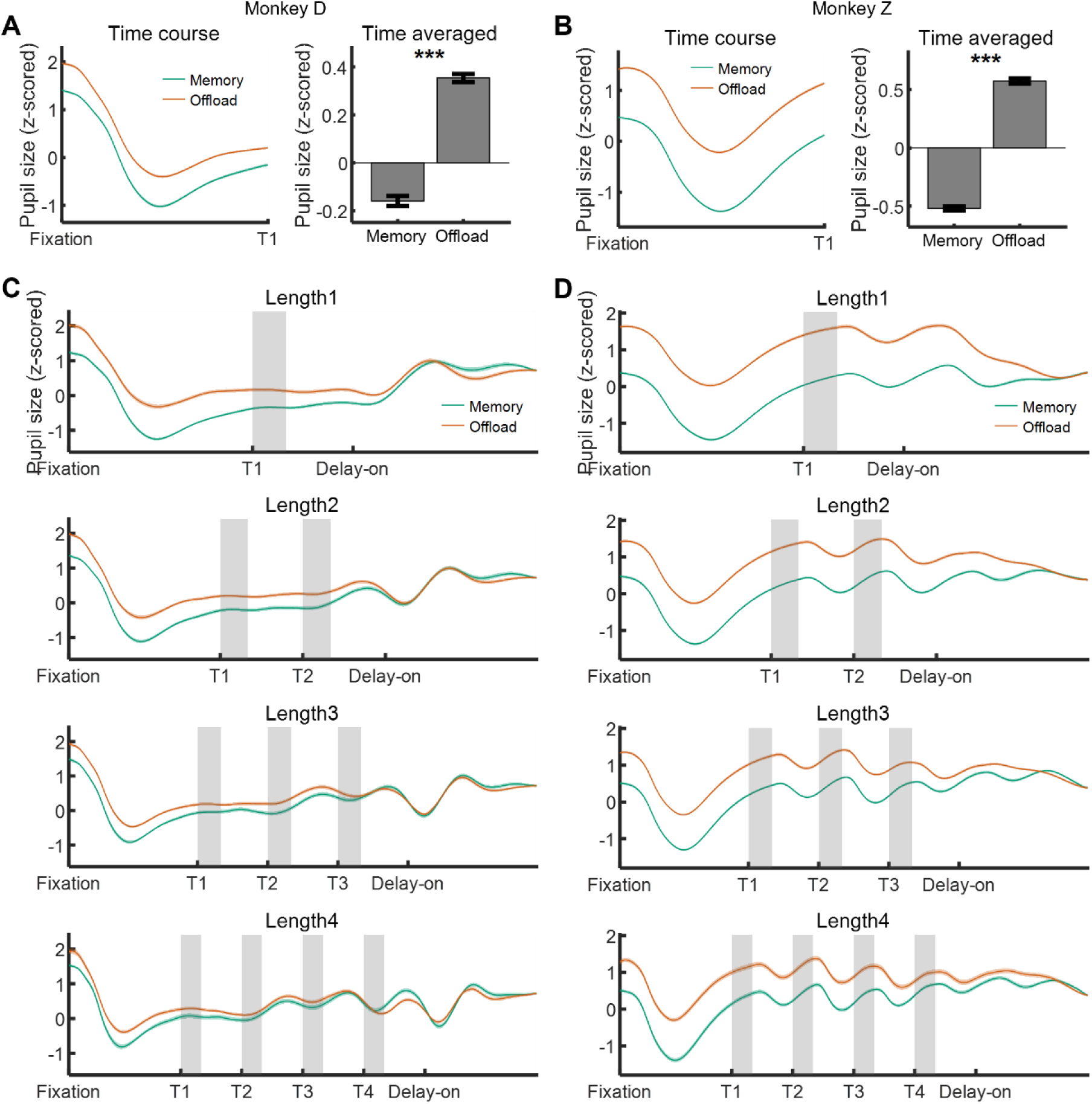
Pupil size details. (**A-B**) Pupil size during baseline period in monkey D (A) and monkey Z (B), averaged across all sessions and trials. (Left) The time course of pupil size during the baseline period of memory trials and offload trials. (Right) The time-averaged pupil size during the baseline period of memory trials and offload trials. The error bar was computed as SEM across sessions. (**C-D**) Pupil size in monkey D (C) and monkey Z (D) during the entire fixation time course in trials with different sequence lengths (1 to 4).

**Fig. S12.**
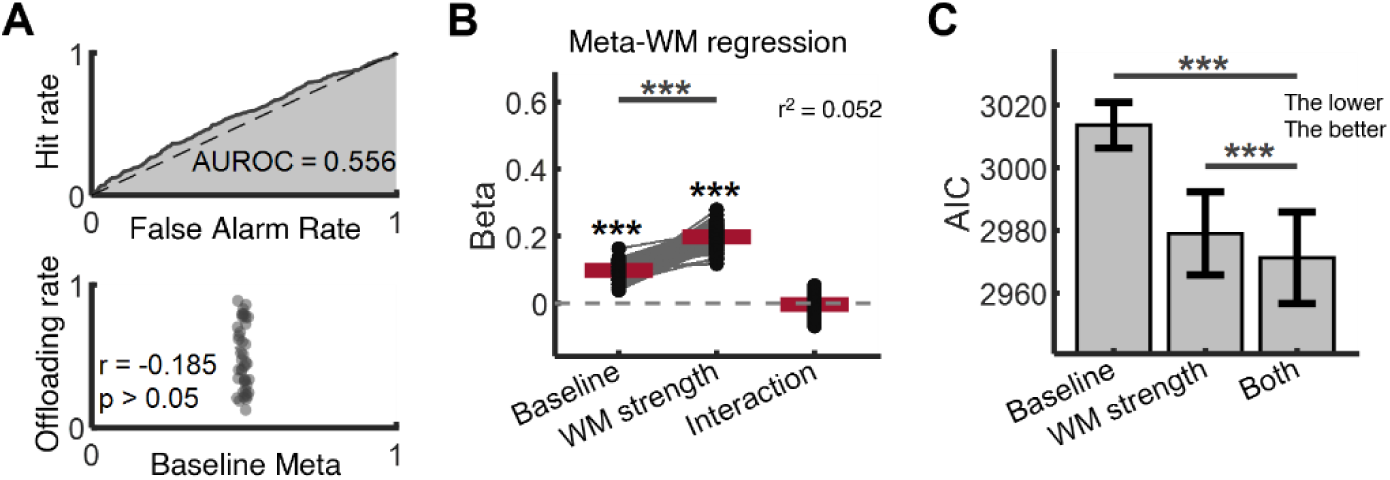
Prior belief results from monkey. **Z.** (**A**) Performance of the baseline meta-WM decoder. (Top) AUROC. (Bottom) Pearson correlation between offloading rates and sequence-averaged baseline meta-WM. Each dot represents one sequence. Results from FOV 13 of monkey Z using choice (memory + offload) trials. (**B**) Meta-WM linear regression coefficients of baseline and WM strength. While both baseline meta-WM and WM strength (measured during the delay) positively impacted meta-WM (single-tailed one-sample t-tests), WM strength was a stronger predictor of meta-WM overall (two-tailed paired-sample t-tests between coefficients). Each dot represents a coefficient from a random resampling-based regression model, with red lines representing mean values. Results from FOV 13 of monkey Z, using choice (memory + offload) trials. ***: p < 0.001. (**C**) Comparison of Akaike information criterion (AIC) in three meta-memory linear regression models (***: p < 0.001, two-tailed t-test). Models with lower AIC are considered to have higher performance, as AIC incorporates a penalty for the number of parameters. The input factors were baseline meta-WM, WM strength, and both, respectively. Error bars represent STDs. Results from FOV 13 of monkey Z using choice (memory + offload) trials.

**Fig. S13.**
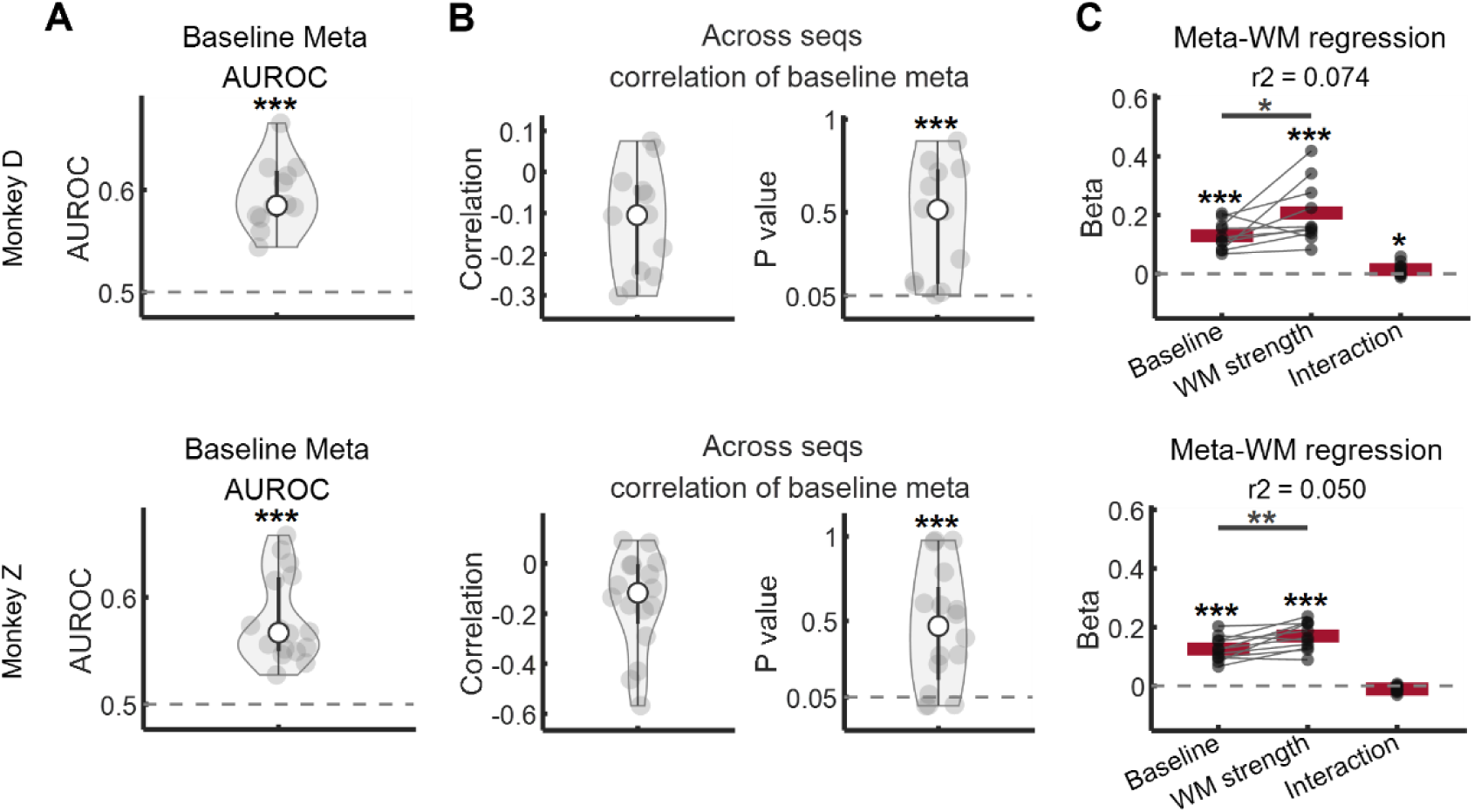
Prior belief results across FOVs and two monkeys. Multi-FOVs’ results. Each data point represents an averaged value from one session and red lines represent the mean. In these analyses, we used choice (memory + offload) trials. (**A-B**) Performance of the baseline meta-WM decoder. (A) Comparison of AUROC with the chance level of 0.5 (two-tailed one-sample t-test, p < 0.001). (B) Correlation between sequence-averaged baseline meta-WM and behavioral offloading rate across sequences. Comparison of p-values of the correlation with 0.05 (two-tailed one-sample t-test, p < 0.001). The dashed lines represent the chance level. (**C**) Meta-WM linear regression coefficients of baseline and WM strength. Comparison of each regression coefficient with the chance level of 0 (single-tailed one-sample t-test, p < 0.001 for both the baseline term and WM strength term for both monkeys; p > 0.05 for the interaction term for both monkeys). Comparison between the two coefficients (single-tailed paired-sample t-test, for monkey D, p < 0.05; for monkey Z, p < 0.01). Dashed lines represent chance level. In this analysis, only FOVs with regression coefficients that were not outliers and exceeded the 10th percentile were used. This analysis used all trials.

**Fig. S14.**
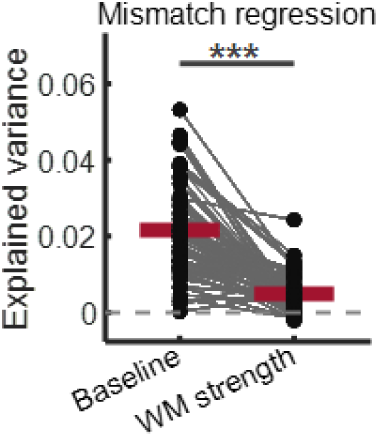
Prior belief results of mismatch regression. Explained variance (*r*^2^) of two linear regression was assessed to distinguish trial types (low-strength mismatch or offload) using baseline or WM strength, including only low-strength mismatch and offload trials. There is significant difference between two *r*^2^(two-tailed paired-sample t-test, p < 0.001). Each dot represents a result from one resampling, and red lines represent the mean across 1,000 random samples. Results from FOV 8 of monkey D.

**Fig. S15.**
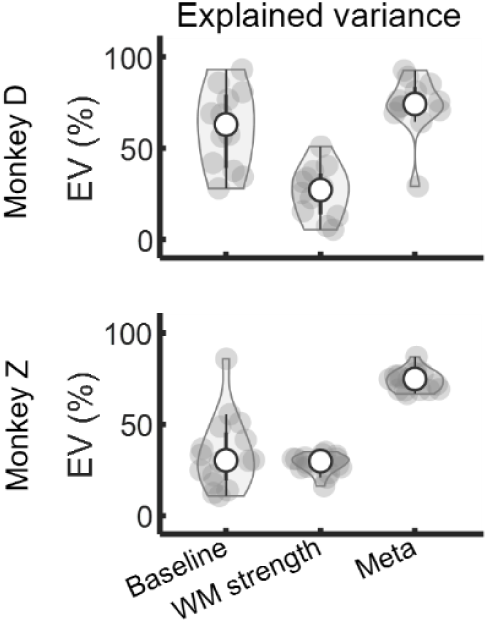
Explained variance of decoded variables in each subspace across FOVs and two monkeys. Explained variance of decoded variables (scores) across trials in three subspaces. Based on the previously decoded variables (baseline meta, WM strength, and meta-WM), we performed linear regression to model these variables across trials using neuronal population activity. For example, we regressed meta-WM scores on neuronal population activity. To prevent parameter overfitting, we employed Lasso regularization with 3-fold cross-validation in the linear regression. The beta coefficients obtained from these regressions defined a one-dimensional subspace for each variable, forming vectors within the neuronal state space. The explained variance presented here was calculated from these regressions. Each data point represents a result from one session.

**Fig. S16.**
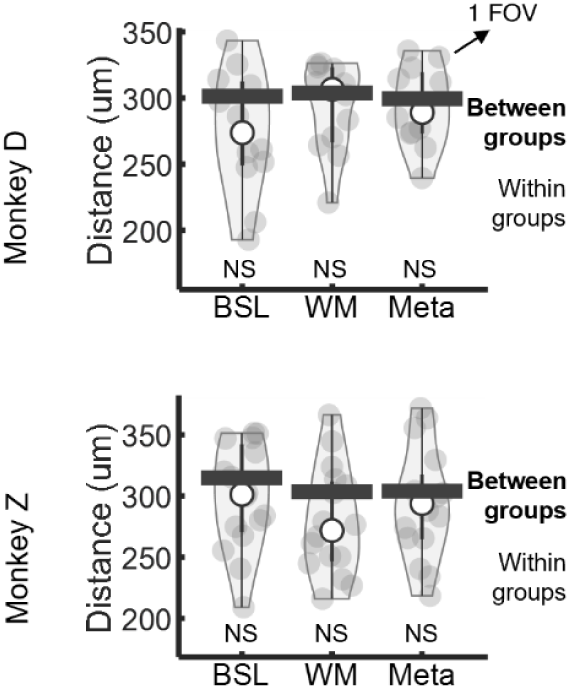
Spatial distances analysis across FOVs and two monkeys. Comparison of spatial distances within groups and between groups across FOVs (two-tailed t-test). Each gray dot represents the median distance of neurons within a single FOV, where each neuron’s distance was calculated as the average spatial distance between that neuron and others within the same group. The black bar represents a similar calculation as the gray dots but for results between different groups, and it shows the median value among FOVs. NS: non-significant. BSL: Baseline.

**Fig. S17.**
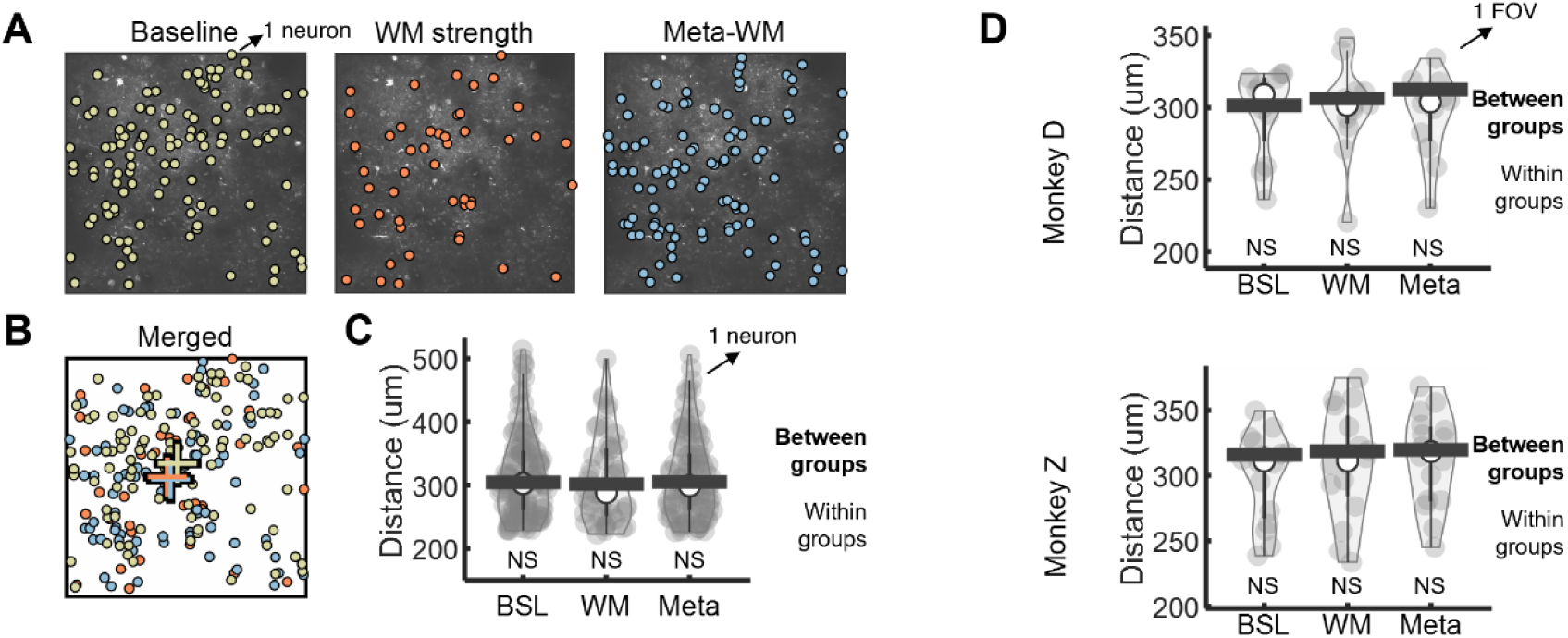
Spatial distances analysis of subspace-selective neurons. (**A-B**) (A) Spatial organization of subspace-selective neurons (see Methods subsection “Subspace-selective neurons”) for baseline meta, WM strength, and meta-WM (from FOV 4 of monkey D; 113, 48, 100 neurons, respectively). Each dot represents a neuron. In the merged panel (B), crosses represent spatial centroids. (**C**) Comparison of spatial distances within groups and between groups (two-tailed t-test; FOV 4 of monkey D). Each gray dot represents the average spatial distance between one neuron and others within the same group in (A). The black bar represents a calculation similar to that of the gray dots, but specifically for results between different groups, and it shows the median value among neurons. NS: non-significant. (**D**) Comparison of spatial distances within groups and between groups across FOVs (two-tailed t-test). Each gray dot represents the median distance of neurons within a single FOV, where each neuron’s distance was calculated as the average spatial distance between that neuron and others within the same group. The black bar represents a similar calculation as the gray dots but for results between different groups, and it shows the median value among FOVs. NS: non-significant. BSL: Baseline.

**Table S1.**
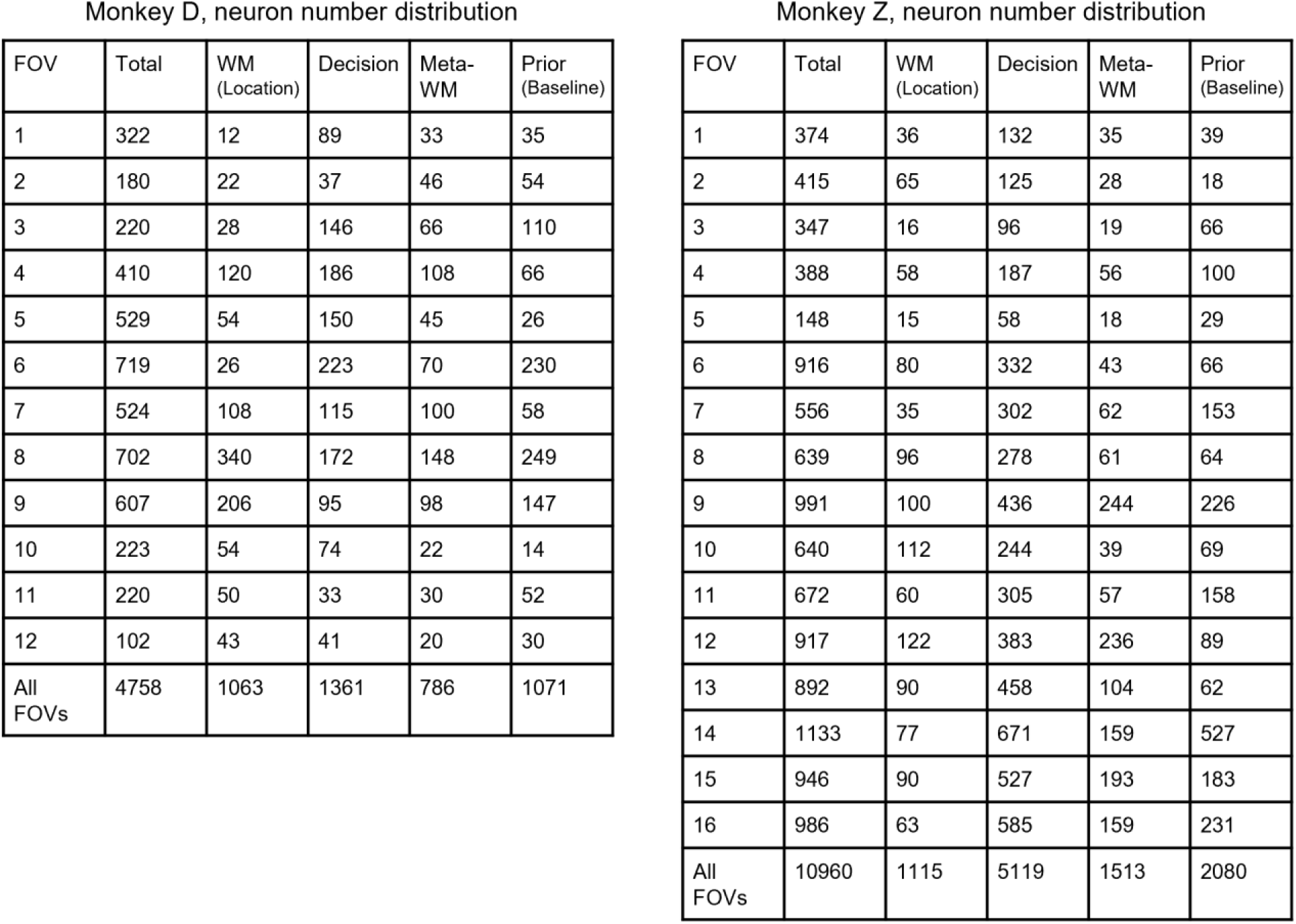
Distributions of neurons according to their task involvement across FOVs for both monkeys.

## References

1. Baddeley, A.D., and Hitch, G. (1974). Working Memory. In Psychology of Learning and Motivation, G.H. Bower, ed. (Academic Press), pp. 47–89. 10.1016/s0079-7421(08)60452-1.

2. Luck, S.J., and Vogel, E.K. (1997). The capacity of visual working memory for features and conjunctions. Nature 390, 279–281. 10.1038/36846.

3. Cowan, N. (2001). The magical number 4 in short-term memory: a reconsideration of mental storage capacity. Behav Brain Sci 24, 87–114; discussion 114–185. 10.1017/s0140525×01003922.

4. Risko, E.F., and Gilbert, S.J. (2016). Cognitive Offloading. Trends Cogn Sci 20, 676–688. 10.1016/j.tics.2016.07.002.

5. Flavell, J.H., and Wellman, H.M. (1975). Metamemory.

6. Yoo, A.H., Acerbi, L., and Ma, W.J. (2021). Uncertainty is maintained and used in working memory. J Vis 21, 13. 10.1167/jov.21.8.13.

7. Li, H.H., Sprague, T.C., Yoo, A.H., Ma, W.J., and Curtis, C.E. (2021). Joint representation of working memory and uncertainty in human cortex. Neuron 109, 3699–3712 e3696. 10.1016/j.neuron.2021.08.022.

8. Honig, M., Ma, W.J., and Fougnie, D. (2020). Humans incorporate trial-to-trial working memory uncertainty into rewarded decisions. Proc Natl Acad Sci U S A 117, 8391–8397. 10.1073/pnas.1918143117.

9. Xie, Y., Hu, P., Li, J., Chen, J., Song, W., Wang, X.J., Yang, T., Dehaene, S., Tang, S., Min, B., and Wang, L. (2022). Geometry of sequence working memory in macaque prefrontal cortex. Science 375, 632–639. 10.1126/science.abm0204.

10. Chen, J., Zhang, C., Hu, P., Min, B., and Wang, L. (2024). Flexible control of sequence working memory in the macaque frontal cortex. Neuron 112, 3502–3514 e3506. 10.1016/j.neuron.2024.07.024.

11. Tian, Z., Chen, J., Zhang, C., Min, B., Xu, B., and Wang, L. (2024). Mental programming of spatial sequences in working memory in the macaque frontal cortex. Science 385, eadp6091. 10.1126/science.adp6091.

12. Fleming, S.M., and Dolan, R.J. (2012). The neural basis of metacognitive ability. Philos Trans R Soc Lond B Biol Sci 367, 1338–1349. 10.1098/rstb.2011.0417.

13. Vaccaro, A.G., and Fleming, S.M. (2018). Thinking about thinking: A coordinate-based meta-analysis of neuroimaging studies of metacognitive judgements. Brain Neurosci Adv 2, 2398212818810591. 10.1177/2398212818810591.

14. Miyamoto, K., Trudel, N., Kamermans, K., Lim, M.C., Lazari, A., Verhagen, L., Wittmann, M.K., and Rushworth, M.F.S. (2021). Identification and disruption of a neural mechanism for accumulating prospective metacognitive information prior to decision-making. Neuron 109, 1396–1408 e1397. 10.1016/j.neuron.2021.02.024.

15. Middlebrooks, P.G., and Sommer, M.A. (2012). Neuronal correlates of metacognition in primate frontal cortex. Neuron 75, 517–530. 10.1016/j.neuron.2012.05.028.

16. Miyamoto, K., Osada, T., Setsuie, R., Takeda, M., Tamura, K., Adachi, Y., and Miyashita, Y. (2017). Causal neural network of metamemory for retrospection in primates. Science 355, 188–193. 10.1126/science.aal0162.

17. Kepecs, A., Uchida, N., Zariwala, H.A., and Mainen, Z.F. (2008). Neural correlates, computation and behavioural impact of decision confidence. Nature 455, 227–231. 10.1038/nature07200.

18. Joo, H.R., Liang, H., Chung, J.E., Geaghan-Breiner, C., Fan, J.L., Nachman, B.P., Kepecs, A., and Frank, L.M. (2021). Rats use memory confidence to guide decisions. Curr Biol 31, 4571–4583 e4574. 10.1016/j.cub.2021.08.013.

19. Baird, B., Smallwood, J., Gorgolewski, K.J., and Margulies, D.S. (2013). Medial and lateral networks in anterior prefrontal cortex support metacognitive ability for memory and perception. J Neurosci 33, 16657–16665. 10.1523/JNEUROSCI.0786-13.2013.

20. Kwok, S.C., Cai, Y., and Buckley, M.J. (2019). Mnemonic Introspection in Macaques Is Dependent on Superior Dorsolateral Prefrontal Cortex But Not Orbitofrontal Cortex. J Neurosci 39, 5922–5934. 10.1523/JNEUROSCI.0330-19.2019.

21. Fleming, S.M. (2024). Metacognition and Confidence: A Review and Synthesis. Annu Rev Psychol 75, 241–268. 10.1146/annurev-psych-022423-032425.

22. Lau, H., Michel, M., LeDoux, J.E., and Fleming, S.M. (2022). The mnemonic basis of subjective experience. Nature Reviews Psychology 1, 479–488. 10.1038/s44159-022-00068-6.

23. Kiani, R., and Shadlen, M.N. (2009). Representation of confidence associated with a decision by neurons in the parietal cortex. Science 324, 759–764. 10.1126/science.1169405.

24. Okazawa, G., Hatch, C.E., Mancoo, A., Machens, C.K., and Kiani, R. (2021). Representational geometry of perceptual decisions in the monkey parietal cortex. Cell 184, 3748–3761 e3718. 10.1016/j.cell.2021.05.022.

25. Alter, A.L., and Oppenheimer, D.M. (2009). Uniting the tribes of fluency to form a metacognitive nation. Pers Soc Psychol Rev 13, 219–235. 10.1177/1088868309341564.

26. Koriat, A., Bjork, R.A., Sheffer, L., and Bar, S.K. (2004). Predicting one’s own forgetting: the role of experience-based and theory-based processes. J Exp Psychol Gen 133, 643–656. 10.1037/0096-3445.133.4.643.

27. Hu, X., Zheng, J., Su, N., Fan, T., Yang, C., Yin, Y., Fleming, S.M., and Luo, L. (2021). A Bayesian inference model for metamemory. Psychol Rev 128, 824–855. 10.1037/rev0000270.

28. Fiacconi, C.M., Peter, E.L., Owais, S., and Kohler, S. (2016). Knowing by heart: Visceral feedback shapes recognition memory judgments. J Exp Psychol Gen 145, 559–572. 10.1037/xge0000164.

29. Ferrigno, S., Kornell, N., and Cantlon, J.F. (2017). A metacognitive illusion in monkeys. Proc Biol Sci 284. 10.1098/rspb.2017.1541.

30. Rahnev, D., and Denison, R.N. (2018). Suboptimality in perceptual decision making. Behav Brain Sci 41, e223. 10.1017/S0140525X18000936.

31. Gavas, R.D., Tripathy, S.R., Chatterjee, D., and Sinha, A. (2018). Cognitive load and metacognitive confidence extraction from pupillary response. Cognitive Systems Research 52, 325–334. 10.1016/j.cogsys.2018.07.021.

32. Lempert, K.M., Chen, Y.L., and Fleming, S.M. (2015). Relating Pupil Dilation and Metacognitive Confidence during Auditory Decision-Making. PLoS One 10, e0126588. 10.1371/journal.pone.0126588.

33. Purcell, B.A., and Kiani, R. (2016). Hierarchical decision processes that operate over distinct timescales underlie choice and changes in strategy. Proc Natl Acad Sci U S A 113, E4531–4540. 10.1073/pnas.1524685113.

34. Rouault, M., and Fleming, S.M. (2020). Formation of global self-beliefs in the human brain. Proc Natl Acad Sci U S A 117, 27268–27276. 10.1073/pnas.2003094117.

35. Hattori, R., Danskin, B., Babic, Z., Mlynaryk, N., and Komiyama, T. (2019). Area-Specificity and Plasticity of History-Dependent Value Coding During Learning. Cell 177, 1858–1872 e1815. 10.1016/j.cell.2019.04.027.

36. Stuber, G.D., Steinmetz, N.A., Bowen, A.J., Hjort, M.M., and Ottenheimer, D.J. (2023). A stable, distributed code for cue value in mouse cortex during reward learning. eLife 12. 10.7554/eLife.84604.3.

37. Li, N., Daie, K., Svoboda, K., and Druckmann, S. (2016). Robust neuronal dynamics in premotor cortex during motor planning. Nature 532, 459–464. 10.1038/nature17643.

38. Sun, Y., Nitz, D.A., Xu, X., and Giocomo, L.M. (2024). Subicular neurons encode concave and convex geometries. Nature 627, 821–829. 10.1038/s41586-024-07139-z.

39. Meyniel, F., Schlunegger, D., and Dehaene, S. (2015). The Sense of Confidence during Probabilistic Learning: A Normative Account. PLoS Comput Biol 11, e1004305. 10.1371/journal.pcbi.1004305.

40. Bang, D., and Fleming, S.M. (2018). Distinct encoding of decision confidence in human medial prefrontal cortex. Proc Natl Acad Sci U S A 115, 6082–6087. 10.1073/pnas.1800795115.

41. Walker, E.Y., Cotton, R.J., Ma, W.J., and Tolias, A.S. (2020). A neural basis of probabilistic computation in visual cortex. Nat Neurosci 23, 122–129. 10.1038/s41593-019-0554-5.

42. Geurts, L.S., Cooke, J.R.H., van Bergen, R.S., and Jehee, J.F.M. (2022). Subjective confidence reflects representation of Bayesian probability in cortex. Nat Hum Behav 6, 294–305. 10.1038/s41562-021-01247-w.

43. Walker, E.Y., Pohl, S., Denison, R.N., Barack, D.L., Lee, J., Block, N., Ma, W.J., and Meyniel, F. (2023). Studying the neural representations of uncertainty. Nat Neurosci 26, 1857–1867. 10.1038/s41593-023-01444-y.

44. Lebreton, M., Abitbol, R., Daunizeau, J., and Pessiglione, M. (2015). Automatic integration of confidence in the brain valuation signal. Nat Neurosci 18, 1159–1167. 10.1038/nn.4064.

45. Rutishauser, U., Aflalo, T., Rosario, E.R., Pouratian, N., and Andersen, R.A. (2018). Single-Neuron Representation of Memory Strength and Recognition Confidence in Left Human Posterior Parietal Cortex. Neuron 97, 209–220 e203. 10.1016/j.neuron.2017.11.029.

46. Komura, Y., Nikkuni, A., Hirashima, N., Uetake, T., and Miyamoto, A. (2013). Responses of pulvinar neurons reflect a subject’s confidence in visual categorization. Nat Neurosci 16, 749–755. 10.1038/nn.3393.

47. Rutishauser, U., Ye, S., Koroma, M., Tudusciuc, O., Ross, I.B., Chung, J.M., and Mamelak, A.N. (2015). Representation of retrieval confidence by single neurons in the human medial temporal lobe. Nat Neurosci 18, 1041–1050. 10.1038/nn.4041.

48. Vivar-Lazo, M., and Fetsch, C.R. (2024). Neural basis of concurrent deliberation toward a choice and degree of confidence. bioRxiv. 10.1101/2024.08.06.606833.

49. Rigotti, M., Barak, O., Warden, M.R., Wang, X.J., Daw, N.D., Miller, E.K., and Fusi, S. (2013). The importance of mixed selectivity in complex cognitive tasks. Nature 497, 585–590. 10.1038/nature12160.

50. Jazayeri, M., and Ostojic, S. (2021). Interpreting neural computations by examining intrinsic and embedding dimensionality of neural activity. Curr Opin Neurobiol 70, 113–120. 10.1016/j.conb.2021.08.002.

51. Beiran, M., Dubreuil, A., Valente, A., Mastrogiuseppe, F., and Ostojic, S. (2021). Shaping Dynamics With Multiple Populations in Low-Rank Recurrent Networks. Neural Comput 33, 1572–1615. 10.1162/neco_a_01381.

## Methods references

54. Brainard, D.H. (1997). The Psychophysics Toolbox. Spat Vis 10, 433–436.

55. Pelli, D.G. (1997). The VideoToolbox software for visual psychophysics: transforming numbers into movies. Spat Vis 10, 437–442. 10.1163/156856897X00366.

56. Petrides, M., Tomaiuolo, F., Yeterian, E.H., and Pandya, D.N. (2012). The prefrontal cortex: comparative architectonic organization in the human and the macaque monkey brains. Cortex 48, 46–57. 10.1016/j.cortex.2011.07.002.

57. Sadakane, O., Masamizu, Y., Watakabe, A., Terada, S., Ohtsuka, M., Takaji, M., Mizukami, H., Ozawa, K., Kawasaki, H., Matsuzaki, M., and Yamamori, T. (2015). Long-Term Two-Photon Calcium Imaging of Neuronal Populations with Subcellular Resolution in Adult Non-human Primates. Cell Rep 13, 1989–1999. 10.1016/j.celrep.2015.10.050.

58. Pnevmatikakis, E.A., and Giovannucci, A. (2017). NoRMCorre: An online algorithm for piecewise rigid motion correction of calcium imaging data. J Neurosci Methods 291, 83–94. 10.1016/j.jneumeth.2017.07.031.

59. Pachitariu, M., Stringer, C., Dipoppa, M., Schröder, S., Rossi, L.F., Dalgleish, H., Carandini, M., and Harris, K.D. (2017). Suite2p: beyond 10,000 neurons with standard two-photon microscopy. 10.1101/061507.

60. Osorio, S., Irani, M., Herrada, J., and Aboitiz, F. (2022). Neural responses to sensory novelty with and without conscious access. Neuroimage 262, 119516. 10.1016/j.neuroimage.2022.119516.

61. Hattori, R., Danskin, B., Babic, Z., Mlynaryk, N., and Komiyama, T. (2019). Area-Specificity and Plasticity of History-Dependent Value Coding During Learning. Cell 177, 1858–1872 e1815. 10.1016/j.cell.2019.04.027.

62. Stuber, G.D., Steinmetz, N.A., Bowen, A.J., Hjort, M.M., and Ottenheimer, D.J. (2023). A stable, distributed code for cue value in mouse cortex during reward learning. eLife 12. 10.7554/eLife.84604.3.

63. Xie, Y., Hu, P., Li, J., Chen, J., Song, W., Wang, X.J., Yang, T., Dehaene, S., Tang, S., Min, B., and Wang, L. (2022). Geometry of sequence working memory in macaque prefrontal cortex. Science 375, 632–639. 10.1126/science.abm0204.

64. Tian, Z., Chen, J., Zhang, C., Min, B., Xu, B., and Wang, L. (2024). Mental programming of spatial sequences in working memory in the macaque frontal cortex. Science 385, eadp6091. 10.1126/science.adp6091.

65. Sun, Y., Nitz, D.A., Xu, X., and Giocomo, L.M. (2024). Subicular neurons encode concave and convex geometries. Nature 627, 821–829. 10.1038/s41586-024-07139-z.

