## Supplementary material for "Multiple Routes to Metacognitive Judgments of Working Memory in the Macaque Prefrontal Cortex": Supp

We computed *entropy* in a classical manner, using the distribution of all possible response probabilities of sequences, where:

$$Entropy(stimuli\ seq) = -\sum_{response\ seq} P(response\ seq \mid stimuli\ seq) * \log\big(P(response\ seq \mid stimuli\ seq)\big)$$

$$Entropy \in [0,\ Max]$$

Here, we also compute the theoretical upper bound of *entropy* by calculating the entropy of a uniform distribution, denoted as *Max*. Finally, we determined the recall variability by normalizing the entropy from [0, Max] to [0, 1], where:

$$Recall\ variability(stimuli\ seq) = \frac{Entropy(stimuli\ seq)}{Max} \in [0,\ 1]$$

In this context, a recall variability of 1 represents a uniform probability distribution (which would be noisy WM), and a recall variability of 0 indicates that the probability of one specific sequence is 1 (representing very high precision of WM).

For example, in Figure 1C, responses to the sequence [1-3] of monkey D were mostly concentrated around its corresponding physical stimulus, indicating low variability. In contrast, responses to the sequence [2-5-6] were widely distributed, reflecting high variability.

***WM selectivity of each neuron***

We conducted linear regression for each neuron to quantify its tuning properties as follows:

$$y = X * \beta + \beta_0$$

For the regression of location WM, $y$ represented the neuronal activity for each trial (a column vector with 1 by number of trials), $\beta_0$ was the intercept term, $\beta$ was the regression coefficient to be fitted (a 1-by-6 vector, indicating weights of 6 locations), and $X$ represented the stimulus sequence for each trial (a design matrix with 6 by number of trials, with a one-hot-like 1-by-6 vector for each trial, e.g. stimulus sequence [3-5] would be [0 0 1 0 1 0].). Only correct trials (in both the memory and forced-to-test conditions) were used in the regression of location WM.

After conducting linear regression, we obtained $\beta$, $r^2$(the coefficient of determination, representing regression explained variance), and $p$ value of regression (representing the overall effectiveness of location $\beta$s to predict neural activity) for each neuron. Those neurons whose $p$ values of linear regression were less than 0.01 would be considered location WM selective neurons. We further computed 2-norm of 6 location $\beta$s to evaluate tuning strength of each neuron, thereby summarizing the six coefficients by a single scalar metric.

***Decision selectivity of each neuron***

We conducted linear regression for each neuron to quantify the tuning properties as follows:

$$y = x * \beta + \beta_0$$

For the regression of decision, $y$ represented the neuronal activity for each trial (a column vector with 1 by number of trial), $\beta_0$ was the intercept term, $\beta$ was the regression coefficient to be fitted, and $x$ represented the behavioral label for each trial (0 for offload and 1 for memory).

$$P(\text{seq} = 35 \mid activity)$$
$$= \big(1 - p(location = 1|activity)\big) * \big(1 - p(location = 2|activity)\big)$$
$$* \ p(location = 3|activity) * \big(1 - p(location = 4|activity)\big)$$
$$* \ p(location = 5|activity) * (1 - p(location = 6|activity))$$

Then, we computed *entropy* in a classical way, through distribution of all possible decoded probabilities of sequences, where:

$$Entropy(activity) = -\sum_{seq} P(seq \mid activity) * \log\big(P(seq \mid activity)\big) \in [0,\ Max]$$

Here, we also compute the theoretical upper bound of the *entropy*, by computing entropy of a uniform distribution, denoted as *Max*. Finally, we computed WM strength by inverting and normalizing the entropy from [0, Max] to [0, 1], where:

$$WM\ strength(activity) = \frac{Max - Entropy(activity)}{Max} \in [0,\ 1]$$

WM strength = 0 represented a uniform probability distribution over all possible sequences and WM strength = 1 represented distributions where probability of one specific sequence came to exactly 1.

For analyses utilizing pseudo-population where data from all sessions were combined, we resampled the same number of trials (including memory correct trials and forced-to-test correct trials) of each sequence type across sessions. The resampling trial numbers across sequences were determined in a sequence length-dependent and location-balanced manner. Within each sequence length, resampling was performed to balance the number of trials across different locations. This resampling was repeated 32 times. For each resampling, we trained and tested a new set of decoders from which to recompute WM strength. After all resampling procedures, we averaged the testing results to produce the final pseudo-population-based evaluations of WM (Fig. S3).

**Neural activity-based meta-WM analysis (Related to Figure 3)**

***Meta-WM selectivity of each neuron***

We conducted linear regression for each neuron to quantify its tuning properties regarding putative meta-WM (or how well the monkeys might consider their WM of stimuli locations to be) as follows:

$$y = x * \beta + \beta_0$$

Where $y$ represented the neuronal activity for each sequence (a column vector with 1 by number of sequences), $\beta_0$ was the intercept term, $\beta$ was the regression coefficient to be fitted (a scalar), and $x$ represented the behavioral offloading rate for each sequence (a column vector with 1 by number of sequences).

***Linear regression of trial-level meta-WM***

In this analysis, we utilized baseline meta-WM and (delay) WM strength as predictors to regress the (delay) meta-WM of each trial, where:

$$Meta\_\mathrm{WM} = \beta_1 * baseline\ meta + \beta_2 * WM\ strength + \beta_{12} * baseline\ meta * WM\ strength + \beta_0$$

The regression coefficients were estimated using a resampling approach with replacement. For each resampling iteration, the number of samples matched the total number of trials. We used the resampled trials to conduct linear regression and to calculate the $\beta$ coefficients and regression explained variance. This process was repeated 1,000 times to generate a distribution of $\beta$ coefficients for statistical testing.

**Three task variables relationship analysis (Related to Figure 6)**

***Subspace estimation and principal angles between subspaces***

Based on the previously decoded variables (baseline meta-WM, WM strength, and meta-WM), we performed linear regression to model these variables across trials using neuronal population activity. For example, we regressed meta-WM scores on neuronal population activity. To avoid overfitting, we employed Lasso regularization with 3-fold cross-validation in the linear regression. The beta coefficients obtained from these regressions defined an one-dimensional subspace (axis) for each variable, forming vectors within the neuronal state space. We then computed the angles between these one-dimensional subspaces[63]. As a control, we randomly split the trials into two halves to obtain separate estimations of each subspace and computed the angles between them; this process was repeated 112 times.

***Variance accounted for (VAF) ratio between subspaces***

For given two one-dimensional subspaces, which are vectors of beta coefficients, $\vec{\beta}_a$ and $\vec{\beta}_b$, the VAF ratio[63,64] for subspace pair (a, b) was defined as

$$VAF_{ab} = \frac{Var(\vec{\beta}_{b\_unit} * \vec{\beta}_{b\_unit}^{T} * \vec{\beta}_a)}{Var(\vec{\beta}_a)}$$

Where $\vec{\beta}_{b\_unit}$ is the unit vector of $\vec{\beta}_b$. As a control, we randomly split the trials into two halves to obtain separate estimations of each subspace and computed the VAF ratio between them, and this process was repeated 112 times.

***Time course of AUROC within subspaces***

As previously described, for each variable, we derived a vector of regression coefficients ($\vec{\beta}$) via linear regression to quantify the subspace. Taking the meta-WM subspace as an example, we projected the neuronal population activity matrix (F) at each time bin (33 ms) onto this vector to compute the estimated meta-WM scores (EMS):

$$\mathrm{EMS} = \vec{\beta} * F + \beta_0$$

Here, $\vec{\beta}$ is a 1-by-n vector, where n denotes the number of neurons; $F$ is an n-by-m matrix, with m representing the number of trials; and $\beta_0$ is the intercept term from the linear regression model. This projection, from neuronal activity to the meta-WM subspace, yielded a scalar meta-WM score for each trial at each time bin. The same procedure was employed to estimate scores for the baseline meta and WM strength subspaces. Subsequently, we analyzed the time courses of AUROC to differentiate between memory and offload trials within both the baseline meta and meta-WM subspaces, as well as between low-strength and high-strength trials within the WM strength subspace.

***Spatial organization of selective neurons***

We quantified and compared the spatial distances within and between groups (baseline meta-WM, WM, meta-WM)[65]. We quantified the distance within groups by computing the average distance of each neuron to other neurons within the same group. We quantified the distance between groups by computing the average distance of each neuron to other neurons in different groups.

***Subspace-selective neurons (Related to Fig. S17)***

We randomly split the trials into two halves to obtain two separate estimations for each subspace and the corresponding neuronal coefficients, denoted as $\beta_A$ and $\beta_B$ for a given example neuron. For each of the 112 resampling iterations, we computed the difference between the coefficients ($\beta_{diff} = \beta_A - \beta_B$) and their mean ($\beta_{mean} = (\beta_A + \beta_B)/2$). We then computed the mean difference between the coefficients across all resampling iterations ($\beta_{diff\_mean}$). A neuron was considered to contribute to the subspace in a given resampling if the absolute value of $\beta_{mean}$ exceeded five times the absolute value of $\beta_{diff\_mean}$. Neurons meeting this criterion in more than 99% of the resampling iterations were classified as subspace-selective, indicating a statistically significant and consistent contribution to the subspace.

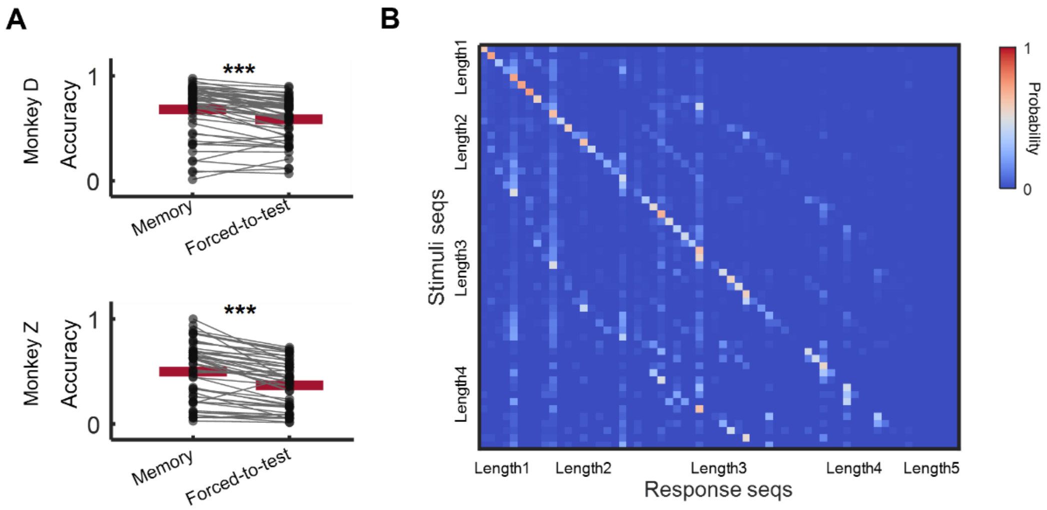


**Fig. S1. Additional behavior performance.**

(**A**) Within-sequence comparison of accuracy of memory trials and forced-to-test trials (two-tailed paired-sample t-test, $p < 0.001$ for both monkeys). Each dot represents one sequence, and red lines represent the mean accuracies across sequences.

(**B**) Stimuli-to-response matrix of behavior data for monkey Z, including both correct trials (diagonal) and error trials (non-diagonal). Each element of the matrix denotes the probability for monkey Z to execute one particular sequence given the presentation of another specific sequence during the sample period.

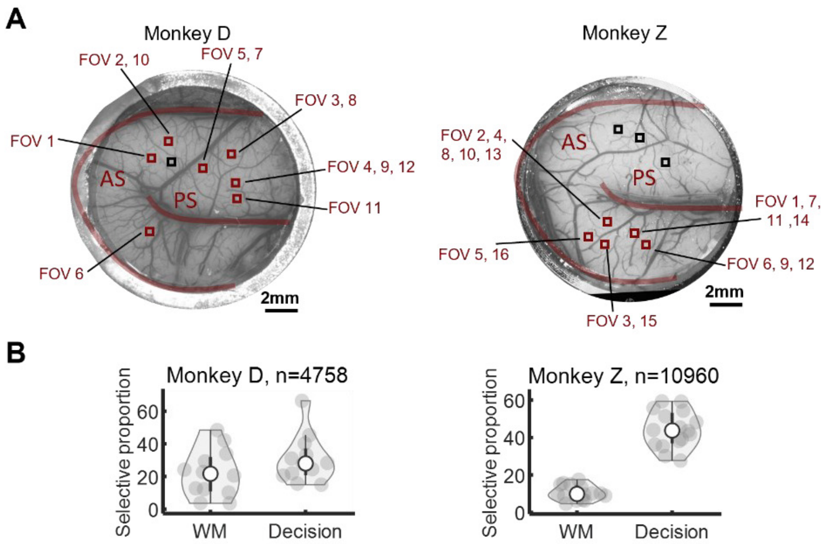


**Fig. S2. FOV distribution and task information-selective neuron proportions.**

(**A**) FOV distribution across imaging windows in both monkeys. Red squares denote FOVs utilized in data analysis, while black squares indicate those excluded from analysis due to poor GCaMP expression.

(**B**) WM- and decision-selective neuron proportions in both monkey D and monkey Z. Each dot represents the proportion of neurons belonging to a selectivity category from one FOV. Overall, 22.3%/10.2% of neurons were WM-selective neurons and 28.6%/46.7% of neurons were decision-selective neurons in monkeys D and Z, respectively, after merging all FOVs.

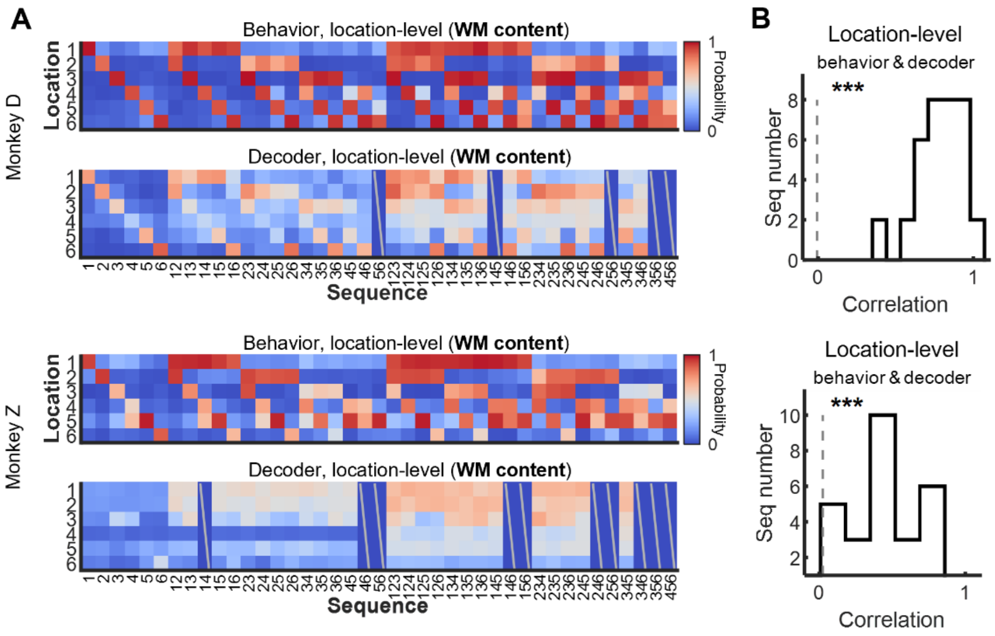


**Fig. S3. Pseudo-population results of neural WM location decoding for both monkeys.** The pseudo-population results were obtained from resampled trials across sessions; see Methods subsection “Neural decoder-based WM strength”. For Monkey D, the pseudo-population results were derived from 10 FOVs encompassing 3,907 neurons. For Monkey Z, the results originated from 10 FOVs consisting of 7,215 neurons. The WM decoder was trained and tested on forced-to-test-correct and memory-correct trials, resampled from each FOV.

(**A**) Illustration of location probability distribution of behavior data (based on accuracy) and the neuron decoder (based on pseudo-population data from 10 FOVs).

(**B**) Histogram of Pearson correlation between location probability distributions of neural population decoder and behavior (A) (***: $p < 0.001$, two-tailed t-test against correlation value by chance).

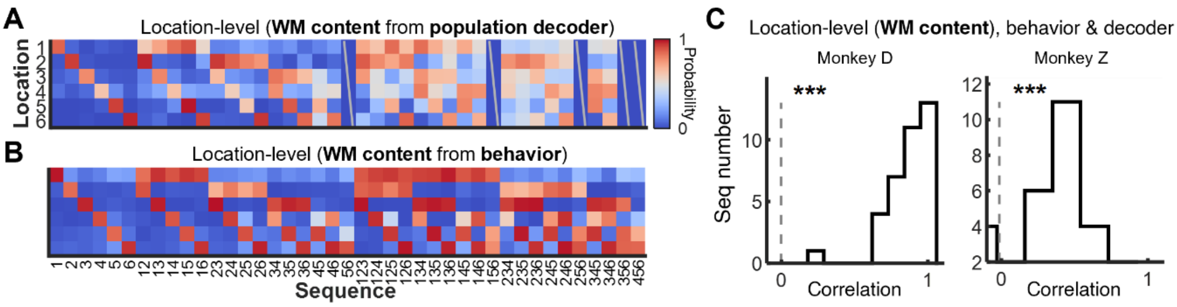


**Fig. S4. Additional analyses of stimulus location decoders-derived WM metrics.**

(**A**) Location probability distributions based on an example neural population decoder (monkey D, FOV 8), representing, for each sequence in the freely-choosing memory condition, the decoded probabilities of each location.

(**B**) Behavioral location probability distributions (monkey D), representing, for each sequence in the forced-to-test condition, the response probabilities of each location based on task performance.

(**C**) Histogram of Pearson correlation between location probability distributions of neural population decoder (A) and behavior (B) (***: $p < 0.001$, two-tailed t-test against correlation value by chance).

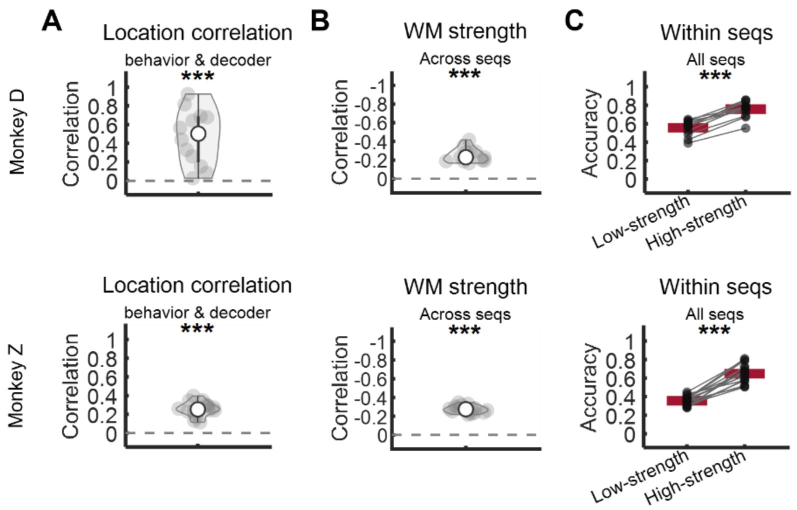


**Fig. S5. WM strength across FOVs and two monkeys.**

Each data point represents an averaged value from one FOV and red lines represent the mean across FOVs.

(**A**) Pearson correlation of the location probability distribution between the population decoder (correct trials) and behavior. This correlation was compared to the chance level using a two-tailed one-sample t-test, with $p < 0.001$ for both monkeys. The chance level was computed as substituting behavior location probability distributions as random values sampled from a uniform probability distribution. In this analysis, we used forced-to-test-correct and memory-correct trials.

(**B**) Pearson correlation between behavior recall variability and sequence-averaged WM strength, based on testing performance from forced-to-test (correct and error) trials. This correlation was compared to the chance level, with a two-tailed one-sample t-test showing $p < 0.001$ for both monkeys.

(**C**) Within-sequence accuracy comparison between low WM strength and high WM strength trials using a two-tailed paired-sample t-test, showing $p < 0.001$ for both monkeys. The threshold dividing low and high WM strength trials was based on the median WM strength of all choice trials. Each dot represents averaged results across sequences from one session. WM strength was assessed from forced-to-test and memory (correct and error) trials.

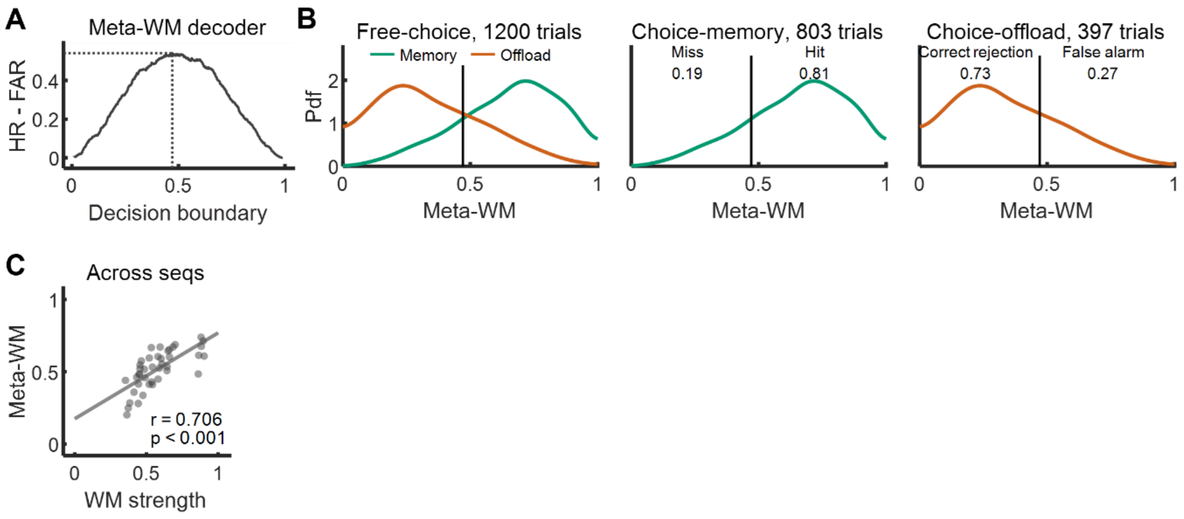


**Fig. S6. Meta-WM decoder performance details, including computation of the optimal decision boundary of the decoder.**

Results from FOV8 of monkey D.

(**A**) The difference between the hit rate (HR) and the false alarm rate (FAR) for each decision boundary of the meta-WM decoder, ranging from 0.01 to 0.99 with a step of 0.001. The decoder reached its optimal value at 0.4710.

(**B**) The trial distribution of meta-WM scores across all memory and offload trials, along with discrimination details of the meta-WM decoder (hit rate, miss rate, correct rejection rate, false alarm rate), given the optimal decision boundary identified in (A).

(**C**) Pearson correlation between meta-WM and WM strength (both decoded from the neuron population) across sequences ($r = 0.705$, $p < 0.001$). Each dot represents a sequence, using all trials of the choice condition.

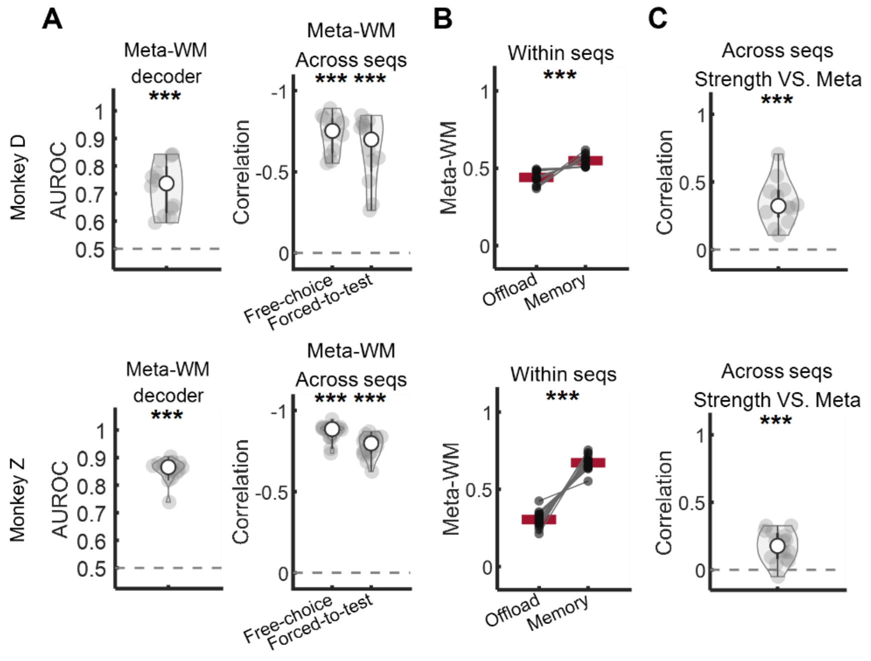


**Fig. S7. Meta-WM decoder performance across FOVs and two monkeys.**

Each data point represents an average value from a single session using choice (memory + offload) trials. ***: $p < 0.001$.

(**A**) Performance of the meta-WM decoder. (Left) Comparison of AUROC with a chance level of 0.5 (using a two-tailed one-sample t-test, $p < 0.001$). (Right) Pearson correlation between the sequence-averaged behavioral offloading rate and the meta-WM scores of choice trials and forced-to-test trials, respectively. The correlation was compared to a chance level of 0 (two-tailed one-sample t-test, $p < 0.001$).

(**B**) Within-sequence comparison of meta-WM scores between offload trials and memory trials (two-tailed paired-sample t-test, $p < 0.001$). Each dot represents the averaged results across sequences from a single session, with red lines indicating the mean. Paired dots originate from the same session.

(**C**) Pearson correlation between meta-WM and WM strength across sequences, along with a comparison of this correlation to a chance level of 0 (two-tailed one-sample t-test, $p < 0.001$). Each dot represents the averaged result across sequences from a single session.

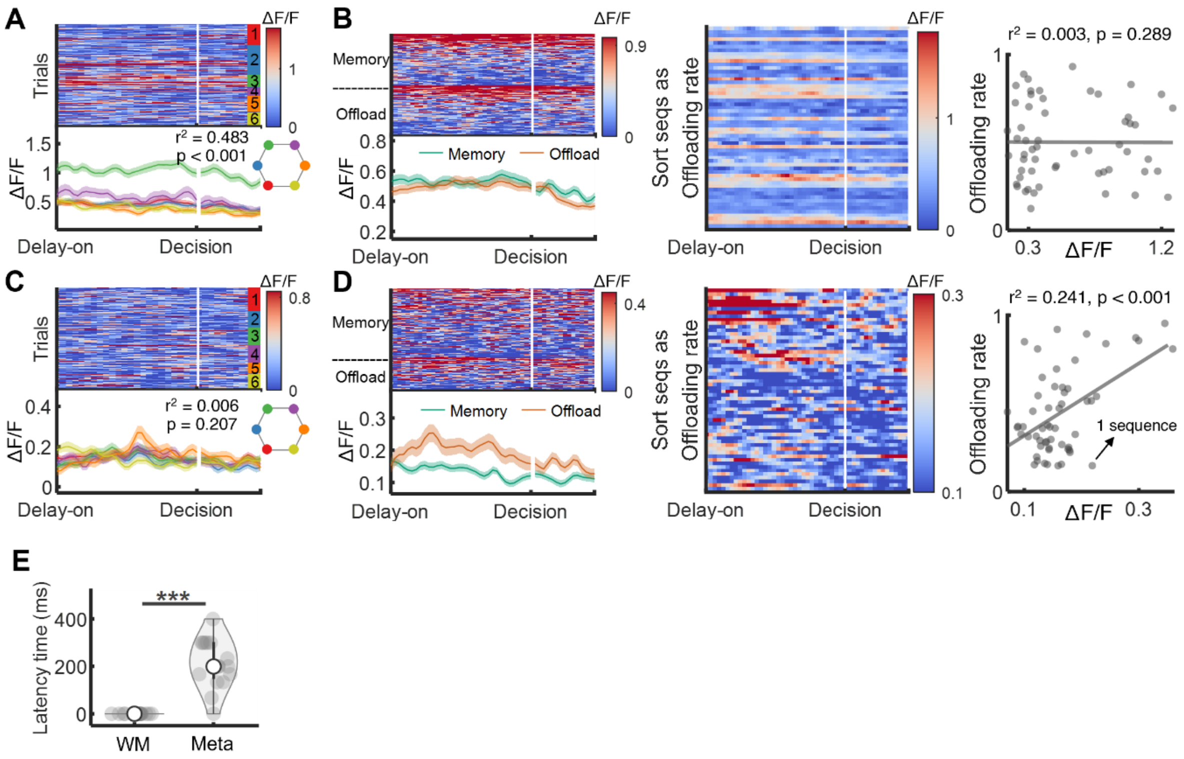


**Fig. S8. Pure selective neurons and latency time between WM strength and meta-WM representations.**

(**A-B**), Characteristics of a pure WM selective neuron. (A) Neuronal activity in response to different stimulus locations. The trials were grouped by whether any of the 6 specific locations was presented in a trial. (B) Neuron activity in memory and offload trials (left, choice trials were sorted as neuron activity descending), as well as in different sequences sorted by offloading rate (middle and right). Shaded areas represent SEMs. Each dot represents one sequence.

(**C-D**), Same as (A-B), but of a pure meta-WM selective neuron.

(**E**) Comparison of latency times between WM and meta-WM as in Fig. 4d. Resampled 16 times to compare latency times. Latency time of WM (after the last target of stimuli): mean = 0 ms, std < 0.001 ms. Latency time of meta-WM (after the last target of stimuli): mean = 212.287 ms, std = 102.367 ms. Each dot represents a result from one resampling, with a total of 16 resampling. Results from FOV8 of monkey D, using choice (memory + offload) trials.

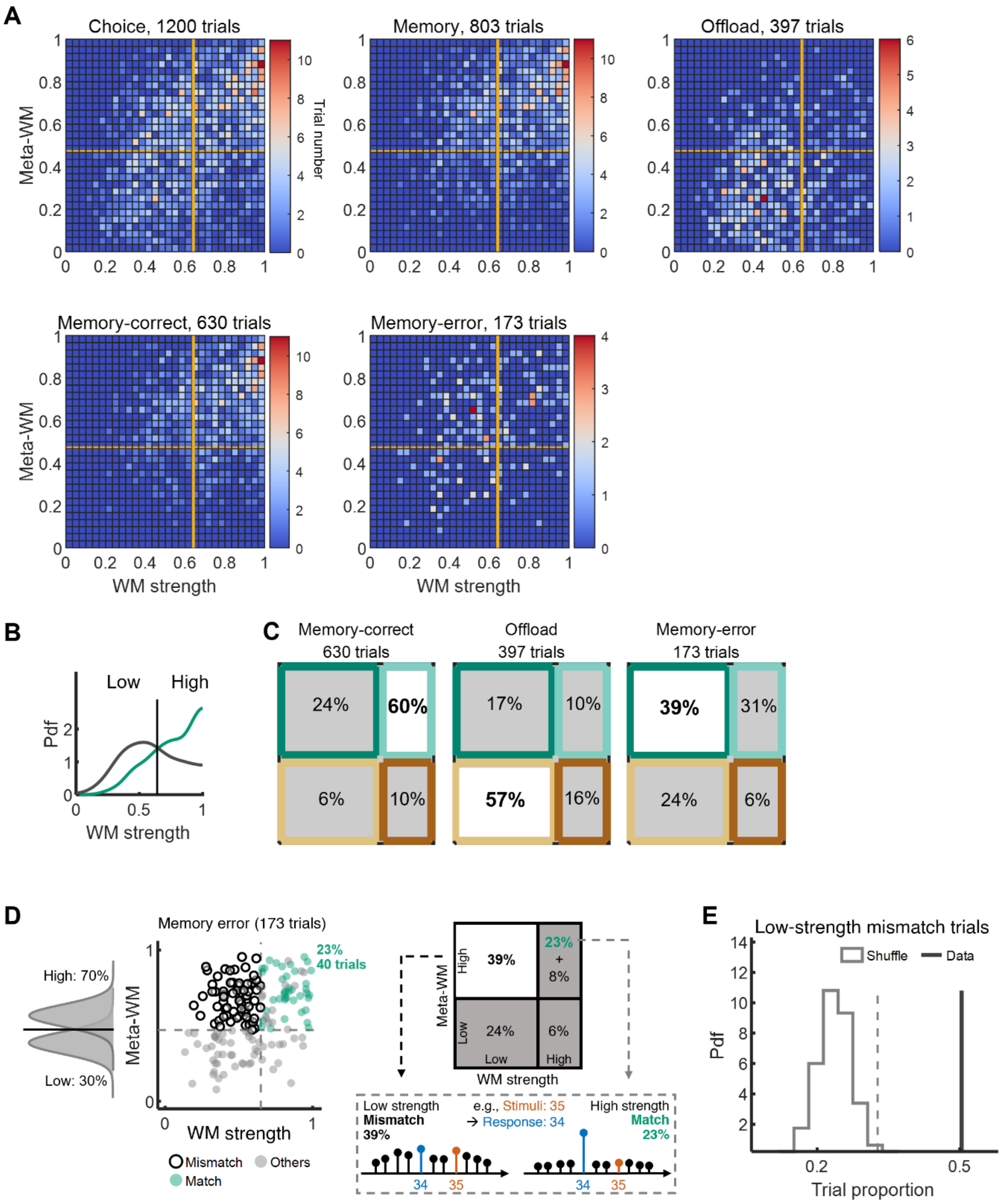


**Fig. S9. More additional results of relationship between WM strength and meta-WM using optimal boundaries.**

Results from an example FOV (FOV8 of monkey D) using choice (memory + offload) trials. The threshold dividing low meta-WM and high meta-WM was determined from the optimal decision boundary of the meta-WM decoder. Similarly, the threshold separating low WM strength from high WM strength was derived from the optimal boundary of WM strength, distinguishing between memory-correct and memory-error trials, akin to the criterion used in meta-WM.

(**A**) Detailed two-dimensional trial distribution heatmaps of all conditions.

(**B**) Probability density functions (Pdf) of the estimated WM strength for all memory-correct trials and all memory-error trials. The threshold (solid line) dividing low WM strength and high WM strength was derived from the optimal boundary.
(**C**) Trial proportions of memory-correct trials, offload trials, and memory-error trials, categorized into four parts using the optimal boundaries.
(**D**) (Top) Memory-error trials are categorized into four groups according to high or low levels of WM strength and meta-WM, determined by median values. Among error trials, 31% exhibited high WM strength and high meta-WM, including 23% match trials (where the decoding probability of response locations was the highest among all sequences) and 8% others. Meanwhile, 39% showed a mismatch with low WM strength and high meta-WM. (Bottom) Two example memory-error trials with sequence of stimuli [3-5] and response [3-4] illustrate match and mismatch. The high-strength match trial had concentrated probabilities responding as sequence [3-4], while the low-strength mismatch trial had widely distributed probabilities across many response sequences.
(**E**) Test of trial proportion validity for low-strength mismatch trials in memory-error trials. Red line represents real data. Gray dashed line represents 99th percentile of a shuffle distribution.

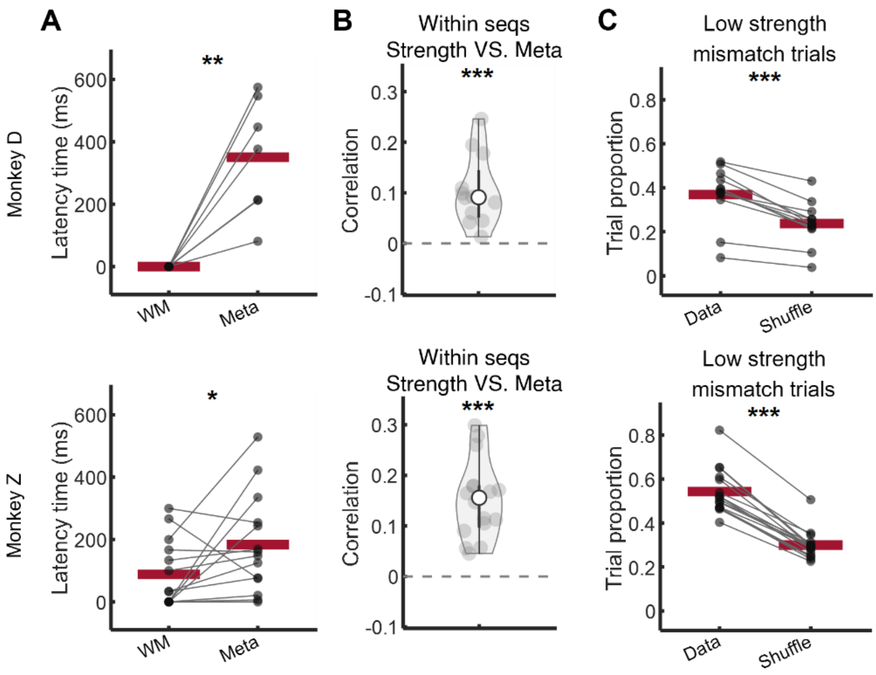


**Fig. S10. Relationship between WM strength and meta-WM across FOVs and two monkeys.**

In multi-FOV results here, each data point represents an averaged value from one FOV and red lines represent the means over FOVs.

(**A**) Comparison of latency time between WM and meta-WM (single-tailed paired-sample t-test, for monkey D, $p < 0.01$; for monkey Z, $p < 0.05$). Time for each session was obtained in the same manner as in Fig. 4d. Latency time of WM (after last target of stimuli): for monkey D, mean = 0 ms, std < 0.001 ms; for monkey Z, mean = 92.764 ms, std = 114.000 ms. Latency time of meta-WM (after last target of stimuli): for monkey D, mean = 350.621 ms, std = 186.821 ms; for monkey Z, mean = 183.150 ms, std = 158.827 ms. Here, only FOVs whose latency time were not outliers were used in this analysis. The outliers were computed as the data points that more than three scaled MAD from the median, which is Matlab "isoutlier" default method. The scaled MAD is defined as c*median(abs(A-median(A))), where c=-1/(sqrt(2)*erfcinv(3/2)) and A is input.

(**B**) Pearson correlation between meta-WM and WM strength within each sequence. Overall mean correlation values are greater than chance for both monkeys (two-tailed one-sample t-test). Each dot represents averaged results across sequences from one session.

(**C**) Comparison of trial proportion of low-strength mismatch trials in memory-error trials with shuffle level (two-tailed paired-sample t-test, for both monkeys). The 'Data' term represents proportions of low-strength mismatch trials in memory-error trials from real data. The 'Shuffle' term represents the results computed from a shuffled resampling (1,000 times of resampling; see methods subsection "Test of trial proportion validity of low-strength mismatch trials in memory-error trials").

*: $p < 0.05$; **: $p < 0.01$; ***: $p < 0.001$.

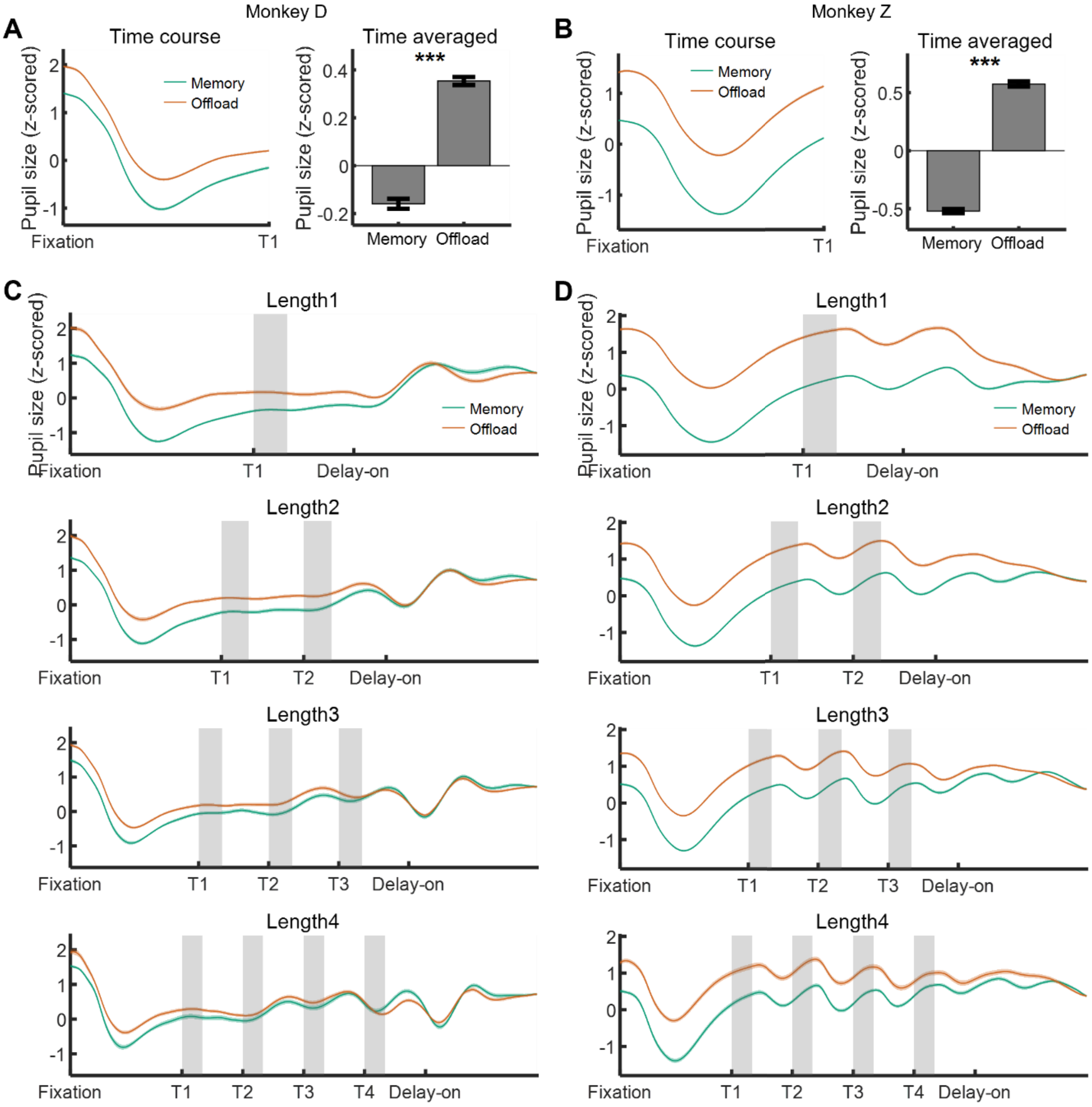


**Fig. S11. Pupil size details.**

(**A-B**) Pupil size during baseline period in monkey D (A) and monkey Z (B), averaged across all sessions and trials. (Left) The time course of pupil size during the baseline period of memory trials and offload trials. (Right) The time-averaged pupil size during the baseline period of memory trials and offload trials. The error bar was computed as SEM across sessions.

(**C-D**) Pupil size in monkey D (C) and monkey Z (D) during the entire fixation time course in trials with different sequence lengths (1 to 4).

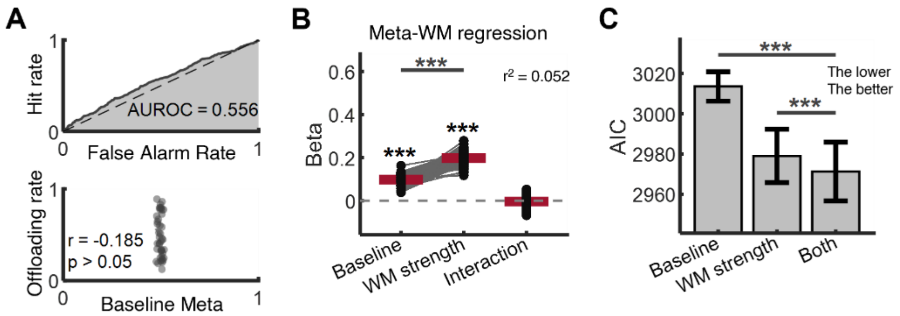


**Fig. S12. Prior belief results from monkey Z.**

(**A**) Performance of the baseline meta-WM decoder. (Top) AUROC. (Bottom) Pearson correlation between offloading rates and sequence-averaged baseline meta-WM. Each dot represents one sequence. Results from FOV 13 of monkey Z using choice (memory + offload) trials.

(**B**) Meta-WM linear regression coefficients of baseline and WM strength. While both baseline meta-WM and WM strength (measured during the delay) positively impacted meta-WM (single-tailed one-sample t-tests), WM strength was a stronger predictor of meta-WM overall (two-tailed paired-sample t-tests between coefficients). Each dot represents a coefficient from a random resampling-based regression model, with red lines representing mean values. Results from FOV 13 of monkey Z, using choice (memory + offload) trials. ***: $p < 0.001$.

(**C**) Comparison of Akaike information criterion (AIC) in three meta-memory linear regression models (***: $p < 0.001$, two-tailed t-test). Models with lower AIC are considered to have higher performance, as AIC incorporates a penalty for the number of parameters. The input factors were baseline meta-WM, WM strength, and both, respectively. Error bars represent STDs. Results from FOV 13 of monkey Z using choice (memory + offload) trials.

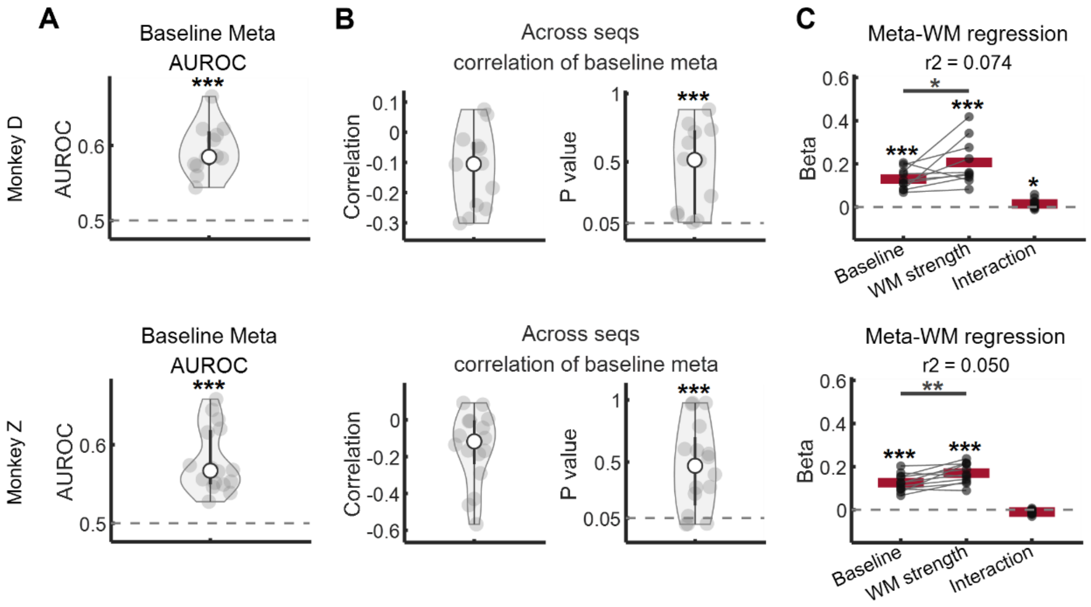


**Fig. S13. Prior belief results across FOVs and two monkeys.**

Multi-FOVs' results. Each data point represents an averaged value from one session and red lines represent the mean. In these analyses, we used choice (memory + offload) trials.

(**A-B**) Performance of the baseline meta-WM decoder. (A) Comparison of AUROC with the chance level of 0.5 (two-tailed one-sample t-test, $p < 0.001$). (B) Correlation between sequence-averaged baseline meta-WM and behavioral offloading rate across sequences. Comparison of p-values of the correlation with 0.05 (two-tailed one-sample t-test, $p < 0.001$). The dashed lines represent the chance level.

(**C**) Meta-WM linear regression coefficients of baseline and WM strength. Comparison of each regression coefficient with the chance level of 0 (single-tailed one-sample t-test, $p < 0.001$ for both the baseline term and WM strength term for both monkeys; $p > 0.05$ for the interaction term for both monkeys). Comparison between the two coefficients (single-tailed paired-sample t-test, for monkey D, $p < 0.05$; for monkey Z, $p < 0.01$). Dashed lines represent chance level. In this analysis, only FOVs with regression coefficients that were not outliers and exceeded the 10th percentile were used. This analysis used all trials.

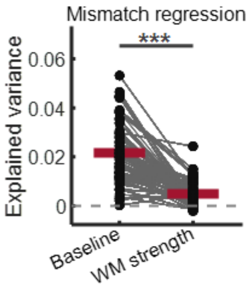


**Fig. S14. Prior belief results of mismatch regression.**

Explained variance ($r^2$) of two linear regression was assessed to distinguish trial types (low-strength mismatch or offload) using baseline or WM strength, including only low-strength mismatch and offload trials. There is significant difference between two $r^2$(two-tailed paired-sample t-test, p < 0.001). Each dot represents a result from one resampling, and red lines represent the mean across 1,000 random samples. Results from FOV 8 of monkey D.

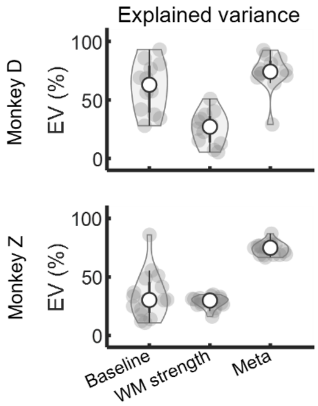


**Fig. S15. Explained variance of decoded variables in each subspace across FOVs and two monkeys.**

Explained variance of decoded variables (scores) across trials in three subspaces. Based on the previously decoded variables (baseline meta, WM strength, and meta-WM), we performed linear regression to model these variables across trials using neuronal population activity. For example, we regressed meta-WM scores on neuronal population activity. To prevent parameter overfitting, we employed Lasso regularization with 3-fold cross-validation in the linear regression. The beta coefficients obtained from these regressions defined a one-dimensional subspace for each variable, forming vectors within the neuronal state space. The explained variance presented here was calculated from these regressions. Each data point represents a result from one session.

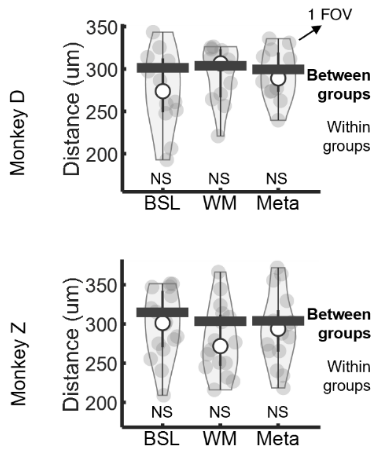


**Fig. S16. Spatial distances analysis across FOVs and two monkeys.**
Comparison of spatial distances within groups and between groups across FOVs (two-tailed t-test). Each gray dot represents the median distance of neurons within a single FOV, where each neuron's distance was calculated as the average spatial distance between that neuron and others within the same group. The black bar represents a similar calculation as the gray dots but for results between different groups, and it shows the median value among FOVs. NS: non-significant. BSL: Baseline.

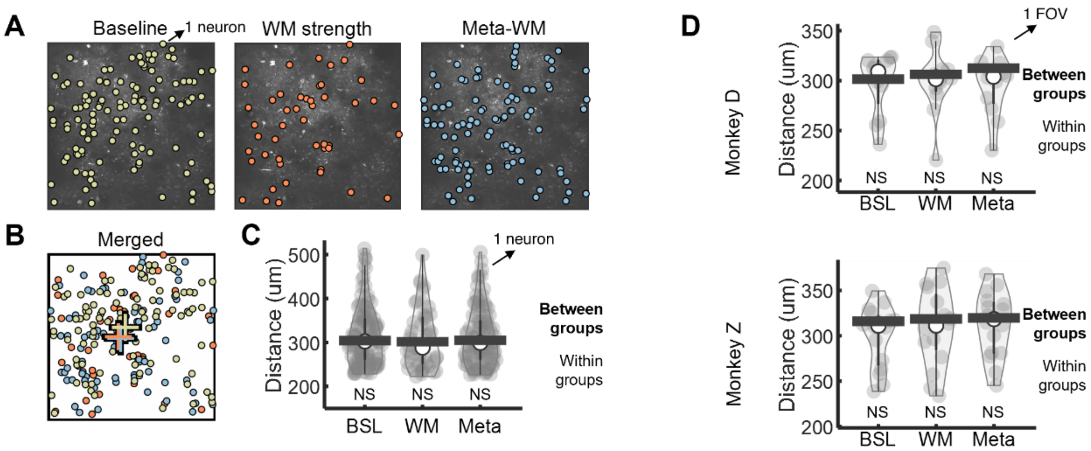


**Fig. S17. Spatial distances analysis of subspace-selective neurons.**

(**A-B**) (A) Spatial organization of subspace-selective neurons (see Methods subsection “Subspace-selective neurons”) for baseline meta, WM strength, and meta-WM (from FOV 4 of monkey D; 113, 48, 100 neurons, respectively). Each dot represents a neuron. In the merged panel (B), crosses represent spatial centroids.

(**C**) Comparison of spatial distances within groups and between groups (two-tailed t-test; FOV 4 of monkey D). Each gray dot represents the average spatial distance between one neuron and others within the same group in (A). The black bar represents a calculation similar to that of the gray dots, but specifically for results between different groups, and it shows the median value among neurons. NS: non-significant.

(**D**) Comparison of spatial distances within groups and between groups across FOVs (two-tailed t-test). Each gray dot represents the median distance of neurons within a single FOV, where each neuron's distance was calculated as the average spatial distance between that neuron and others within the same group. The black bar represents a similar calculation as the gray dots but for results between different groups, and it shows the median value among FOVs. NS: non-significant. BSL: Baseline.

Monkey D, neuron number distribution

| FOV | Total | WM (Location) | Decision | Meta-WM | Prior (Baseline) |
|---|---|---|---|---|---|
| 1 | 322 | 12 | 89 | 33 | 35 |
| 2 | 180 | 22 | 37 | 46 | 54 |
| 3 | 220 | 28 | 146 | 66 | 110 |
| 4 | 410 | 120 | 186 | 108 | 66 |
| 5 | 529 | 54 | 150 | 45 | 26 |
| 6 | 719 | 26 | 223 | 70 | 230 |
| 7 | 524 | 108 | 115 | 100 | 58 |
| 8 | 702 | 340 | 172 | 148 | 249 |
| 9 | 607 | 206 | 95 | 98 | 147 |
| 10 | 223 | 54 | 74 | 22 | 14 |
| 11 | 220 | 50 | 33 | 30 | 52 |
| 12 | 102 | 43 | 41 | 20 | 30 |
| All FOVs | 4758 | 1063 | 1361 | 786 | 1071 |

Monkey Z, neuron number distribution

| FOV | Total | WM (Location) | Decision | Meta-WM | Prior (Baseline) |
|---|---|---|---|---|---|
| 1 | 374 | 36 | 132 | 35 | 39 |
| 2 | 415 | 65 | 125 | 28 | 18 |
| 3 | 347 | 16 | 96 | 19 | 66 |
| 4 | 388 | 58 | 187 | 56 | 100 |
| 5 | 148 | 15 | 58 | 18 | 29 |
| 6 | 916 | 80 | 332 | 43 | 66 |
| 7 | 556 | 35 | 302 | 62 | 153 |
| 8 | 639 | 96 | 278 | 61 | 64 |
| 9 | 991 | 100 | 436 | 244 | 226 |
| 10 | 640 | 112 | 244 | 39 | 69 |
| 11 | 672 | 60 | 305 | 57 | 158 |
| 12 | 917 | 122 | 383 | 236 | 89 |
| 13 | 892 | 90 | 458 | 104 | 62 |
| 14 | 1133 | 77 | 671 | 159 | 527 |
| 15 | 946 | 90 | 527 | 193 | 183 |
| 16 | 986 | 63 | 585 | 159 | 231 |
| All FOVs | 10960 | 1115 | 5119 | 1513 | 2080 |

**Table S1. Distributions of neurons according to their task involvement across FOVs for both monkeys.**
